# The rate and potential relevance of new mutations in a colonizing plant lineage

**DOI:** 10.1101/050203

**Authors:** Moises Exposito-Alonso, Claude Becker, Verena J. Schuenemann, Ella Reiter, Claudia Setzer, Radka Slovak, Benjamin Brachi, Jörg Hagmann, Dominik G. Grimm, Jiahui Chen, Wolfgang Busch, Joy Bergelson, Rob W. Ness, Johannes Krause, Hernán A. Burbano, Detlef Weigel

## Abstract

By following the evolution of populations that are initially genetically homogeneous, much can be learned about core biological principles. For example, it allows for detailed studies of the rate of emergence of *de novo* mutations and their change in frequency due to drift and selection. Unfortunately, in multicellular organisms with generation times of months or years, it is difficult to set up and carry out such experiments over many generations. An alternative is provided by “natural evolution experiments” that started from colonizations or invasions of new habitats by selfing lineages. With limited or missing gene flow from other lineages, new mutations and their effects can be easily detected. North America has been colonized in historic times by the plant *Arabidopsis thaliana*, and although multiple intercrossing lineages are found today, many of the individuals belong to a single lineage, HPG1. To determine in this lineage the rate of substitutions – the subset of mutations that survived natural selection and drift –, we have sequenced genomes from plants collected between 1863 and 2006. We identified 73 modern and 27 herbarium specimens that belonged to HPG1. Using the estimated substitution rate, we infer that the last common HPG1 ancestor lived in the early 17^th^ century, when it was most likely introduced by chance from Europe. Mutations in coding regions are depleted in frequency compared to those in other portions of the genome, consistent with purifying selection. Nevertheless, a handful of mutations is found at high frequency in present-day populations. We link these to detectable phenotypic variance in traits of known ecological importance, life history and growth, which could reflect their adaptive value. Our work showcases how, by applying genomics methods to a combination of modern and historic samples from colonizing lineages, we can directly study new mutations and their potential evolutionary relevance.

## SUMMARY

A consequence of an increasingly interconnected world is the spread of species outside their native range — a phenomenon with potentially dramatic impacts on ecosystem services. Using population genomics, we can robustly infer dynamics of colonization and successful population establishment. We have compared hundred genomes of a single *Arabidopsis thaliana* lineage in North America, including genomes of contemporary individuals as well as 19^th^ century herbarium specimens. These differ by an average of about 200 mutations, and calculation of the nuclear evolutionary rate enabled the dating of the initial colonization event to about 400 years ago. We also found mutations associated with differences in traits among modern individuals, suggesting a role of new mutations in recent adaptive evolution.

## INTRODUCTION

Colonizing or invasive populations sampled through time (1, 2) constitute “natural experiments” where it is possible to study evolutionary processes in action (3). Colonizations, which are dramatically increasing in number (4, 5), sometimes are characterized by strong bottlenecks and genetic isolation (6, 7), and thus greatly facilitate the observation of new mutations and potentially their effects under natural population dynamics and selection (8). Colonizations thus offer a complementary approach to other studies of new mutations, which often minimize natural selection, for example in laboratory mutation accumulation experiments (9) and parent-offspring comparisons (10). The study of colonizations is also complementary to the investigation of genetic divergence over long time scales, e.g., between distant species (11), where the results are largely independent of short-term demographic fluctuations. There is broad interest in understanding how genetic diversity is generated (12), (12)and how new mutations can provide a path for rapid adaptive evolution (13–15). Additionally, accurate evolutionary rates permit dating historic population splits, which is fundamental to the study of population history (16).

The analysis of colonizing populations can also contribute to resolving the “genetic paradox of invasion” (17). This paradox comes from the observation that colonizing populations can be surprisingly successful and spread very widely even when strongly bottlenecked, suggesting some level of adaptation to new environments that goes beyond the exploitation of unoccupied ecological niches (17). Much of the work in plant ecology and evolution has focused on evidence that populations can rapidly adapt from standing variation (18). In invasive lineages, initial standing variation may originate from incomplete bottlenecks, multiple introductions, or admixture with local relatives (19). Much less work has been done with respect to the role of *de novo* mutations as a solution to the genetic paradox of invasion, although this has been proposed as an alternative explanation for rapid adaptation by colonizing lineages (3, 17, 20).

The self-fertilizing plant *Arabidopsis thaliana* is native to Africa and Eurasia (21,22) but has recently colonized N. America, where it likely experienced a strong founder effect (23). At nearly half of N. American sites sampled during the 1990s and early 2000s, more than 80% of plants belong to a single haplogroup, HPG1, as inferred from genotyping with 149 intermediate-frequency markers evenly spread throughout the genome (23). The HPG1 lineage has been reported from many sites along the East Coast and in the Midwest as well as at a few sites in the West (23) (Figure 1, Table S1). The great ubiquity of HPG1 in comparison to any other haplogroup could be due to either some adaptive advantage, or, more parsimoniously, be the result of HPG1 being derived from one of the first arrivals of *A. thaliana* in the continent.

**Figure 1.**
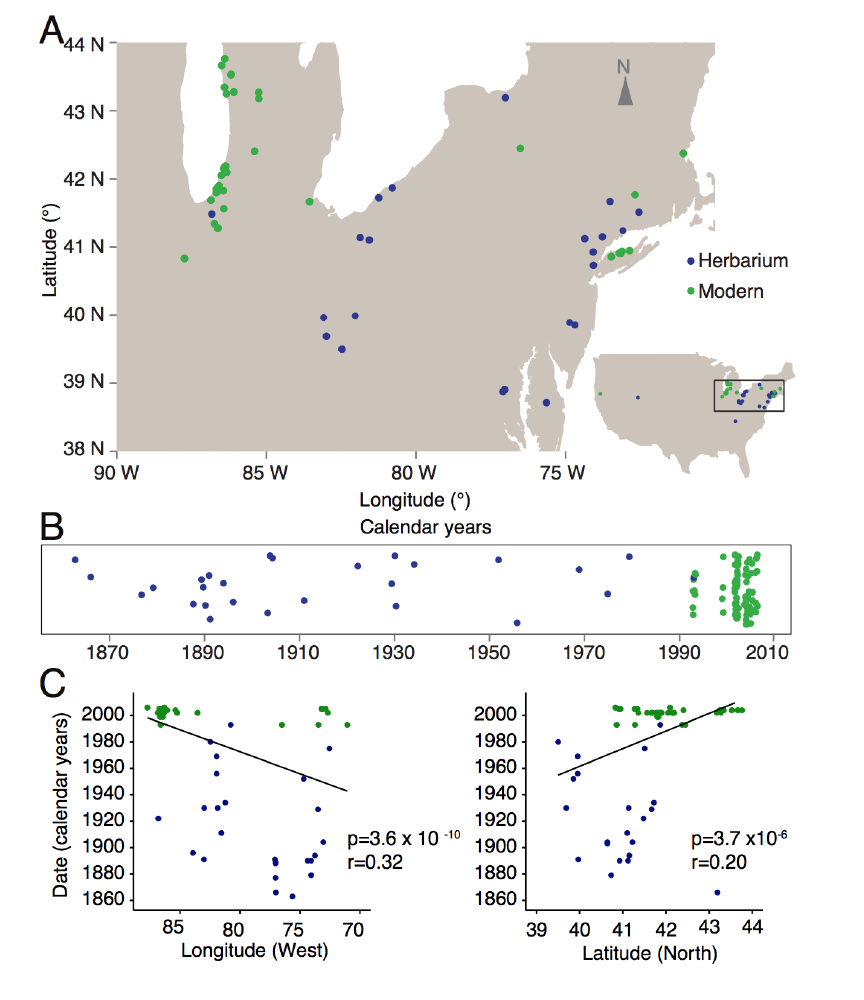
Geographic location and temporal distribution of HPG1 samples. **(A)** Sampling locations of herbarium (blue) and modern individuals (green). **(B)** Temporal distribution of samples (random vertical jitter for visualization purposes). **(C)** Linear regression of latitude and longitude as a function of collection year (p-value of the slope and Pearson correlation coefficient are indicated).

Here, we focus on 100 HPG1 individuals that do not show any evidence of outcrossing with other lineages. We combine genomes from herbarium specimens and live individuals, collectively covering the time span from 1863 to 2006, to infer mutation rates, to date the birth of the HPG1 lineage, and to investigate the evolutionary forces that shape genetic diversity. Our analyses of this lineage serves as a model for future studies of similar colonizing or otherwise recently bottlenecked plant populations, in order to better understand how diversity is generated and to which extent it contributes to adaptation in nature.

## RESULTS AND DISCUSSION

### Historic and modern genomes

In a self-fertilizing species, a single individual can give rise to an entire lineage of millions of offspring, which then diversify through new mutations and eventually intra-lineage recombination. If self-fertilization is much more common than outcrossing, the founder is likely to have been homozygous throughout almost the entire genome. Because it is so wide spread, HPG1 presents an opportunity to sample many natural populations that have been potentially derived from a common, very recent ancestor with such characteristics. In the best possible case, this would allow for new mutations to be directly observed through time. To test these assumptions and to better understand the evolution of HPG1, we sequenced two different groups of plants. The first group were live descendants of 87 plants that had been collected between 1993 and 2006 (Fig. 1; Table S1), and which had been identified as likely members of the HPG1 lineage with 149 genome-wide markers spaced at roughly 1-Mb-intervals (23). We aimed for broad geographic representation, with at least two accessions per collection site, where available. The second group comprised 35 herbarium specimens, collected between 1863 and 1993, for which we had no a priori information whether they may or may not belong to the HPG1 lineage, but which were selected from the herbarium records to cover the full historical geographic range and overlap with modern samples when possible (Fig. 1).

The DNA from the herbarium specimens showed biochemical features typical of ancient DNA (aDNA) from plants, which we have previously described in detail (24). Such DNA damage included a median fragment length of 60 bp, an excess of C-to-T substitutions of about 2.5% at the first base of sequencing reads and a 1.5 to 1.8 fold enrichment of purines at DNA breakpoints (Fig. S1, Supplementary Text 2). To remove aDNA associated damage and produce high-quality genomes, chemically-repaired libraries (see Methods) were later sequenced. These reads were mapped against an HPG1 pseudo-reference genome (25), focusing on single nucleotide polymorphisms (SNPs) because the short sequence reads of herbarium samples preclude accurate calling of structural variants. Genome sequences were of high quality, with herbarium samples covering 96.8–107.2 Mb of the 119 Mb reference, and modern samples covering 108.0–108.3 Mb (Table S1).

### Genetic diversity of HPG1 and delineation from other lineages

We visualized the relationships between the sequenced historic and modern plants building a neighbor joining tree of all 123 samples and confirmed that the majority fell within a almost-identical clade, the HPG1 (Fig. 2A) (23). Because any degree of introgression from other non-HPG1 lineages would confound the discovery of new mutations downstream, we removed all divergent samples and built a neighbour joining tree (n = 103 samples), which revealed that the HPG1 samples were very similar to each other, with very little within-population structure (Fig. 2B). A parsimony network was used to detect recombinant genomes within this HPG1 clade (Fig. 2C), which led us to remove three potential intra-lineage recombinants. Repeating the parsimony network cleared all previously inferred reticulations due to recombinations (Fig. 2D). After such stringent filtering, we kept 27 of the 35 herbarium samples, and 73 of the 87 modern samples (Table S1). These constitute a set of non-admixed, non-recombined and quasi-identical HPG1 individuals.

**Figure 2.**
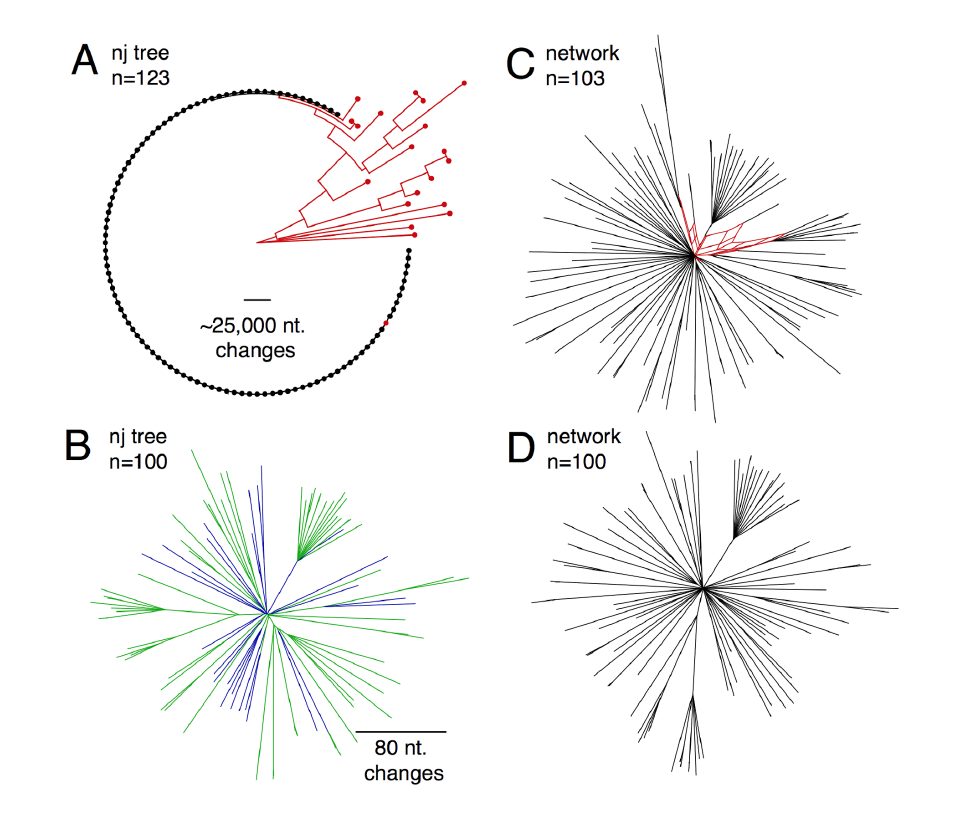
Relationship among herbarium and modern samples. **(A)** Neighbor joining tree with all 123 samples (dots) and rooted with the most distant sample. The black clade of almost-identical samples is the HPG1 lineage. Scale line shows the equivalent branch length of over 25,000 nucleotide changes. **(B)** Neighbor joining tree only with the HPG1 black clade from (A). Colors represent herbarium (blue) and modern individuals (green). Scale line shows the equivalent branch length of 80 nucleotide changes. Note that no outgroup was included. **(C, D)** Network of samples using the parsimony splits algorithm, before **(C)** and after **(D)** removing three intra-HPG1 recombinants (in red). Note that the network algorithm returns in (D) a network devoid of any reticulation, which indicates absence of intra-haplogroup recombination.

Pairs of HPG1 herbarium genomes differed by 28-207 SNPs genome-wide, pairs of HPG1 modern genomes by 2-259 SNPs, and pairs of historic-modern HPG1 genomes by 56-244 SNPs. That is, whole-genome identity was at least 99.9997% in any of pair-wise comparison. Of the approximately five to six thousand segregating SNPs in the HPG1 population, the vast majority, about 95% (Supplementary Text 3), have not been reported outside of this lineage (21). Importantly, the density of SNPs along the genome was low and evenly distributed (typically fewer than 20 SNPs / 100 kb) with no peaks of much higher frequency, which makes us confident that chunks of introgressions from other lineages do not exist in this putatively pure HPG1 set (Fig. 4). As a reminder, random pairs of *A. thaliana* accessions from the native range or pairs of non-HPG1 typically differ by about 500 SNPs / 100 kb (21) (see scale in Fig. 2A).

There were no SNPs in mitochondrial nor chloroplast genomes, which already suggested a recent common origin, and genome-wide nuclear diversity (π = 0.000002, θ_W_ = 0.00001, with 5,013 full informative segregating sites) was two orders of magnitude lower than in the native range of the species (θ_W_ = 0.007) (21) (Table S1) (Supplementary Text 6). The population recombination parameter was also four orders of magnitude lower (4*N*_e_*r* = ϱ = 3.0x10^−6^ cM bp^−1^) than in the native range (ϱ = 7.5x10^−2^cM bp^−1^) (26) (Supplementary Text 6). While recombination occurs in every generation, regardless of self-fertilization or outcrossing, it is only observable after outcrossing between genetically non-identical individuals, and this is what the population recombination parameter reports. We must stress that because *A. thaliana* can outcross at rates of several percent per generation (23,27), but because the HPG1 population is genetically so homogeneous, we are mostly “blind” to the consequences of outcrossing in this special case. The lack of “observable recombination” in the genome is important, as it allows for the use of straightforward phylogenetic methods to calculate a mutation rate. The enrichment of low frequency variants in the site frequency spectrum (Tajima’s *D* =-2.84; species mean =-2.04, (21)) and low levels of polymorphism are consistent with a recent bottleneck followed by population expansion (Fig. 3). The obvious explanation is that the strong bottleneck corresponds to a colonization founder event, likely by very few closely related individuals, or perhaps only a single plant.

**Figure 3.**
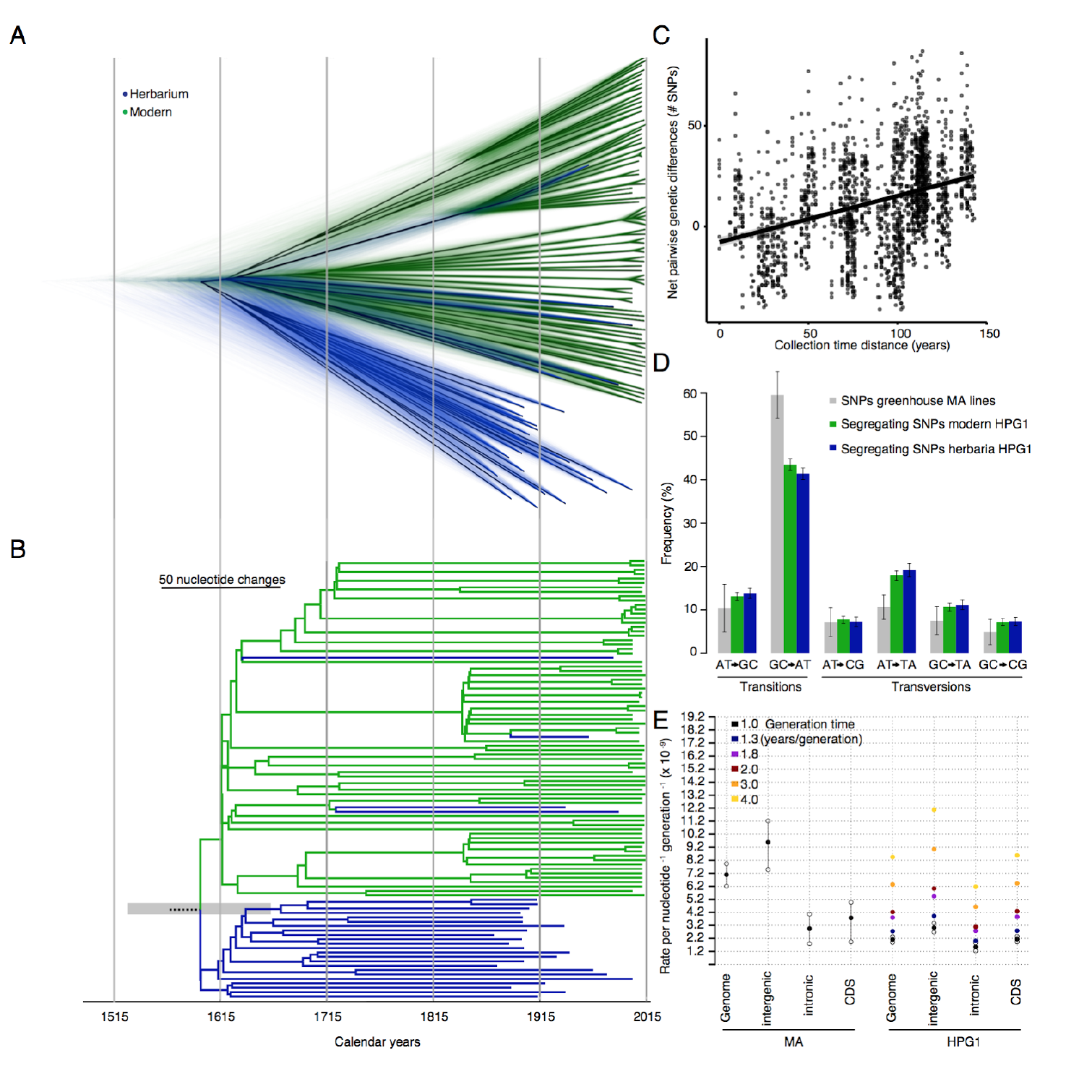
Substitution rates. **(A)** Bayesian phylogenetic analyses employing tip-calibration. A total of 10,000 trees were superimposed as transparent lines, and the most common topology was plotted solidly. Tree branches were calibrated with their corresponding collection dates. **(B)** Maximum Clade Credibility (MCC) tree summarizing the trees in (A). Note the scale line shows the equivalent branch length of 50 nucleotide changes. The grey transparent bar indicates the 95% Highest Posterior Probability of the root date. **(C)** Regression between pairwise net genetic and time distances. The slope of the linear regression line corresponds to the genome substitution rate per year. **(D)** Substitution spectra in HPG1 samples, compared to greenhouse-grown mutation accumulation (MA) lines. **(E)** Comparison of genome-wide, intergenic, intronic, and genic substitution rates in HPG1 and mutation rates in greenhouse-grown MA lines. Substitution rates for HPG1 were re-scaled to a per generation basis assuming different generation times. Confidence intervals in HPG1 substitution rates were obtained from 95% confidence intervals of the slope from 1,000 bootstraps (Table S4 for actual values).

Altogether these patterns indicate that the collection of HPG1 plants we investigated constitute a quasi-clonal and quasi-identical set of individual genomes, mostly devoid of observable recombination and population structure, and thus eminently suited for the study of naturally arising *de novo* mutations.

### The genome-wide substitution rate

It is important to distinguish between the *mutation rate*, which is the rate at which genomes change due to DNA damage, faulty repair, gene conversion and replication errors, and *substitution rate*, which is the rate at which mutations survive and accumulate under the influence of demographic processes and natural selection (28,29). Under neutral evolution, mutation and substitution rates should be equal (29). The simple evolutionary history of the HPG1 population enables direct estimates of substitution rates, and the comparison of theses between different genome annotations, as well as with mutation rates from controlled conditions experiments, could reveal the role played by both demographic and selective forces.

To estimate the substitution rate in the HPG1 lineage, we used distance-and phylogeny-based methods that take advantage of the known collection dates (Supplementary Text 7). The distance method is independent of recombination and has been previously applied to viruses (30) and humans (31). The substitution rate is calculated from correlation between differences in collection time in historic-modern sample pairs, and the number of nucleotide differences between those pairs relative to a reference (Fig. 3C), scaled to the size of the genome accessible to Illumina sequencing. This method resulted in an estimated rate of 2.11x10^−9^substitutions site^−1^year^−1^ (95% bootstrap Confidence Interval [CI]: 1.88–2.33x10^−9^) using rigorous SNP calling quality thresholds. Relaxing the thresholds for base calling and minimum genotyped rate affects both the number of called SNPs and the length of the interrogated reference sequence (32). These largely cancelled each other out, and the adjusted estimates were relatively stable, between 2.1–3.2x10-9 substitutions site ^−1^year^−1^ (Table S3, Supplementary Text 3).

The second method, a Bayesian phylogenetic approach, uses the collection years for tip-calibration and assumes a relaxed molecular clock. It summarizes thousands of plausible coalescent trees, and it has been extensively used to calculate evolutionary rates in various organisms (33–35). This method yielded a substitution rate of 4.0x10^−9^, with confidence ranges overlapping the above estimates (95% Highest Posterior Probability Density [HPPD]: 3.2–4.7x10^−9^).

Based on the similar results obtained with two very different methods, we can confidently say that the substitution rate in the wild populations of HPG1 is between 2 and 5 x10^−9^ site^−1^year ^−1^.

To date the colonization of N. America by HPG1 *A. thaliana* and to improve the description of intra-HPG1 relationships compared to that from a NJ tree, we further used a Bayesian phylogeny. At first sight, the 73 modern samples appeared separated from the herbarium samples (Fig. 3B), but the superimposition of thousands of possible trees showed that the apparent separation of samples was less clear near the root (Fig. 3A). Long terminal branches reflected that the majority of the variants are singletons, typical of populations that expand after bottlenecks.

The mean estimate of the last common HPG1 ancestor, the average tree root, was the year 1597 (HPPD 95%: 1519–1660) (Fig. 3A, B), and an alternative non-phylogenetic method gave a similar estimate, 1625. Both estimates are older than a previously suggested date in the 19^th^ century, using a laboratory mutation rate estimate and having no information from herbarium samples (25). Because HPG1 appears to have been the most abundant lineage in N. America since the 1860s, we believe it could have been one of the first, if not the first colonizer that could establish itself in N. America. If that is true, the time of coalescence of the HPG1 diversity could be close to the time of HPG1 introduction to N. America. During the colonial period, many European immigrants settled on the East coast, consistent with N. American *A. thaliana* lineages being genetically closest to British and coastal West European populations (21). Coincidently, the oldest herbarium samples (12 out of the 27) were HPG1 and came from the East Coast, and we found a significant correlation between collection date and both latitude and longitude (Fig. 1C). This could indicate that after the colonization they moved from the East Coast to the Midwest–the other main area of the distribution that experienced an agricultural expansion in the 19^th^ century (36). Still, these conclusions need to be treated with caution, since regardless of the robustness of the results and our attempts to sample evenly from available collections, there could be unknown biases in the 19^th^ century herbaria.

### Mutation spectra across genome annotations

Although for dating divergence events a substitution rate expressed by years is ideal, in order to compare substitution and mutation rates, both need to be expressed per generation. While *A. thaliana* is an annual plant, seed bank dynamics generate a delay of average generation time at the population scale. A comprehensive study of multiple *A. thaliana* populations in Scandinavia found that dormant seeds could wait for longer than a year in the seed bank, generating overlapping generations and an delayed average generation time of 1.3 years (37) with a notable variance across populations. Multiplication by the mean generation time led to an adjusted rate of 2.7x10^−9^ substitutions site ^−1^generation^−1^ (95% CI 2.4-3.0x10^−9^) (Fig. 3E). To be able to compare this rate with a reference, we also re-sequenced mutation accumulation (MA) lines in the Col-0 reference background grown under controlled conditions in the greenhouse that had been analyzed before with less advanced short read sequencing technology (38). From the new re-sequencing data, we obtained an updated rate of 7.1x10^−9^ mutations site^−1^ generation^−1^ (95% CI 6.3–7.9x10^−9^) (Tables S2, S3, Supplementary Text 4 and 7). This is two- to three-fold higher than the per-generation estimate in the wild, but within the same order of magnitude. The same holds for rates in different genome annotations, i.e. genic, intronic and intergenic regions, but the confidence intervals overlapped in many cases (Table S3).

Differences in per-generation rates between laboratory and wild populations could stem from both methodological as well as biological causes. For instance, if the true average generation time was actually over 3 years / generation, the differences would cancel out (Fig. 3E). Limitations in mapping structural variation in non-reference samples could lower the substitution rate, what explains that we calculated an atypically low substitution rate in regions with transposable elements (see Supplementary Text 7.2.1). Environmentally-driven effects that are not yet well understood, such as variable methylation status of cytosines, which account for much of the variation in local substitution rates (39), could increase or decrease the rate (see Supplementary Text 7.2.3, Fig. S4).

An alternative evolutionary explanation to the aforementioned laboratory and wild populations’ rates differences is that purifying selection in the wild would slow down the accumulation of mutations by removing deleterious mutations (Fig. 3E). This has been observed before and is one of the accepted causes of the discrepancy between the so called long-and short-term substitution rates in a range of organisms (40).

In order to provide evidence for negative purifying selection acting in the wild, we performed three types of analyses involving comparisons across genomic annotations within the HPG1 dataset. Firstly, by calculating contingency tables and computing a Fisher’s exact test, we compared the deviation of expected and observed SNPs between coding regions (more likely under purifying selection), with intergenic regions, intronic regions, and all non-coding regions of genome. All three pairwise comparisons showed a depletion of coding SNPs and an enrichment of intergenic, intronic and non-coding SNPs (odds ratio>2, p<10^−16^). An obvious explanation is that in genome annotations where a mutation is more likely to be deleterious, i.e. coding regions, the number of observed variants should be lower due to selection having removed them from the population before we could sequence them.

Secondly, we studied the Site Frequency Spectrum (SFS) of genetic variants. The rationale was that because purifying natural selection is more efficient at removing intermediate-frequency variants, variants that tend to be deleterious or slightly deleterious should be found at lower frequency than those that only suffer neutral drift (41). We built contingency tables of coding, intergenic, intronic and non-coding variants segregating above and and below the conventional frequency cutoff of 5% to separate low-and intermediate-frequency variants (42). We found that SNPs in coding regions were more likely to be at low frequency than those in intergenic (odds ratio=2.34, p=3.09x10^−11^), intronic (odds ratio=1.48, p=0.02), and all non-coding regions (odds ratio=2.05, p=1.29x10^−8^). We carried out the same analysis using nonsynonymous and synonymous SNPs, which are easily interpretable in terms of the selection regimes under which they evolve. We did not find an enrichment (p=0.67), perhaps a consequence of the small number of such mutations (Table S3).

Thirdly, to verify that the full frequency spectrum of coding SNPs was shifted to lower frequencies (i.e. the results were not dependent on the arbitrary 5% frequency cutoff), we used the nonparametric Kolmogorov-Smirnov test for two samples. We found that the cumulative distribution of the site frequency spectrum (CD_SFS_) of coding regions is above (i.e., the frequency distribution is overall skewed to lower values) both the intergenic CD_SFS_ (p=3.25x10^−6^) and the non-coding regions CD_SFS_ (p=0.001), but not the intronic CD_SFS_(p=0.60) (Fig. S5). As in our previous analysis, the comparison between the nonsynonymous and synonymous CD_SFS_ yielded, likely for similar reasons, no differences (p=0.53).

All in all, these results support that purifying selection is a force shaping to some degree the diversity across the HPG1 genome and might therefore as well contribute to the differences between HPG1 and MA rates.

### Potentially advantageous *de novo* mutations

Finally, having discovered over 5,000 *de novo* mutations in the HPG1 lineage, we wondered whether there is any evidence for an adaptive role of these *de novo* mutations in the colonization of N. America by HPG1. We noted that some new mutations had risen to intermediate or even high frequencies in the HPG1 samples. This might have been the consequence of drift from stochastic demographic processes, or it could have been caused by positive natural selection. To find direct evidence for the latter, we grew the modern accessions in a common garden and studied phenotypes of known importance in ecology of invasions (43), namely flowering time and root traits (see Supplementary Text 8). Using linear mixed models, we calculated the proportion of variance explained (also called narrow sense heritability, h^2^) with a kinship matrix of all SNPs that had become common (>5%, n=391). We found significant heritable variation for multiple traits including the growth rate in length (h^2^ = 0.64) and the average root gravitropic direction (h^2^ = 0.54). As in our study mutations are the main source of genetic variants, these mutations — or mutations linked to them — should be responsible for significant quantitative variation in several traits (Table S4, Supplementary Text 10). The existence of mutation-driven phenotypic variation at least indicates that natural selection could have acted upon such phenotypic variation.

Although linkage disequilibrium (LD) among SNPs is high, the fact that HPG1 genomes differ in very few SNPs greatly reduces the list of candidate loci that might generate the observed phenotypic variation (Fig. S6) (44). With this reasoning in mind and understanding the limitations imposed by LD, we carried out a genome-wide association (GWA) analysis and found 79 SNPs associated with one or more root traits, mostly growth and directionality (Fig. 4). Twelve SNPs were in coding regions and seven resulted in nonsynonymous changes — some producing non-conservative amino-acid changes and thus likely to affect protein structure and/or function (Table 1, based on transition scores from (45)). Due to the aforementioned LD, in some cases the results of associations could not be confidently assigned to a specific SNP and thus we report the number of other associated mutations with r^2^ > 0.5 (Table 1, Fig. S6). For other cases, we were able to pinpoint clear candidates that were not in LD with other SNPs and whose functional annotation had a strong connection to the phenotype (Table 1, Fig. S6). For example, one SNP associated with root gravitropism was not linked to any other SNP hit and it was found at 40% frequency (top 3% percentile). This SNP produces a cysteine to tryptophan change in AT5G19330, which is involved in abscisic acid response and confers salt tolerance when overexpressed (46). Another nonsynonymous SNP associated with root growth is located in AT2G38910, which encodes a calcium-dependent kinase that is a factor regulating root hydraulic conductivity and phytohormone response *in vitro* (47,48).

**Table 1.**
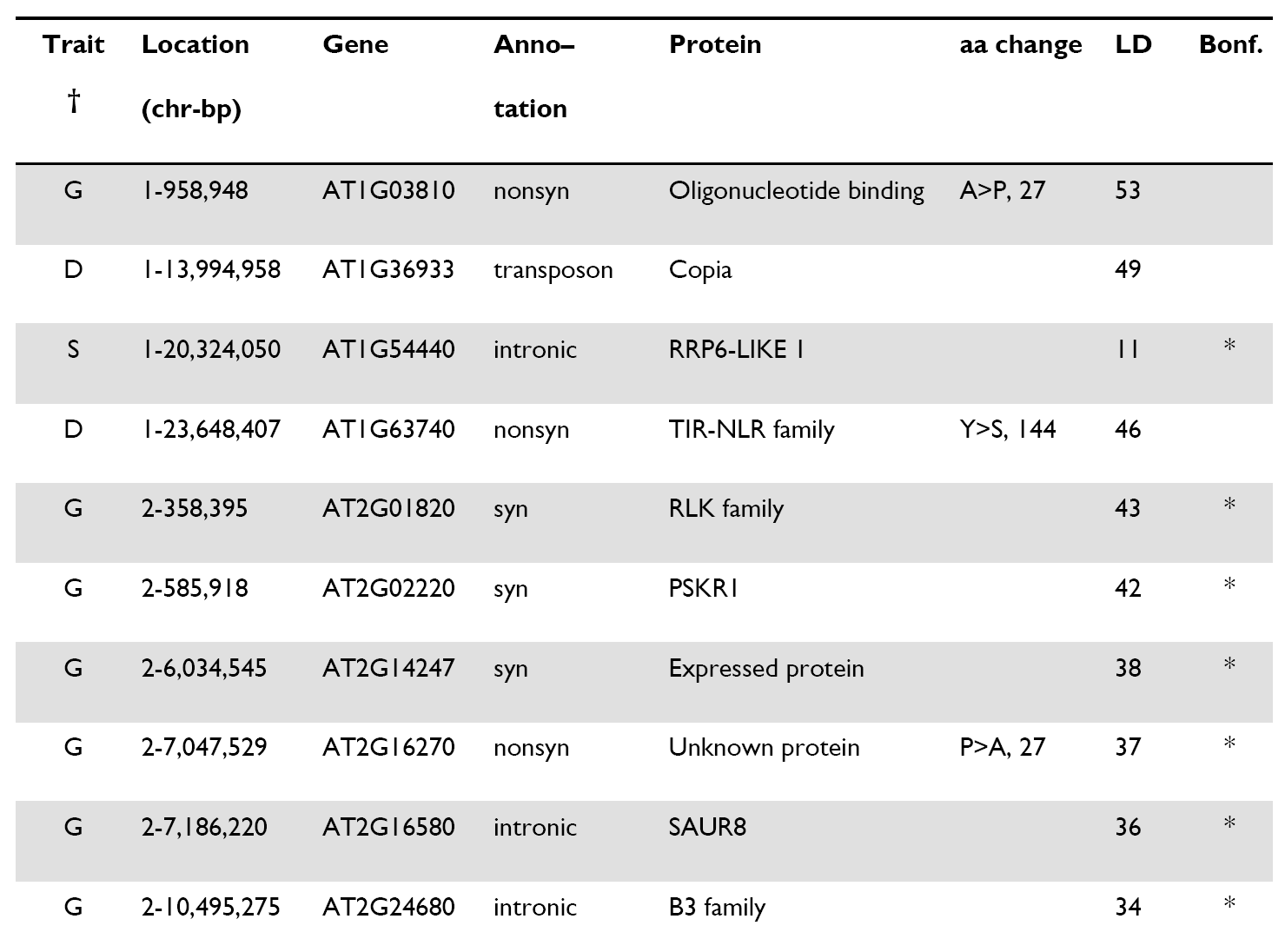
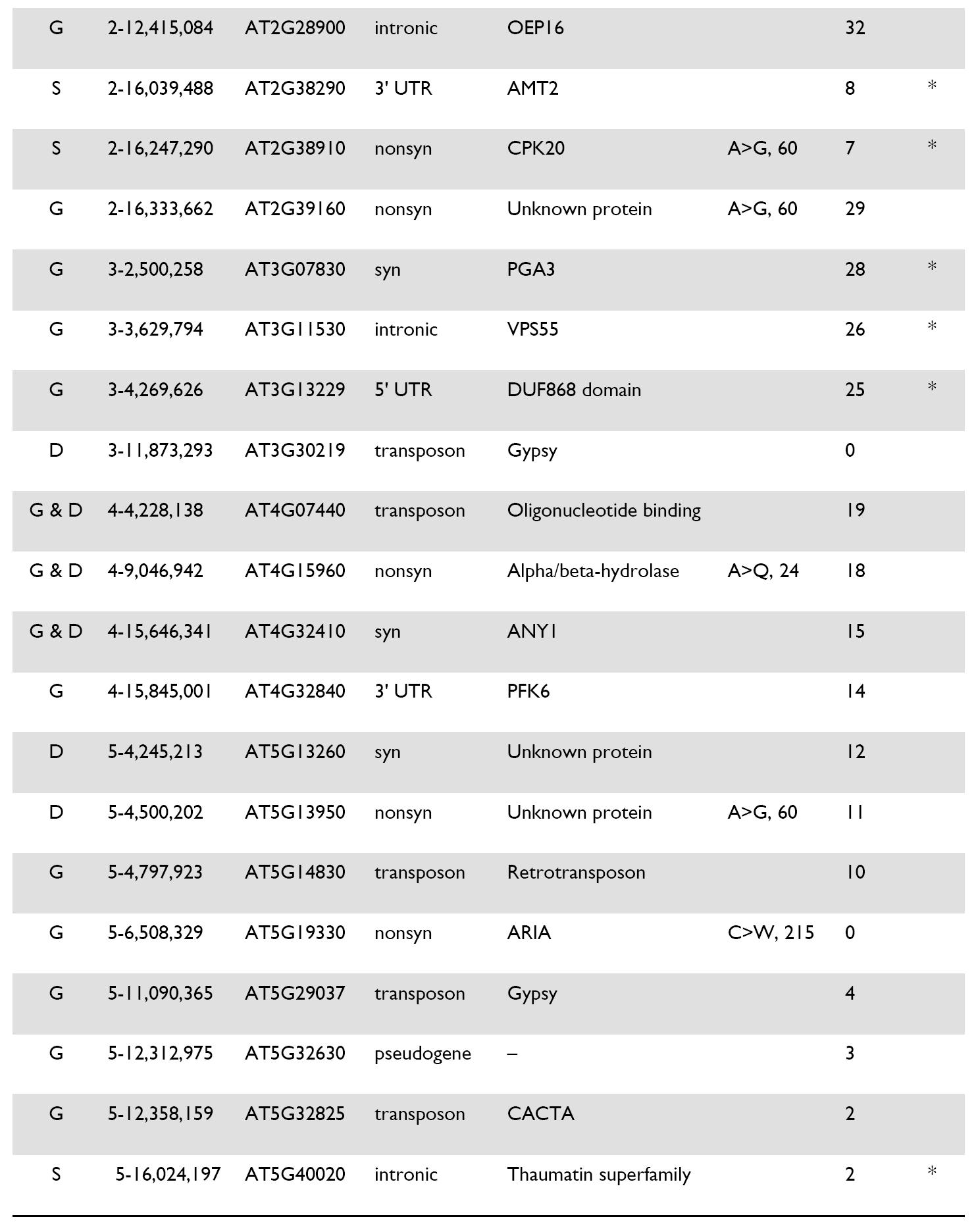
Genic SNPs associated with different traits. For nonsynonymous SNPs, the amino acid change and the Grantham score (ranging from 0 to 215), which measures the physico-chemical properties of the amino acids, are reported. All SNPs in the table were significant (p < 0.05) after raw p-values were corrected by an empirical p-value distribution from a permutation procedure. * highlights those that also passed a double Bonferroni threshold, correcting by number of SNPs and number of phenotypes (p < 0.0001). LD corresponds to how many other SNP hits are in high linkage (r^2^>0.5). Table S5 contains information on all significant SNPs and Table S4 for details on phenotypes and climatic variables.

**Figure 4.**
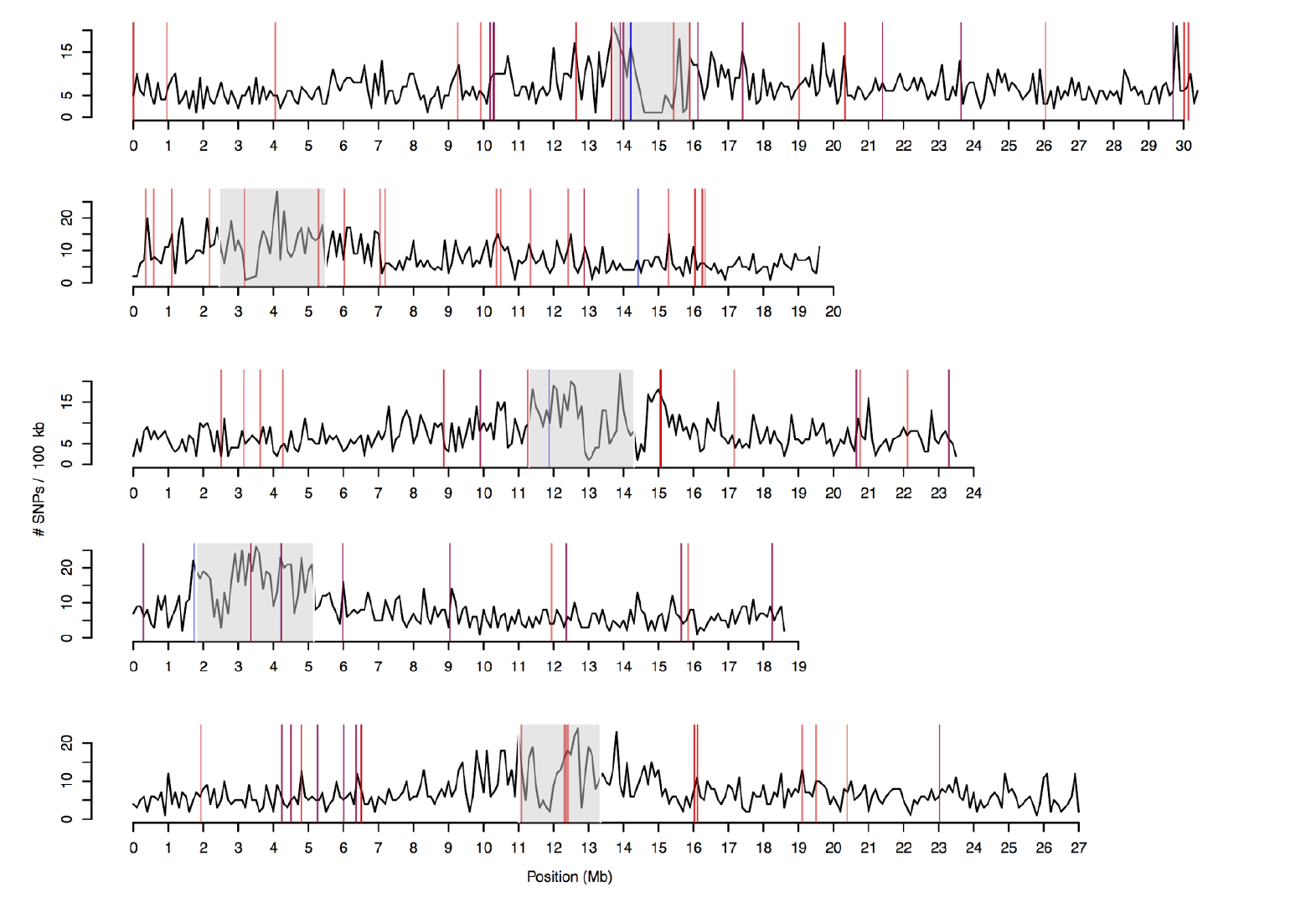
Density of SNPs along all chromosomes and location of GWAS hits. Black line shows number of SNPs per 100 kb window. Centromere locations are indicated by grey shading. Vertical lines indicate SNPs associated with root phenotypes (red) and climatic variables (blue) (Table 1 and Table S5).

Nineteen other SNPs were associated with climate variables after correction for latitude and longitude (www.worldclim.org, Table S4), and generally tended to coincide with top root-associated SNPs (odds ratio = 3.9, Fisher’s Exact test p = 0.002; Fig. 4, and Table S5). Specifically, this means that alleles increasing root length and gravitropic growth were present in areas with lower precipitation, and *vice versa* (Pearson’s correlation r=0.85, p=0.003). This indicates that phenotypic variation generated by mutations coincides with environmental (and not geographic) gradients along the colonized areas. Compared to other mutations with matched allele frequencies, root-associated mutations are first found in older herbarium samples nearer to Lake Michigan (Fig. S5), the area in US that seems to be most populated by *A. thaliana* (21). This could be explained by natural selection having maintained mutations with phenotypic effect for a longer time than neutral mutations or perhaps that this mutations were selected for in a new environment. All in all our results are compatible with natural positive selection having already acted on root morphology variation that was generated by *de novo* mutations in this colonizing lineage. To confirm such hypotheses of local adaptation by *de novo* mutations, it will be necessary to grow collections of divergent HPG1 individuals in multiple contrasting locations over several years, and ideally revive historical specimens to compare performance (49).

### Conclusions

In summary, we have exploited whole-genome information from historic and contemporary collections of a herbaceous plant to empirically characterize evolutionary forces during a recent colonization. With this natural time series experiment we could directly estimate the nuclear substitution rate in wild *A. thaliana* populations – a parameter difficult to characterize experimentally (9). This allowed us to date the colonization time and spread of HPG1 in N. America. We provide evidence that purifying selection has already changed the site frequency spectrum in the course of just a few centuries. Finally, we discovered that a small number of *de novo* mutations that rose to intermediate frequency can together explain quantitative variation in root traits across environments. This strengthens the hypothesis that some *de novo* variation could have had an adaptive value during the colonization and expansion process, a hypothesis that has been put forward as one of the possible solutions to the genetic paradox of invasion in plants (17). This process might be more relevant in self-fertilizing plants, which typically have less diversity than outcrossing ones (50), but have higher growth rates (43) and account for the majority of successful plant colonizers (5). While *A. thaliana* HPG1 is not an invasive, i.e. harmful, species, it can teach us about fundamental evolutionary processes behind successful colonizations and adaptation to new environments. Our work should encourage others to search for similar natural experiments and to unlock the potential of herbarium specimens to study “evolution in action”.

## METHODS

### Sample collection and DNA sequencing

Modern *A. thaliana* accessions were from the collection described by Platt and colleagues (23), who identified HPG1 candidates based on 149 genome-wide SNPs (Table S1, Supplementary Text 1). Herbarium specimens were directly sampled by Max Planck colleagues Jane Devos and Gautam Shirsekar, or sent to us by collection curators from various herbaria (Table S1, Supplementary Text 1). Among the substantial number of specimens in the herbaria of the University of Connecticut, the Chicago Field Museum and the New York Botanical Garden, we selected herbarium specimens spaced in time so there was at least one sample per decade starting from the oldest record (1863). The differences in geographic biases of herbarium and modern collections are difficult to know (2), thus we did choose both historic and modern samples that were as regularly distributed in space as possible, and sample overlapping locations wherever possible. DNA from herbarium specimens was extracted as described (51) in a clean room facility at the University of Tübingen. Two sequencing libraries with sample-specific barcodes were prepared following established protocols, with and without repair of deaminated sites using uracil-DNA glycosylase and endonuclease VIII (refs. (52–54)) (Supplementary Text 2). We also investigated patterns of DNA fragmentation and damage typical of ancient DNA (24) (Supplementary Text 2). DNA from modern individuals was extracted from pools of eight siblings using the DNeasy plant mini kit (Qiagen, Hilgendorf, Germany). Genomic DNA libraries were prepared using the TruSeq DNA Sample or TruSeq Nano DNA sample prep kits (Illumina, San Diego, CA), and sequenced on Illumina HiSeq 2000, HiSeq 2500 or MiSeq instruments. Paired-end reads from modern samples were trimmed and quality filtered before mapping using the SHORE pipeline v0.9.0 (25,55). Because ancient DNA fragments are short (Fig. S1) we merged forward and reverse reads for herbarium samples after trimming, requiring a minimum of 11 bp overlap (51), and treated the resulting as single-end reads. Reads were mapped with GenomeMapper v0.4.5s (56) against an HPG1 pseudo-reference genome (25), and against the Col-0 reference genome, and SNPs were called with SHORE for the HPG1 pseudo-reference genome mappings (25,57) using different thresholds (Supplementary Text 3). Average coverage depth, number of covered genome positions, and number of SNPs identified per accession relative to HPG1 are reported in Table S1. We also re-sequenced the genomes of twelve Col-0 MA lines (57,58) (Table S2) (Supplementary text 4) to recalculate and update the laboratory mutation rate from Ossowski et al. (38) with the newer sequencing technologies.

### Phylogenetic methods and genome-wide statistics

We used the Pegas, Ape and Adegenet packages in R (59–61) to manipulate and visualize the genetic distances of all samples as well as the HPG1 subset (Supplementary Text 7). We constructed parsimony networks using SplitsTree v.4.12.3 (62), with confidence values calculated with 1,000 bootstrap iterations. We built Maximum Clade Credibility Trees using the Bayesian phylogenetic tools implemented in BEAST v.1.8 (63) (see below).

We estimated genetic diversity as Watterson’s θ (64) and nucleotide diversity π, and the difference between these two statistics as Tajimas’s *D (65)* using DnaSP v5 (66). We estimated pairwise linkage disequilibrium (LD) between all possible combinations of informative sites, ignoring singletons, by computing *r*^2^, *D* and *D*’ statistics using DnaSP v5 (66). For the modern individuals, we calculated the recombination parameter rho (*4N*_*e*_*r*) also using DnaSP v5 (66).

### Substitution and mutation rate analyses

Similarly as in Fu et al. (67), we used genome-wide nuclear SNPs to calculate pairwise “net” genetic distances using the equation *D′*_ij_ = *D*_ic_-*D*_jc_, where *D*′ _ij_ is the net distance between a modern sample *i* and a herbarium sample *j*; *D*_ic_ the distance between the modern sample *i* and the reference genome *c*; and *D*_jc_ is the distance between a modern sample (j) and the reference genome (c). We calculated a pairwise time distance in years between the collection times, *T*′ij, and calculated the linear regression: *D*′ = *a* + *bT*′. The slope coefficient *b* describes the number of substitution changes per year. We used either all SNPs or subsets of SNPs at different annotations (genic, intergenic etc.) appropriately scaled by accessible genome length. Because the points used to calculate the regression are non-independent, a bootstrap has been recommended to overcome to a certain extent the anti-conservative confidence intervals (30) (Supplementary Text 7 and Fig. S3).

To fully account for the non-independence of points, we need to work with phylogenies. The Bayesian phylogenetics approach we used is implemented in BEAST v1.8 (63) and is called tip-calibration, and calculates a substitution rate along the phylogeny. Our analysis optimized simultaneously and in an iterative fashion using a Monte Carlo Markov Chain (MCMC) a tree topology, branch length, substitution rate, and a demographic Skygrid model (Supplementary Text 7). The demographic model is a Bayesian nonparametric one that is optimized for multiple loci and that allows for complex demographic trajectories by estimating population sizes in time bins across the tree based on the number of coalescent-branching-events per bin (68). We also performed a second analysis run using a fixed prior for substitution rate of 3x10^−9^ substitutions site ^−1^year^−1^based on our previous net distance estimate to confirm that the MCMC had the same parameter convergence, e.g. tree topology, as in the first “estimate-all-parameters” run.

Having a substitution rate per year we can estimate the time to the most common recent ancestor *L* solving *d = 2L x μ* where *d* is the average pairwise genetic distance between our samples and *μ* is the calculated substitution rate from the distance method. This yielded 363 years, which subtracted to the average collection date of the samples, produced a point estimate of 1615. We compare this estimate with the inferred phylogeny root from the BEAST analysis.

### Inference of genome-wide selection

We separately analyzed sequences at different annotations, since as they might be under different selection regimes (i.e. evolutionary constraints). We computed one-tailed Fisher’s exact test using the base stats package in R (69) on tables of counts of the total number of positions in the genome annotated as a coding or non-coding (intergenic, intronic, all other noncoding) and the number of SNPs of each annotation present in the HPG1 dataset:

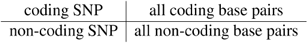

The test will return whether coding regions have a lower number of SNPs than other reference annotation (intronic, interenic, all non-coding regions), as expected by the total number of positions in the genome annotated as such. We also constructed contingency tables to test whether the SNPs are more likely to be found at low (<5%) or intermediate (5≥%) frequency:

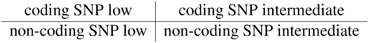

Finally, we calculated the unfolded Site Frequency Spectrum (SFS) based on the order of appearance of genetic variants in the herbarium dataset. We then used the the Kolmogorov–Smirnov two-samples test and 10,000 bootstrap resampling using the R package Matching v. 4.9-2 (ref. (70)) to calculate whether the frequency spectrum was lower for coding SNPs than for other SNPs. Additionally, we also repeated these analyses comparing nonsynonymous and synonymous mutations.

### Association analysis

We collected flowering, seed and root morphology phenotypes for 63 accessions (Supplementary Text 8). For associations with climate parameters, we followed a similar rationale as previously described (71). We extracted information from the bioclim database (http://www.worldclim.org/bioclim) at a 2.5 degrees resolution raster and intersected it with geographic locations of HPG1 samples (n = 100). We performed association analyses under several models and *p*-value corrections using the R package GeneABEL (72) (Supplementary Text 8.2). To calculate the variance of the trait explained by all genetic variants, we used a linear mixed model: *y = Xb + Zu + ε*; where *y* is the phenotype or climate variable, *X* is the genotype states at a given SNP, *b* is the fixed phenotypic effect of such SNP, *Z* is the design matrix of genome identities, *u* is the random genome background effect informed by the kinship matrix and distributed as MVN (0, *σ* _*g*_*A*), and *ε* is the random error term. The ratio of *σ* _*g*_ */ σ*_*T*_ is commonly called narrow sense heritability, “chip” heritability, or proportion of variance explained by genotype (73). Only SNPs with MAF>5% (n = 391) were used to build a kinship or relationship matrix *A*. Note that the differences between any two genotypes were of the order of one or few dozens of SNPs. While this approach is appropriate to calculate a chip heritability, it would not be very useful to detect significant SNP, as the random factor accumulates all the available variation (Table S4). We therefore run regular GWA model without kinship matrix: *y = Xb + ε*; but generated a p-value empirical null distribution based on running such model over 1,000 permuted datasets, which lead to conservative significance calculation (Fig. S6, Data Appendix S1). The p-values from running the association in the real data that were below the 5% tail in the empirical distribution could be considered significant. However, we also established a conservative “double” Bonferroni correction, where the significant threshold was lowered to 0.01% (= 5% / [number of SNPs + number of phenotypes tested]). All significant SNPs are shown in Table S5, and a subset in Table 1. Although many phenotypic traits did not have significant SNPs, we show all the QQ plots in the Data Appendix S1 file.

#### Accession numbers

Short reads have been deposited in the European Nucleotide Archive under the accession number XXXXX.

#### Online Content

This article contains supplementary information including data sets, extended methods and supplementary figures at xxx.

## Acknowledgments

For providing and retrieving herbarium specimens, we thank R. Capers, J. Devos, G. Shirsekar, M. S. Dossmann, J. Freudenstein, C. M. Herring, C. Niezgoda, C. A. McCormick, J. Peter and M. Thines. We thank X. Zhao and I. Henderson for recombination estimates, C. Lanz for sequencing support, C. Goeschl, B. Zierfuss and B. Wohlrab for help with root analyses, and P. Lang, D. Seymour, and D. Koenig for thorough proofreading and comments on the manuscript. We thank to Robert Colautti for useful comments on the theoretical framing of the manuscript, M. Nordborg for discussions and pointing us to the work of A.R. Templeton, K. Pruefer for input on data analysis, and the Weigel and Burbano labs for comments. Supported by the President’s Fund of the Max Planck Society (project “Darwin”), ERC (AdG IMMUNEMESIS) and core funds of the Max Planck Society.

## Author Contributions

H.A.B. and D.W. conceived and supervised the project, and coordinated the collaborative effort. J.B. coordinated the collection of modern seed samples. C.J., B.B. and J.B. performed and analyzed flowering time and seed set greenhouse experiments. C.S. and R.S. performed and analyzed root assays and seed size measurements under the supervision of W.B.; C.B. and J.H. sequenced and curated modern samples, coordinated by D.W.; H.A.B. coordinated the collection and analysis of herbarium samples. J.K. coordinated the extraction of DNA and library preparation of herbarium samples. V.J.S. and E.R. prepared sequencing libraries from herbarium specimens. C.B. called variants in HPG1. J.H. called variants in mutation accumulation lines. M.E.A. performed the population and quantitative genomic analyses with supervision of R.N., C.B. and H.A.B. The first draft was written by M.E.A. and the final manuscript was written by M.E.A., C.B., H.A.B. and D.W. with comments from all coauthors.

Authors declare no conflict of interests.

## Supplemental Information for

Exposito-Alonso, Becker et al.:

## SUPPLEMENTAL TEXT

### 1. Sample collection and preparation

Seeds from modern accessions (Table S1) were bulked at the University of Chicago. Progeny for DNA extraction was grown at the Max Planck Institute for Developmental Biology. We used 2 to 8 mm^2^ of dried tissue for destructive sampling from the herbarium specimens (Table S1).

### 2. Authenticity of aDNA

First, unrepaired sequencing herbarium libraries were screened for authenticity by sequencing at low coverage on Illumina HiSeq 2500 or MiSeq instruments. To verify the DNA retrieved from historical samples of *A. thaliana* was authentic, we checked the percentage of endogenous DNA of the sample (Fig. S1A) as well as typical postmortem DNA damages: high fragmentation of DNA (Fig. S1B), enrichment of substitution from C to T at the first base pair (Fig. S1C) as well as purine enrichment at breakpoints of DNA fragments (Fig. S1D) (for details see (1)). Sequencing to produce the final genomes (101 bp paired end) was carried out on an Illumina HiSeq 2000 instrument after DNA repair by uracil-DNA glycosylase (2–4). For a detailed analysis of authenticity in a fraction of our samples, see Weiss et al. (1).

### 3. SNP calling thresholds

To assess the effect of SNP calling thresholds on the mutation rate, we employed three different SHORE v0.9.0 quality thresholds following previous work (see Table S4 from (5)): allowing at most one intermediate penalty in all strains (most stringent threshold; “32-32”); requesting that at least one strain had at most one intermediate penalty, while all others were allowed up to two high and one intermediate penalties (intermediate stringency, “32-15”); and finally allowing one high and one intermediate penalty for all strains (most lenient stringency, “24-24”). On top of that, we would either allow missing information per SNP in up to 50% of accessions, or request complete information (0% missing rate). Thus, the most rigorous case would be 32-32 quality and 0% missing rate, and the most relaxed 24-24 quality and 50% maximum missing rate. Substitution rate calculations (section 7.2) were done for datasets from all combinations of these quality parameters (Fig. S3), and we chose the regular 32_15 quality threshold and complete information for the final estimate (Fig 3 C, E).

### 4. Resequencing of Col-0 Mutation Accumulation lines

We also sequenced the genomes of twelve greenhouse-grown mutation accumulation (MA) lines, including ten that had been sequenced at lower coverage before (5,6) (Table S2). We called SNPs, indels and structural variants (SVs), following the workflow and parameters described (7), but without iterations. This procedure resulted in 2,203 polymorphisms shared by all lines, indicating errors in the reference sequence (12% of variants replaced N’s in the TAIR9 genome) or genetic differences in the founder plant of the MA population compared to the Col-0 reference genome. In addition, we identified 388 segregating variants across the twelve lines (Table S2), of which 350 were singletons. This analysis revealed on average 25.5 SNPs, 4.9 deletions and 3.2 insertions per MA line at the 31^st^ generation (Table S2), compared to 19.6 SNPs, 2.4 deletions and 1.0 insertions previously detected in the 30^th^ generation with shorter read length and lower read depth (8). The genome length accessed in this sequencing effort, 115,954,227 bp, was used to scale the number of point mutations to a rate of 7.1 x 10^−9^ mutations site^−1^ generation^−1^ (Table S3, Fig. 3E).

### 5. Identification of *bona fide* HPG1 accessions and mutations

#### 5.1 HPG1 and other haplogroups in North America

The modern samples had been originally selected based on previous genotyping efforts of about 2,000 N. American accessions with for 149 nuclear, intermediate-frequency SNPs. This work had pointed to there being a single haplogroup, HPG1, that was invariant at these 149 markers and that accounted for about half of N. American individuals genotyped (9). We extracted from the 123 genomes we had completely sequenced the same 149 SNPs and built a neighbour joining tree (Fig. S1A). We also built the same tree with the whole-genome sequences (Fig. S1B), which was mostly in agreement with the 149 SNP tree.

The previous work had identified several other haplogroup in N. America (9). Not surprisingly, HPG1 individuals outcross with other lineages, and this accounts for some of the individuals which we later removed, because they did not agree completely in all 149 markers with the HPG1 consensus.

#### 5.2 North american private diversity

Having identified these *bona fide* HPG1 individuals, we wanted to confirm that the diversity has a legitimate origin from *de novo* mutations. For that we used the 1001 Genomes resource (www.1001genomes.org), which covers a sampling of populations from the native Eurasian and African range. Subsetting the genomes from this resource to only European accessions, and limiting the SNP set to those with ≥1% frequency of alternative alleles and a maximum of 50% missing data (the same quality rate as our HPG1 SNP call), there were 300 variants out of all 5,181 HPG1 variants that were also found in Europe or Asia (5.7%). Changing the maximum missing data to 10% we get a more conservative estimate of 1.8% overlap, while increasing the maximum missing data to 90%, we get the anti-conservative estimate of 6.5% overlap. Only one of the reported SNPs associated with phenotypes (see Section 8) was among these shared variants.

There are several scenarios that can explain these shared SNPs. One is simply that there was not a single founding seed, but a few of closely related individuals coming from the native range. Other explanations are that parallel mutations occurred in North America and Eurasia, that HPG1 individuals were reintroduced to Europe, or that reversion-mutation occurred in some HPG1 individuals. The latter is not implausible given the large population size of the species and the fact that about 10% of all sites in the genome are SNPs in the 1001 Genomes collection. As explained in the main text, SNP sharing due to admixture with other lineages is extremely unlikely, as such cases should be evident as blocks of high SNP diversity along the genome (Fig. 4).

Finally, regarding chloroplast diversity, we did not find any SNP in the chloroplast of HPG1 individuals. This is probably because chloroplast mutation rates are much slower (10) and because the founder colonizers actually came from a small batch of seeds from an identical mother (chloroplast diversity in the native range is of 2,842 SNPs (11)).

### 6. Extent of linkage disequilibrium and recombination

We estimated pairwise linkage disequilibrium (LD) between all possible combinations of informative sites, ignoring singletons, by computing *r*^2^, *D* and *D*′ statistics. LD decay was estimated using a linear regression approach. Linkage disequilibrium parameter |*D*′| did not decay with physical distance (intercept = 0.99, slope = 0.00) among all SNP pairs. Indeed 99.975% of pairwise SNP comparisons had |*D*′| = 1 meaning that 99.975% of those comparisons only three out of the four possible gametes (ab, aB, Ab, AB) are found and thus mutation alone can explain their existence without the need of invoking recombination. In other words, such three gametes can be represented in a tree structure. LD and recombination related statistics were determined using DnaSP v5 (12).

### 7. Substitution and mutation rate analyses

#### 7.1 Greenhouse grown MA lines

Mutation rates were estimated for each 31^st^ generation greenhouse-grown MA line (5) as the number of mutations divided by the total bp length of the genome (or a given annotation) and by 31 generations (the two MA lines with only three generations were excluded from this analysis). Mean and confidence intervals across lines are reported (Table S3). The genome length was determined as all base pairs with coverage higher or equal to 3, and a SHORE mapping quality score of at least 32 in one sample (Table S2).

#### 7.2 Natural populations of HPG1

##### 7.2.1 Net distances

For the “net genetic distances” method, we computed confidence intervals of the *b* regression slope coefficient (*D*′ = *a* + *bT*′) using a bootstrap with replacement of 1,000 samples to avoid over-confident confidence intervals due to lack of independence of points (13). We used either all SNPs or SNPs at specific annotations to calculate different substitution rates and scaled the slope into a per-base rate using all positions (of the given annotation) that passed alternative or reference call quality thresholds rather than using a single value of genome length (Table S3). For all annotations we calculated substitution rates with three quality thresholds and either full information per SNP or allowing a maximum of 50% missing accessions per SNP (see Section 3 and Fig. S1C).

For some annotations substitution rates were not reliable. For instance, in 3’ and 5’ UTR regions, we did not have enough mutations (on average ∽1 SNP difference between any pair), and thus do not report these regions’ rates. We could also have less power to discover SNPs in annotations with extensive structural variation such as active transposable elements (14). Transposons, which comprise ∽8% of the genome and ∽19% of all the SNPs in greenhouse MA lines, had fewer SNPs called than expected in HPG1. This would explain the atypically low transposon substitution rate (Table S3). Therefore, transposon substitution rates in HPG1 cannot be trusted.

##### 7.2.2 Bayesian tip-calibration

For the second approach to estimate a substitution rate, the Bayesian phylogenetics tip-calibration approach, we performed systematic runs and chain convergence assessments of different demographic and molecular clock models. We found the Skygrid demographic model (15) and the lognormal relaxed molecular clock (16) the most appropriate models. Under a relaxed molecular clock, the substitution rate is allowed to vary across branches with a lognormal distribution. The prior used for molecular clock was a Continuous-Time Markov Chain (CTMC) (15,17). The analysis was carried out remotely at CIPRES PORTAL (v3.1 www.phylo.org) using uninformative priors. The run took about 1,344 CPU hours and performed 1,000 million steps in a Monte Carlo Markov Chain (MCMC), sampling every 100,000 steps. Burn-in was adjusted to 10% of the steps. To visualize the tree output we produced a Maximum Clade Credibility (MCC) tree with a minimum posterior probability threshold of 0.8 and a 10% burn-in using TreeAnnotator (part of BEAST package), and visualized the MCC tree using FigTree (tree.bio.ed.ac.uk/software/figtree/) (Fig. 3B). Additionally, we used DensiTree (18) to simultaneously draw the 10,000 BEAST trees with the highest posterior probability (Fig. 3A). Since all trees were drawn transparently, agreements in both topology and branch lengths appear as densely colored regions, while areas with little agreement appear lighter.

##### 7.2.3 Methylation status of mutated sites

As in many other species, the spectrum of *de novo* mutations in the greenhouse-grown *A. thaliana* MA lines is biased towards G:C*→*A:T transitions (8), leading to an inflated transition-to-transversion ratio (Ts/Tv). This bias is less pronounced in recent mutations in a Eurasian collection of natural accessions (Fig. 5A of (19) and in HPG1 accessions (Fig. 3D). A recent multigenerational salt stress experiment in the greenhouse also showed a more balanced Ts/Tv (20). These findings indicate that less benign conditions might promote a lower Ts/Tv, and one possible cause are methylation patterns, known to change under different environments (21).

We interrogated the potential evolutionary role of cytosine methylation in the mutability of cytosine bases in the HPG1 accessions. For reference DNA methylation data, we used previously generated bisulfite-sequencing data of HPG1 strains (7) and of Col-0 MA lines (5), respectively. For both datasets, methylation status was calculated as the fraction of reads with methylated cytosines by the total number of reads at a certain cytosine position in the genome. Our rationale was that if methylation affected mutability, the degree of methylation at positions were we find a new mutation should be higher. To be sure that a given site in HPG1 was a new mutation, we only considered positions for which we could determine that state by alignment to the *A. lyrata* genome (22). The “tested sites” were positions in HPG1 that had a mutation both from *A. lyrata* and *A. thaliana* Col-0. These positions can be of two kinds, “fixed” if all HPG1 individuals carry the alternative, or “segregating” if both reference and alternative alleles exist in HPG1. As control, “control set”, we used cytosine positions that did not vary across HPG1, *A. lyrata* and *A. thaliana*. To produce the methylation distribution of the control set we randomly chose 1,000 invariant cytosine positions. For the test sets, we averaged the methylation degree and compared it with the control distribution.

Ancestral cytosines with higher methylation in both *A. thaliana* Col-0 reference and HPG1 pseudo-reference methylome datasets were more likely to mutate to thymines in HPG1 (Fig. S2 A-D). Additionally, the methylation degree at substitutions inside genes was higher in the HPG1 methylome (Fig. S2 B,D). While some C*→*T changes could be explained by higher spontaneous deaminations known to happen more often at methylated cytosines, also C*→*A / G substitutions were more likely to have been methylated. If this process is common enough, the Ts / Tv ratio should decrease. We are far from understanding differences in Ts / Tv in natural and controlled conditions, but definitely methylation status seems to have a strong statistical connection with mutability.

### 8. Phenotypic association analyses and dating of newly arisen mutations

#### 8.1. Phenotyping

##### 8.1.1 Root

Fifteen root phenotypes were scored for ≥ 10 replicates per genotype over a time-series experiment at the Gregor Mendel Institute in Vienna, using image analysis as described in detail elsewhere (23). We used the means per genotypes and per time series for association analyses.

##### 8.1.2 Seed size

We spread the seeds of given genotypes on separate plastic square 12 x 12 cm Petri dishes. For faster image acquisition we used a cluster of eight Epson V600 scanners. The scanner cluster was operated by the BRAT Multiscan image acquisition tool (www.gmi.oeaw.ac.at/research-groups/wolfgang-busch/resources/brat/). The resulting 1600 dpi images were analyzed in Fiji software. Scans were converted to 8-bit binary images, thresholded (parameters: setAutoThreshold(“Default dark”); setThreshold(20, 255)) and particles analyzed (inclusion parameters: size = 0.04-0.25 circularity = 0.70-1.00). The 2D seed size was measured in square millimeters (parameters: distance = 1600 known = 25.4 pixel = 1 unit = mm) for 2 plants per genotype, > 500 seeds per plant.

##### 8.1.3 Flowering in the growth chamber

We estimated the flowering time in growth chambers under four vernalization treatments (0, 14, 28 and 63 days of vernalization). We grew 6 replicates per accession divided between two complete randomized blocks for each treatment. Seeds were sown on a 1:1 mixture of Premier Pro-Mix and MetroMix and cold stratified for 6 days (6°C, no light). We then let plants germinate and grow at 18°C, 14 hours of light, 65% humidity. After 3 weeks, we transferred the plants to vernalization conditions (6°C, 8 hours of light, 65% humidity). After vernalization, plants were transferred back to long day conditions. Trays were rotated around the growth chambers every other day throughout the experiment, under both vernalization and ambient conditions. Germination, bolting and flowering dates were recorded every other day until all plants had flowered. Days till flowering or bolting times were calculated from the germination date until the first flower opened and until the first flower bud was developed, respectively. The average flowering time and bolting time per genotype were used for association analyses.

##### 8.1.4 Fecundity in the field

To investigate variation in fecundity in natural conditions, we grew three replicates of each accession in a field experiment following a completely randomized block design. Seeds were sown from 09/20/2012 to 09/22/2012 in 66-well trays (well diameter = 4 cm) on soil from the field site where plants were to be transplanted. The trays were cold stratified for seven days before being placed in a cold frame at the University of Chicago (outdoors, no additional light or heat, but watered as needed and protected from precipitation). Seedlings were transplanted directly into tilled ground at the Warren Wood field station (41.84° N., 86.63° W.), Michigan, USA on 10/13/2012 and 10/14/2012. Seedlings were watered-in and left to overwinter without further intervention. Upon maturation of all fruits, stems were harvested and stored between sheets of newsprint paper. To estimate the fecundity, stems were photographed on a black background and the size of each plant was estimated as the number of pixels occupied by the plant on the image. This measure correlates well with the total length of siliques produced, a classical estimator of fecundity in *A. thaliana* (Spearman’s rho=0.84, *p -* value<0.001, data not shown).

#### 8.2 Quantitative genetic analyses

For 63 modern accessions, we measured time to bolting and flowering, seeds per plant, seed size, and 15 root phenotypes in common chamber or common garden settings. For all 100 accessions, climatic information from the bioclim database (www.worldclim.org/bioclim) was extracted using their geographic coordinates. For historic samples, some locations were only known by county name. In this case we assigned the geographic coordinate location of the centroid of the county.

##### 8.2.1 Heritability

We performed association analyses using the R package GenABEL (24), with measured phenotypes (p = 25) and climatic variables (c = 18) as response variables and SNPs as explanatory variables. A Minimum Allele Frequency (MAF) cutoff of 5% was used. The number of assessed SNPs was 391 in a dataset of only modern samples but with imputed genotypes for missing data using Beagle v4.0 (25), and 456 SNPs with a dataset of modern and historic samples, without imputation. For all associations, at least 63 individuals were genotyped for a specific SNP. We first investigated broad sense heritability (*H*^*2*^) of each trait using ANOVA partition of variance between and within lines using replicates (Table S4). Significance was obtained by common F test in ANOVA. Secondly we used the *polygenic_hglm* function to fit a genome wide kinship matrix to calculate a narrow sense heritability estimate (*h*^*2*^). This fits a model of the type *y = Zu + ε* (see Main text Methods). Significance was calculated employing a likelihood ratio test comparing with a null model. In principle, *h*^*2*^ is a component of *H*^*2*^, then its values should theoretically be *h*^*2*^ < *H*^*2*^. That is not our case. Our result cannot be interpreted in this framework, since the calculation of both was not done with the same samples: for the *h*^*2*^ calculation we employed genotype means whereas for the *H*^*2*^ we used multiple replicated measurements per genotype. The averaging of replicates per genotype in *h*^*2*^ reduced environmental and developmental noise and thus we would expect *h*^*2*^*>H*^*2*^. We did this so the climatic estimates of h^*2*^, for which we only have one value per genotype, would be comparable with the phenotypic h^*2*^ ones (Table S4).

##### 8.2.2 Linear Models

For association analyses we first employed a linear mixed model that fitted the kinship matrix using the *mmscore* function. This model is of the type: *y = Xb + Zu + ε* (see Main text Methods) (26). Only three significant SNP hits were discovered using a 5% significance threshold after False Discovery Rate correction (FDR). This was expected since we have few variants and these would have originated in an approximated phylogeny structure. We concluded that fitting the kinship matrix in our model was not appropriate since there would be no residual variation for association with specific SNPs. With this rationale we employed a fixed effects linear model using the *qtscore* function (27). This model is of the type: *y = Xb + ε*; where no random effect of genome background is fit. To reduce the risk of having false-positives, we took a conservative permutation strategy by carrying out association with over 1,000 randomized datasets (permuting phenotypes across individuals) and used the resulting empirical p-value distribution to correct p-values estimated with the original dataset. SNPs with p-values below 5% in the empirical p-value distribution should be considered significant (but see next section). In climatic models, we included longitude and latitude as covariates to correct for any spurious association between SNPs and climate gradients created by the migratory pattern of isolation by distance.

##### 8.2.3 Evaluation of significance

Significant SNPs were interspersed throughout the genome (Fig. 4) and their p-values and phenotypic effects did not correlate with the minimum age of the SNPs nor with their allele frequency, something that could have indicated that the significance was merely driven by the higher statistical power of intermediate frequency variants. Using QQ plots to assess inflation or deflation of p-values, we observed generally that permutation corrected p-values were deflated — another evidence of our conservative strategy. Straight horizontal series of points in QQ plots indicate that multiple SNPs have identical p-values, a pattern that we attributed to long range LD, i.e. lack of independence (see Data Appendix S1 for trait distributions and QQ plots from each association analysis).

To further ensure that we avoided false positive results, we also prioritized SNPs whose empirical p-value was not below 5% only but also below 5% / (number of SNPs + number of traits) = 0.01%. This “double” Bonferroni correction was very conservative (Table 1, Table S5).

##### 8.2.4 Context of de novo mutations associated with phenotypes

For each SNP in our dataset, we determined the ancestral and derived states, by identifying which allele was found in the oldest herbarium samples. We compared the time of emergence and the centroid of geographic distribution of the alternative alleles of SNP hits to random draws of SNPs with the same MAF filtering (5%) (Fig. S1).

##### 8.2.5 Functional information

On top of phenotypic and climatic associations of SNP hits, we also provide a likely functional effect employing a commonly used amino acid matrix of biochemical effects (28). Functional information of gene name and ontology categorization of SNP hits was obtained from www.arabidopsis.org/portals/genAnnotation/gene_structural_annotation/annotation_data.jsp and www.arabidopsis.org/tools/bulk/go/ (Table 1 and Table S5).

##### 8.2.6 Proof of concept examples

We argue that the power of our association approach relies on the fact that HPG1 lines resemble Near Isogenic Lines (NILs) produced by experimental crosses (29) (Fig. S2A). Similar to genome-wide association studies (GWA), power depends on many factors, namely the noise of phenotype under study, architecture of phenotypic trait, quality of genotyping, population structure, sample diversity, sample size, allele frequency, and recombination. On one hand, association analyses in NILs suffer from large linkage blocks, but confident results can be achieved due to accurate measurement of phenotypes, limited genetic differences between any two lines, and high quality genotypes. In common GWA studies such as in humans, there are multiple confounding effects. Among the confounders are (1) that any two samples differ in hundreds of thousands of SNPs, and (2) that historical and geographic stratification produce non-random correlations among those SNP differences. This considerably complicates the identification of phenotypic effects at specific genes, and power relies greatly on large sample sizes to achieve the sufficient number of recombination between markers.

To provide support for the non-synonymous SNP on chromosome 5, at position 6,508,329 in AT5G19330, we looked for pairs of lines that carry the ancestral and the derived allele, but that differ in few (or no other) SNPs in the genome. When considering all genic substitutions with a minimum allele frequency of 5% (Fig. S2A), we identified 20 pairs of lines differing only in the AT5G19330 SNP and another linked SNP (located on a different chromosome, association p-value > 0.4). The phenotypic differences in mean gravitropic score of these almost-identical pairs were significantly higher than phenotypic differences among all pairs of HPG1 lines, and genetically identical pairs attending to substitutions inside genes (Fig. S2A). Furthermore, this SNP was not in complete linkage with any other SNP hit (*r*^*2*^ < 0.5) (Fig. S2D). The same approach was used to examine the SNPs in AT1G54440 (Fig. S2E) and AT2G16580 (Fig. S2F), which represent an intermediate and a high LD example.

## SUPPLEMENTAL TABLES

**Table S1.** HPG1 sample information.

**Table S2.** Sample information for Col-0 mutation accumulation lines.

**Table S3.** Mutation rate estimates for different annotations in HPG1 and mutation accumulation lines.

**Table S4.** Description of phenotypic and climatic variables for association mapping analyses.

**Table S5.** SNP hits from association analyses and several descriptors.

**Data Appendix S1:**
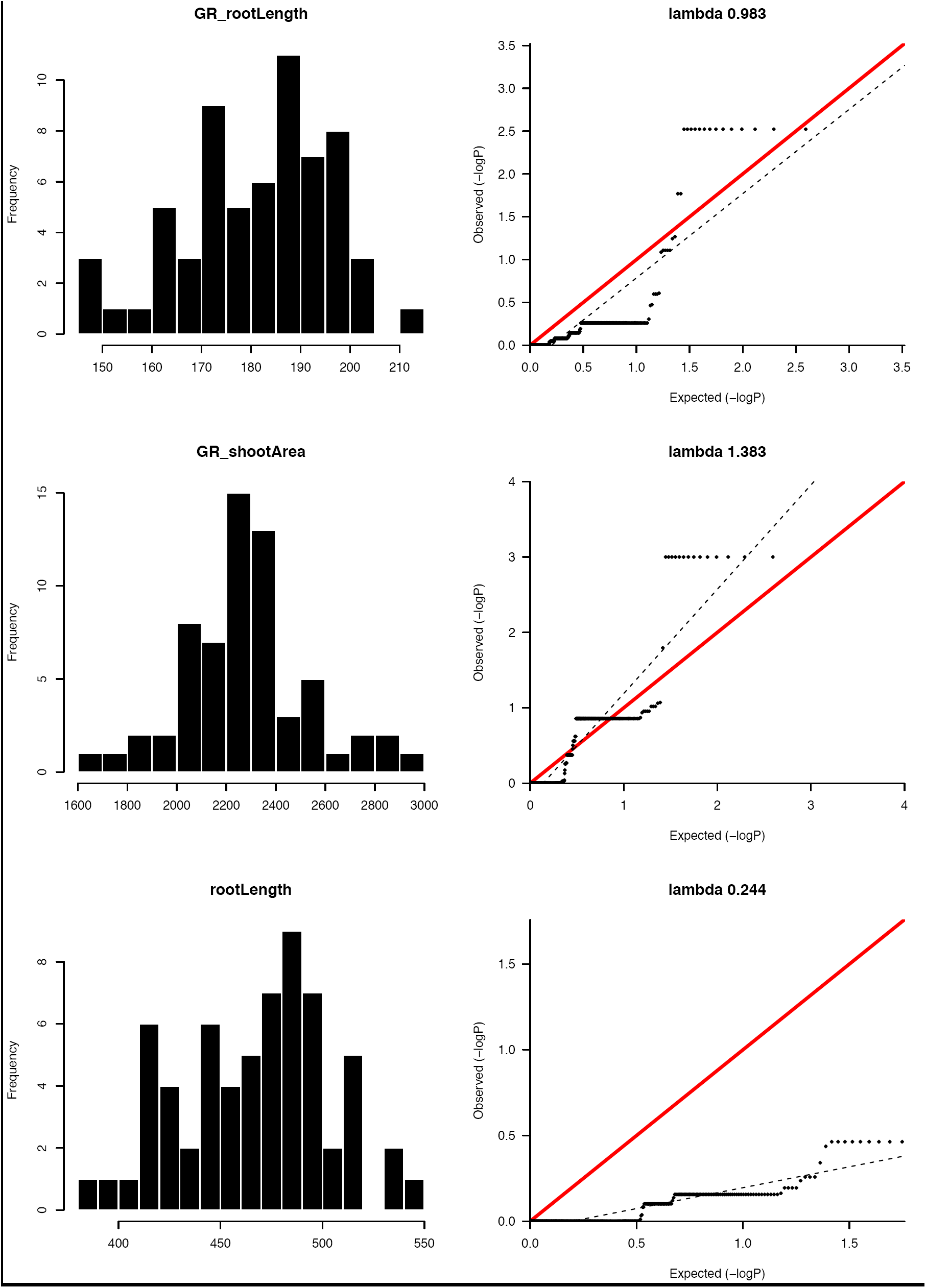
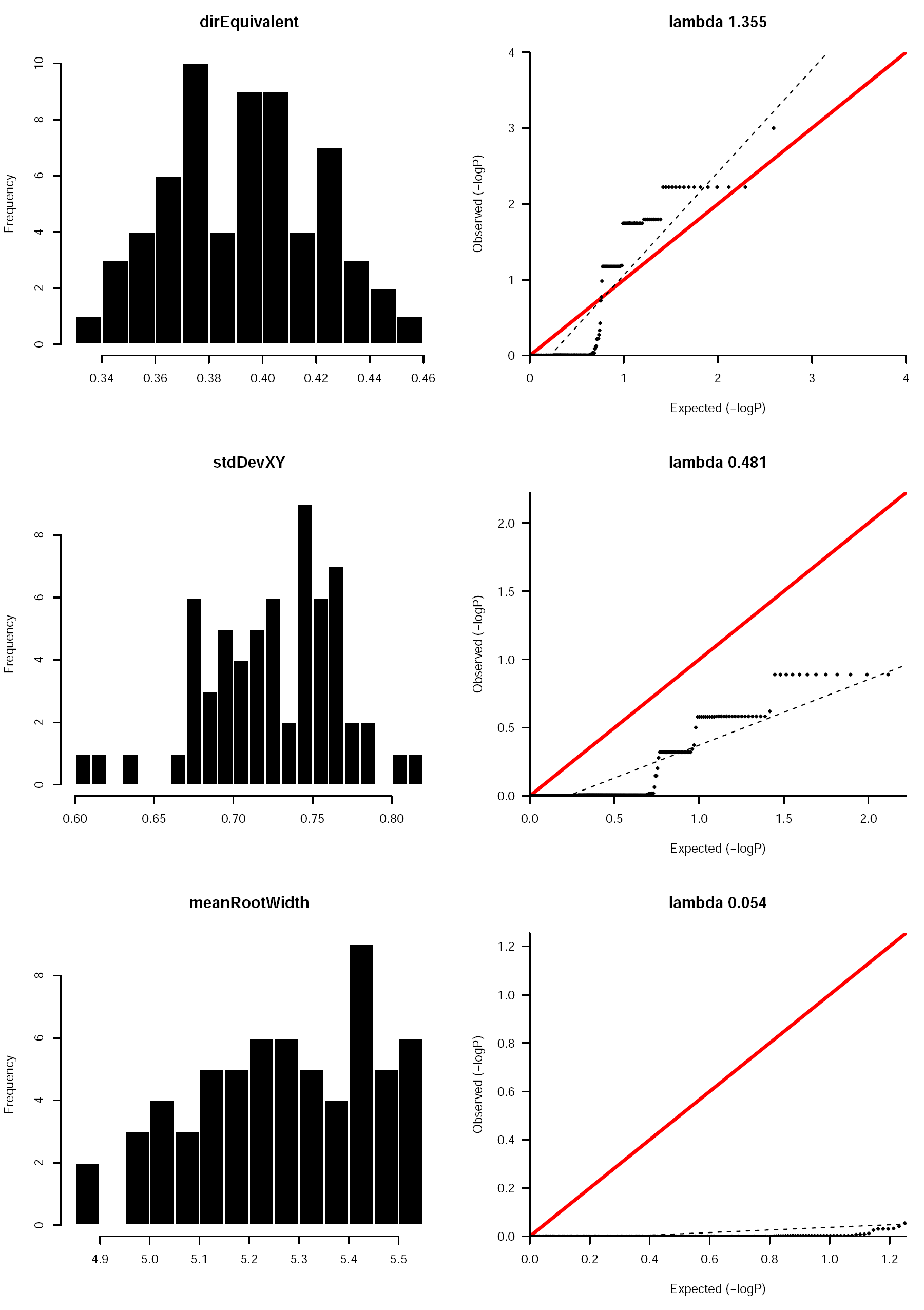
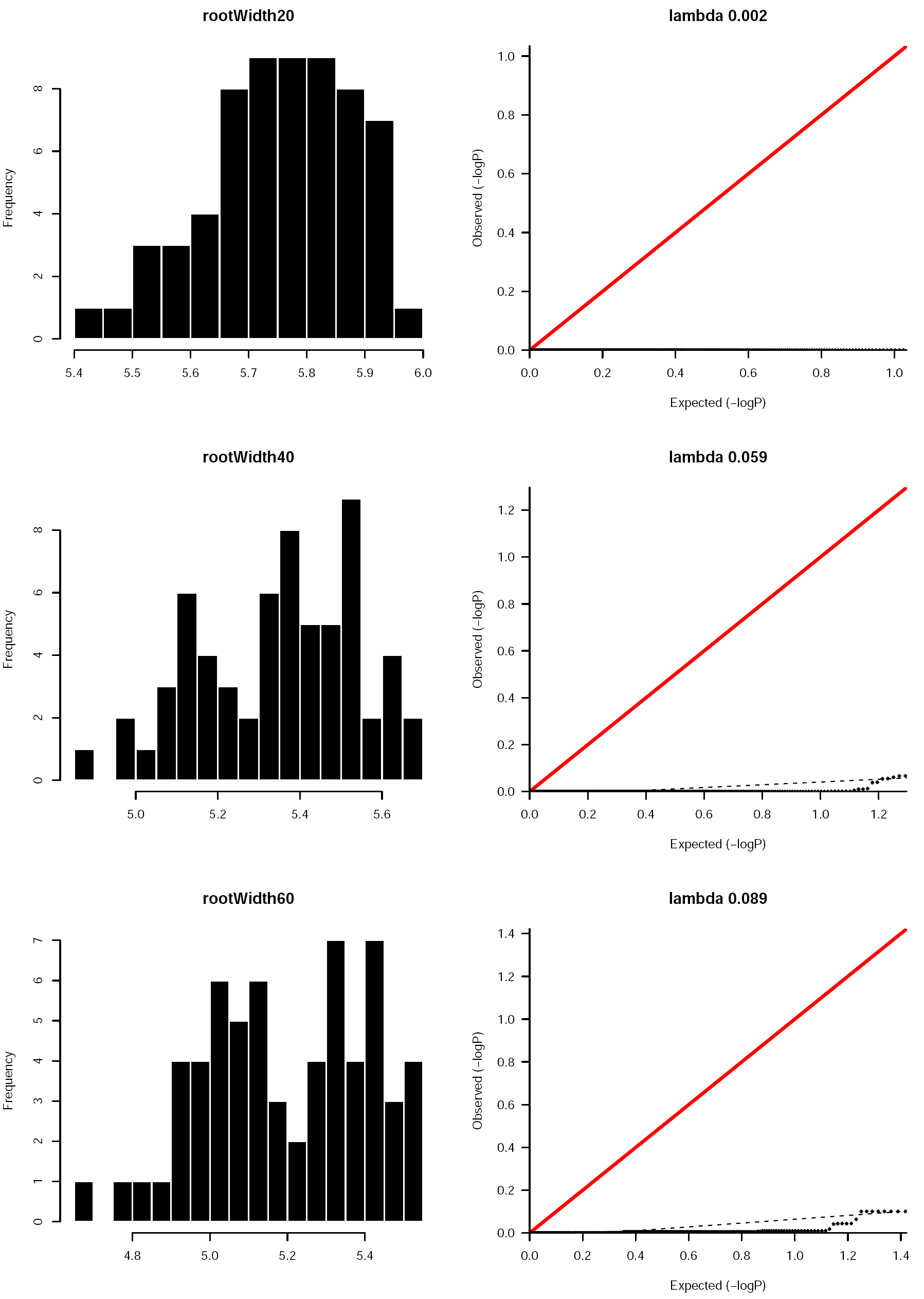
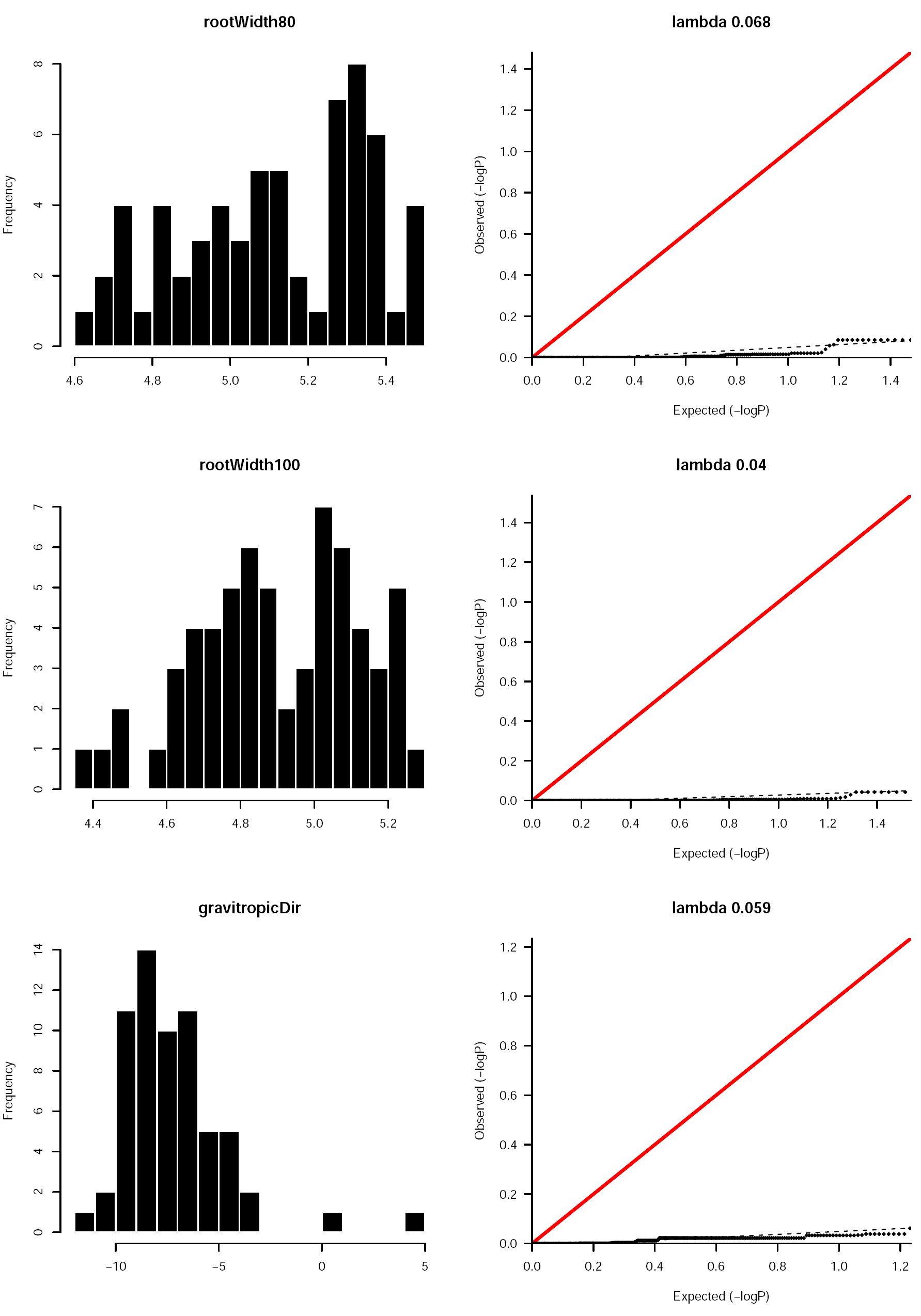
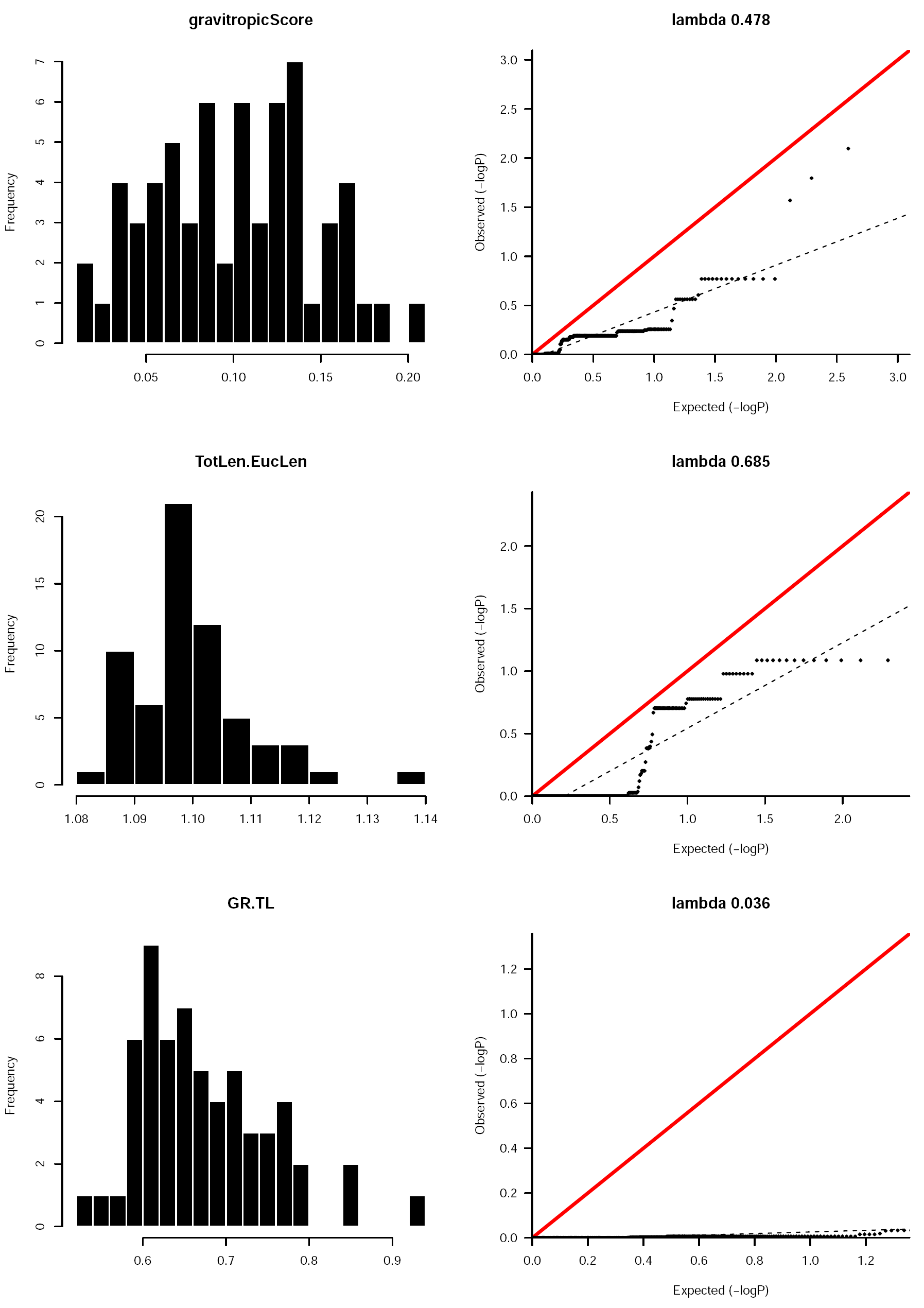
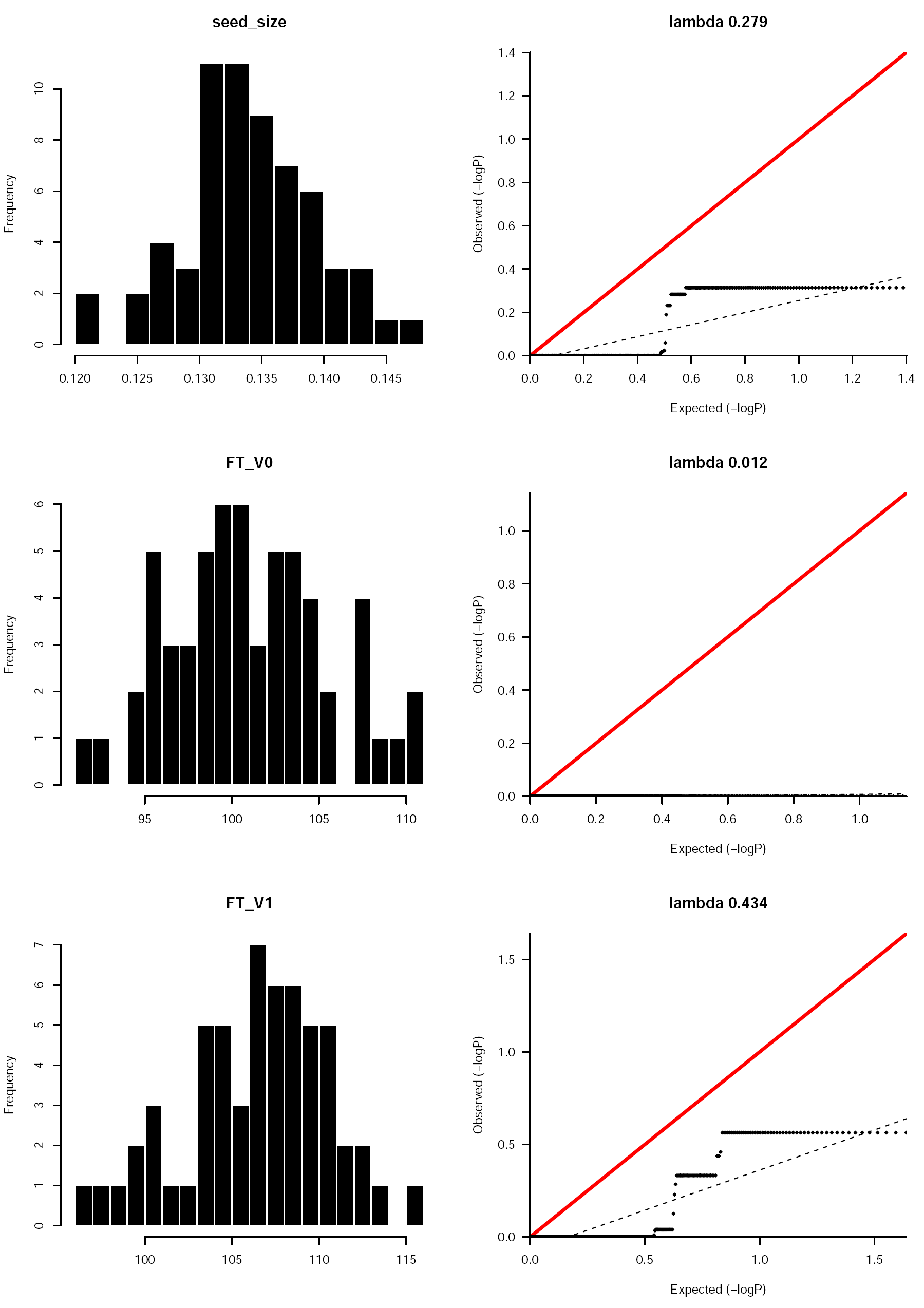
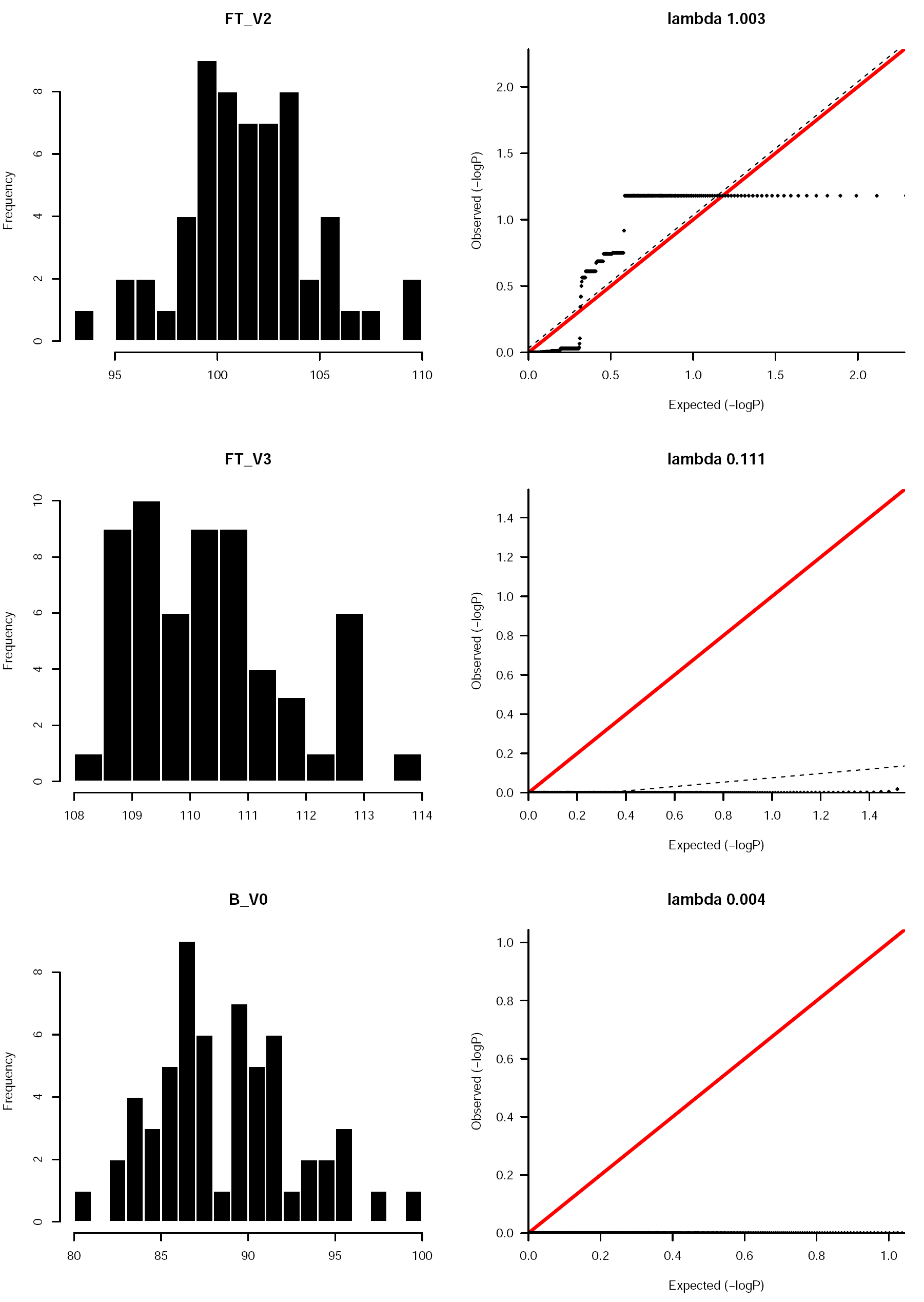
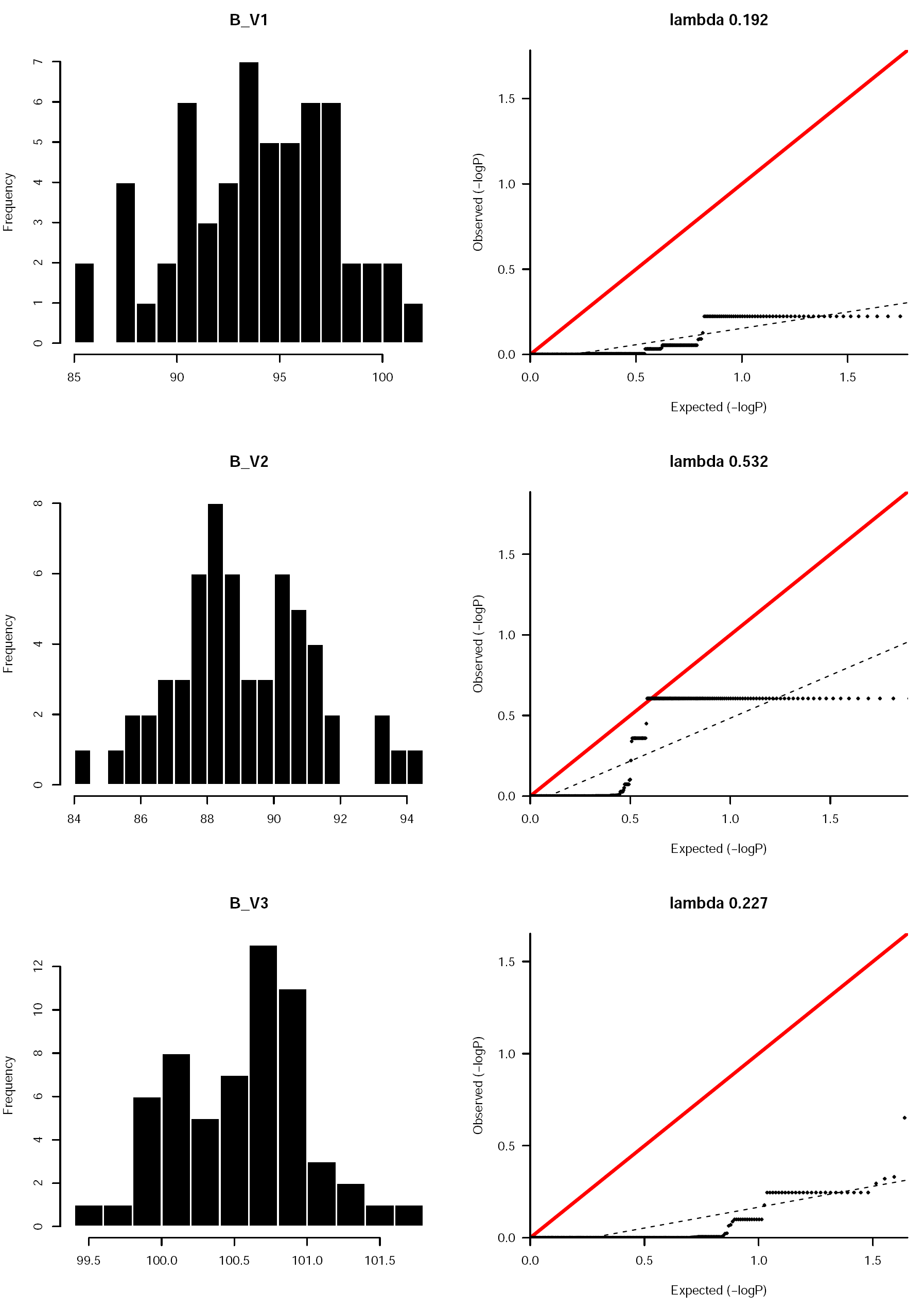
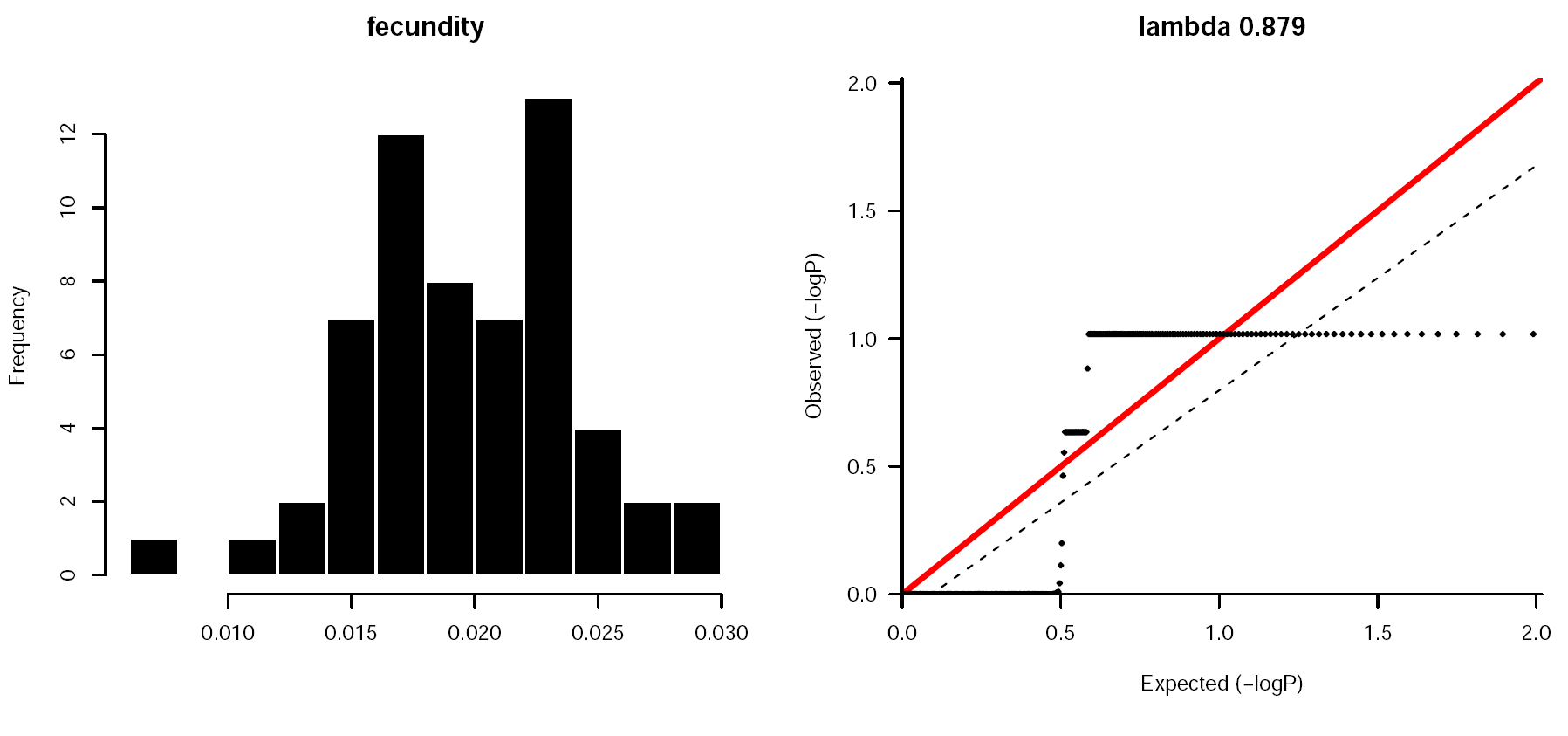
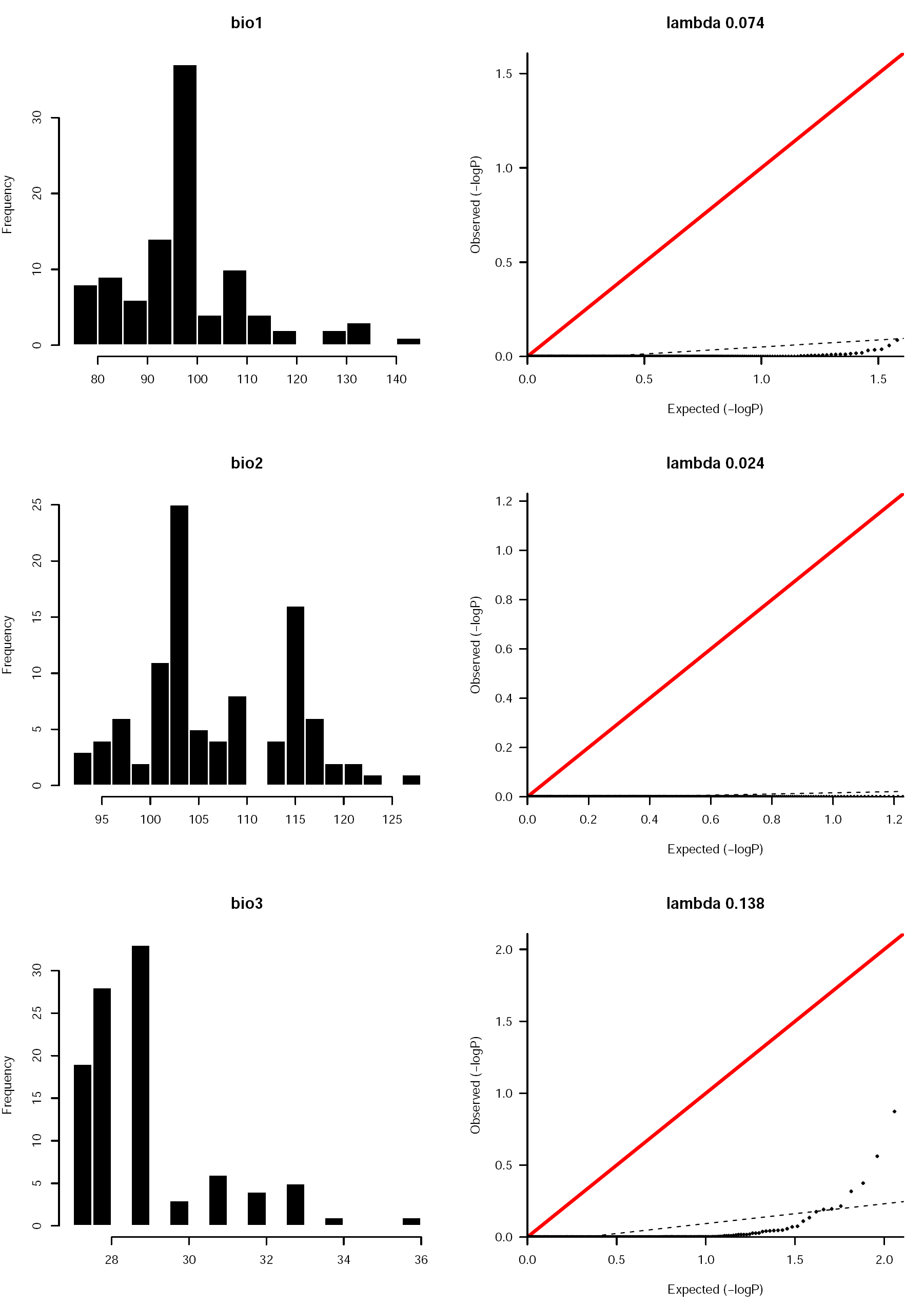
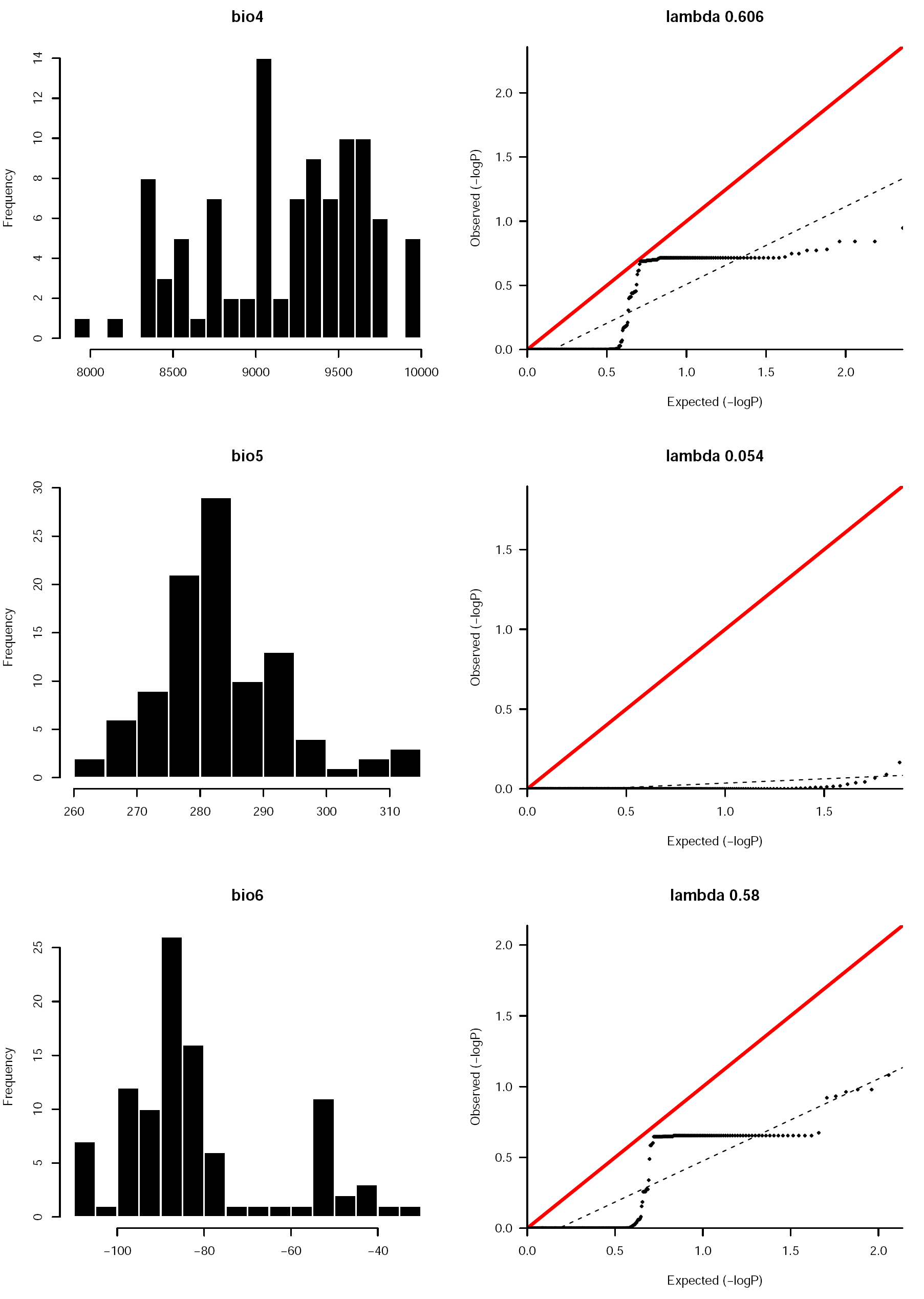
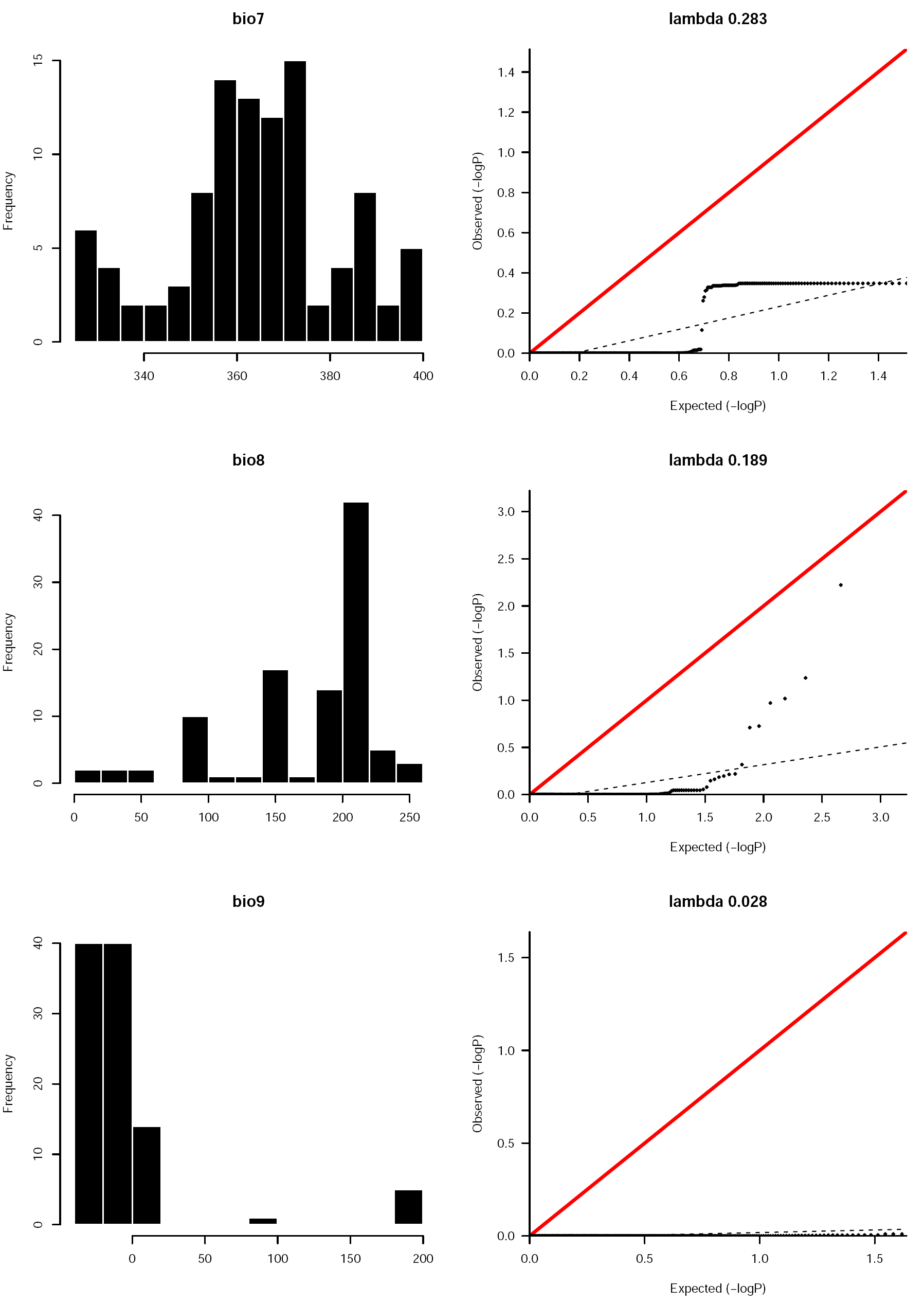
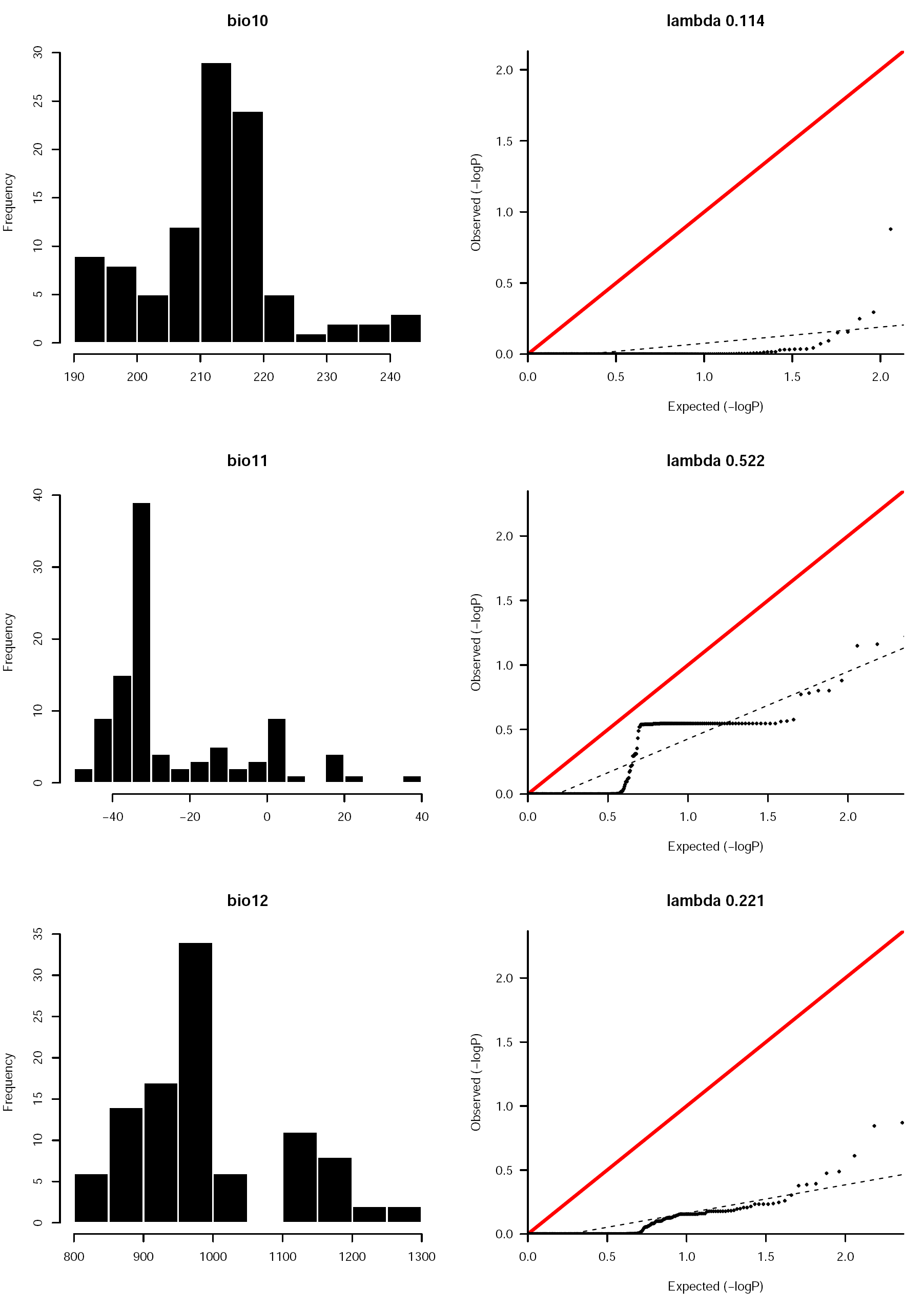
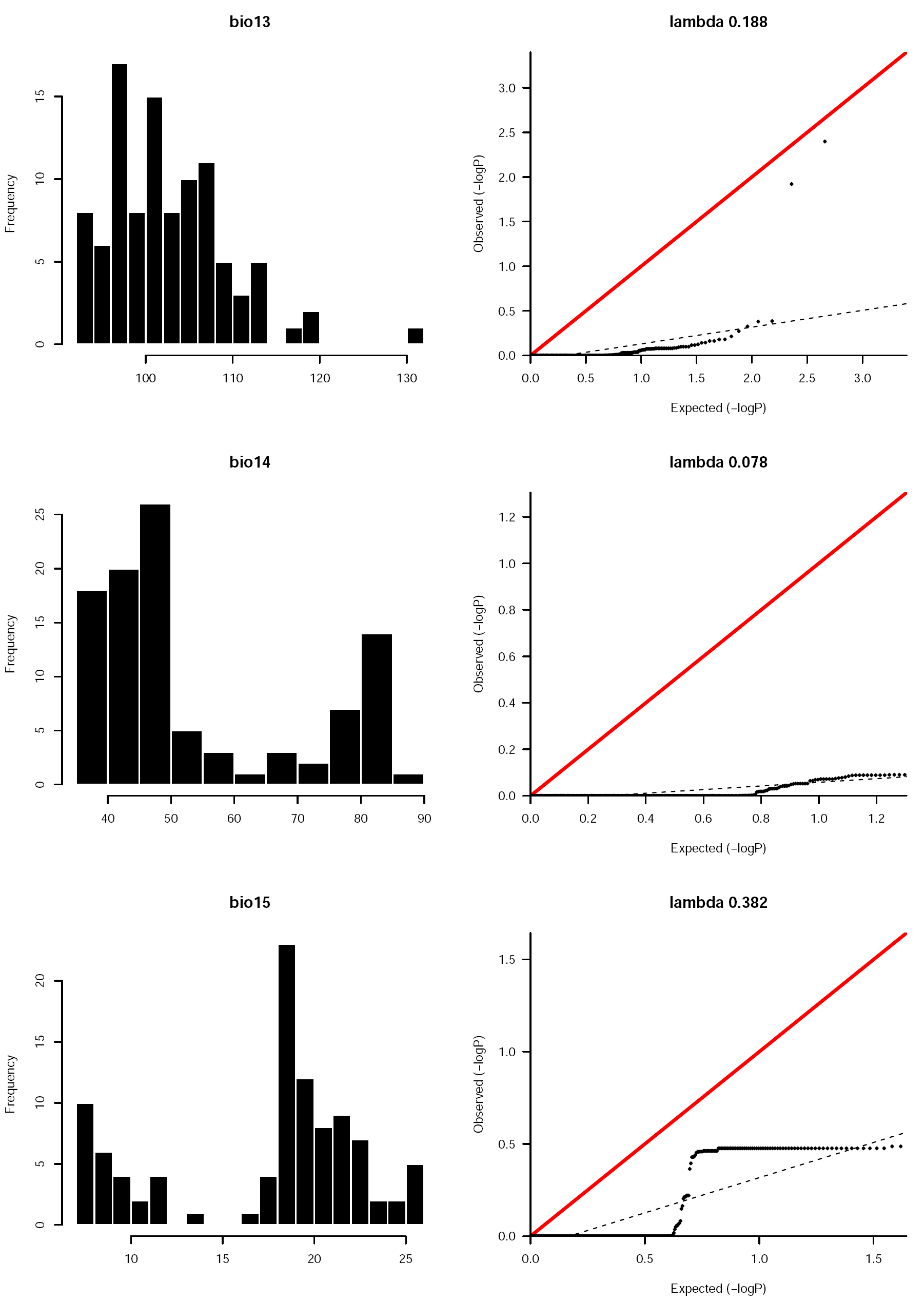
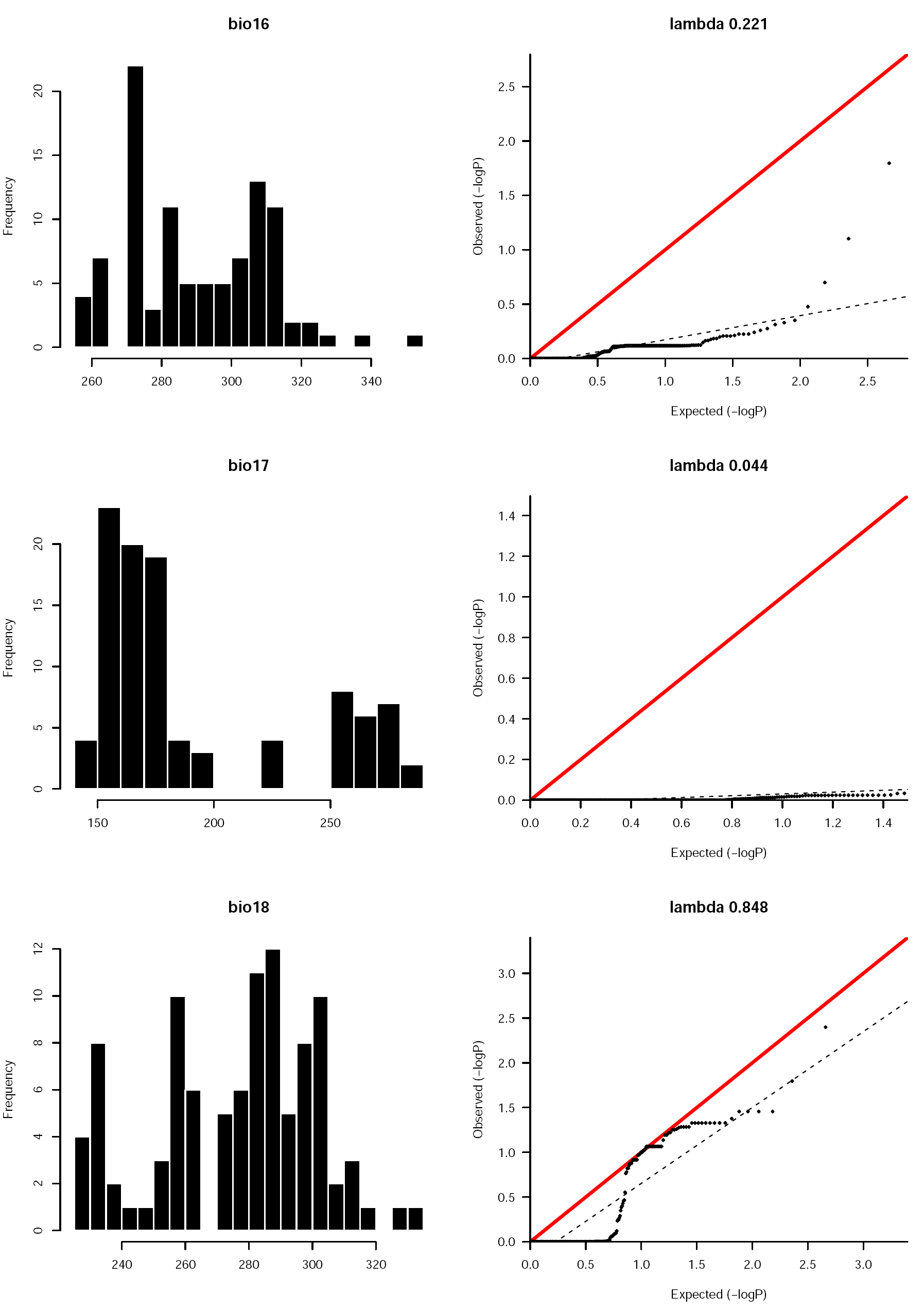
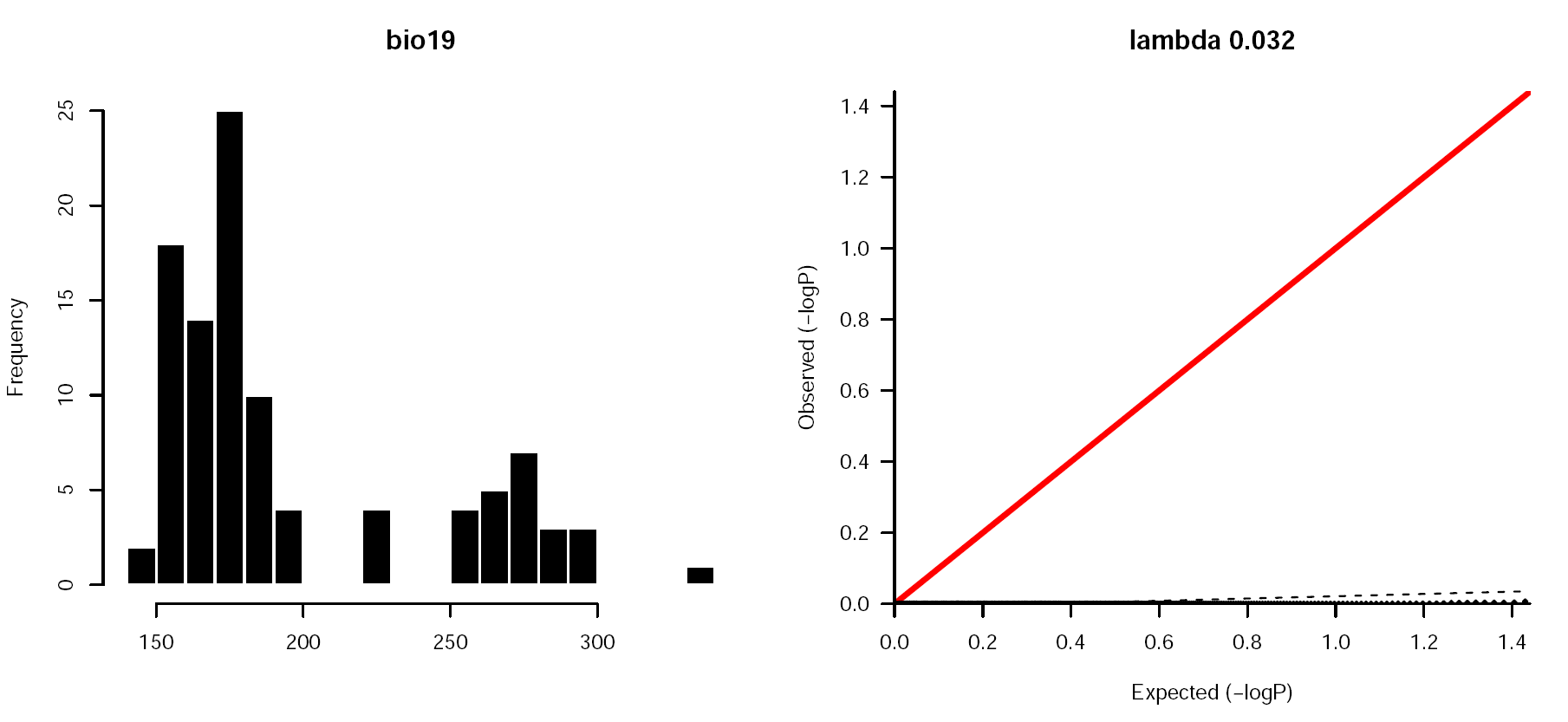
For each trait employed in association analyses, we report the histogram distribution and the QQ plot of p-values to ensure that no trait departs exaggeratedly from the normal distribution, and that no inflation of p-values is observed (when lambda ≤ 1, there is no inflation of false positives).

## SUPPLEMENTAL FIGURES

**Fig S1.**
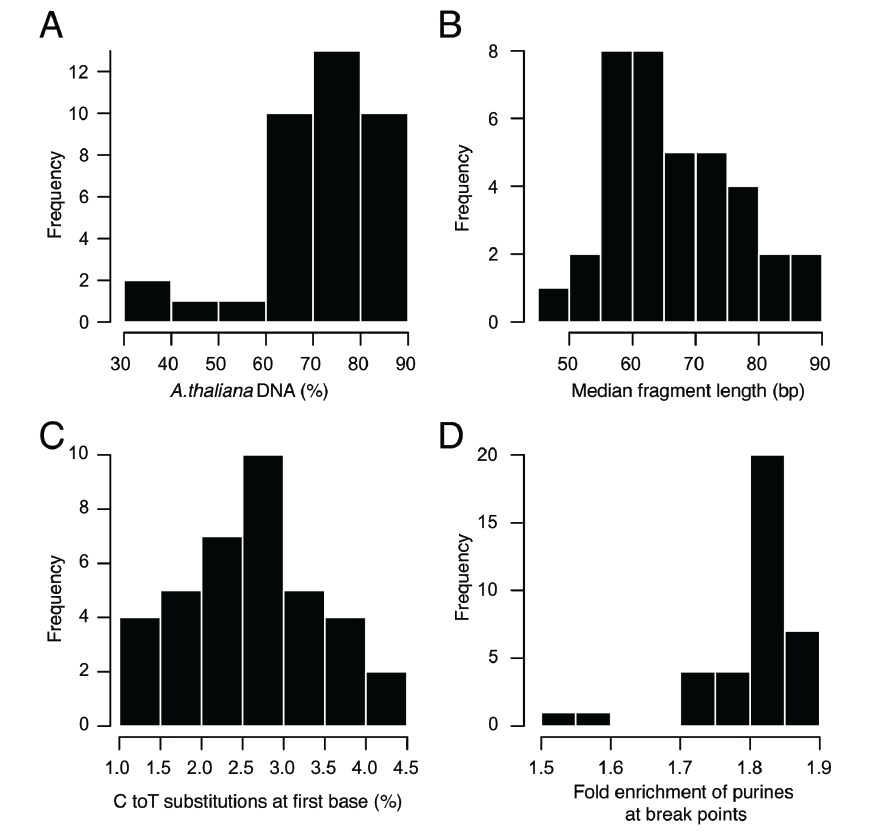
Ancient-DNA characteristics of unrepaired herbarium libraries. **(A)** Fraction of *A. thaliana* DNA in sample. **(B)** Median length of merged reads. **(C)** Fraction of cytosine to thymine (C-to-T) substitutions at first base (5’ end). **(D)** Relative enrichment of purines (adenine and guanine) at 5’ end breaking points. Position-1 is compared with position-5 (negative numbers indicate genomic context before upstream reads’ 5’ end).

**Fig S2.**
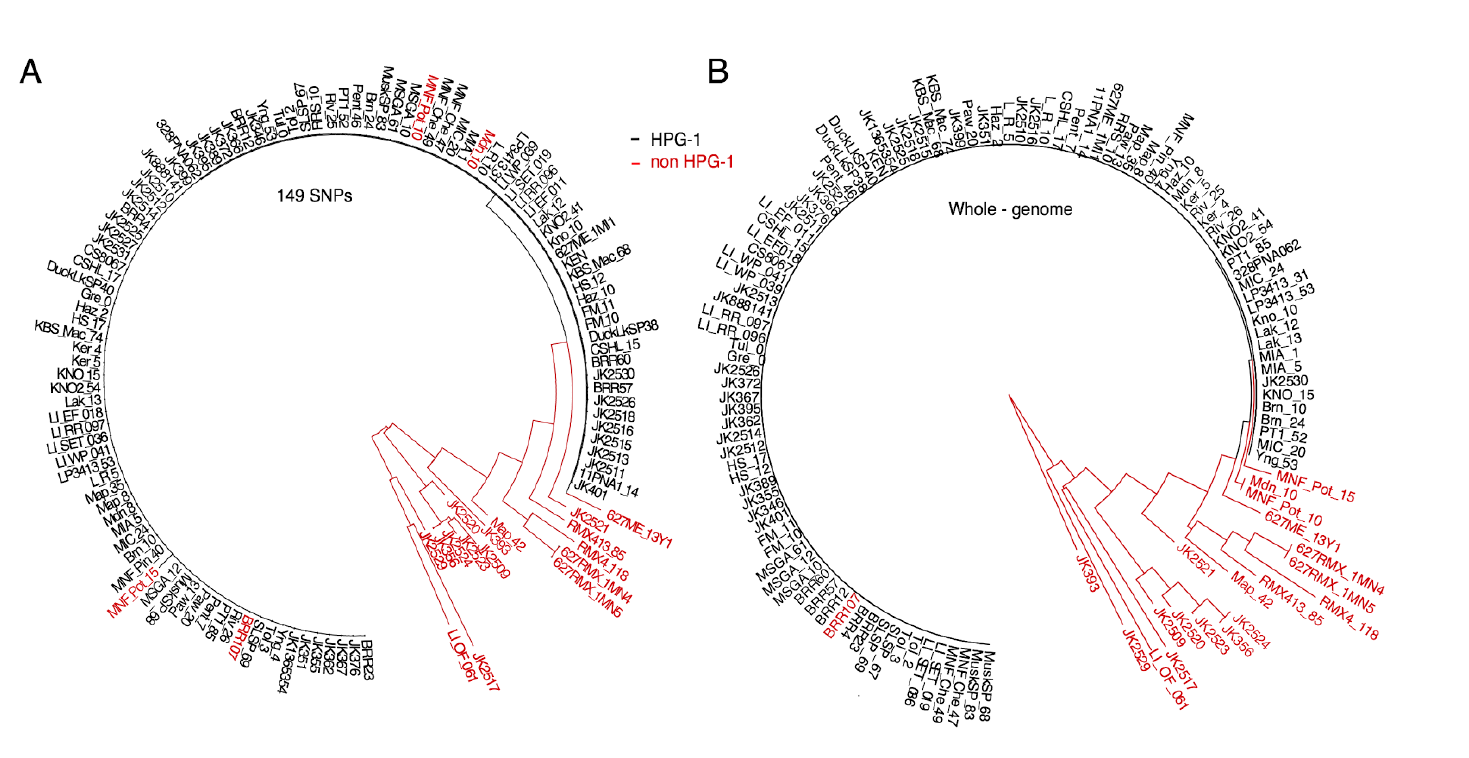
Separation between HPG1 and other North American lineages. **(A)** Neighbor-joining tree built using Illumina-based SNP calls at the 149 genotyping markers originally used to identify HPG1 candidates. HPG1 accessions are shown in black, whereas other North American lineages are depicted in red (see explanation below for four HPG1-like accessions). **(B)** Neighbor-joining tree based on genome-wide SNPs. Accessions colored as in (A). Note that three accessions originally classified as HPG1 based on 149 SNPs (A) are placed outside this clade. A further accession (BRR7) within the HPG1 main branch was a recombinant removed from the analysis.

**Fig S3.**
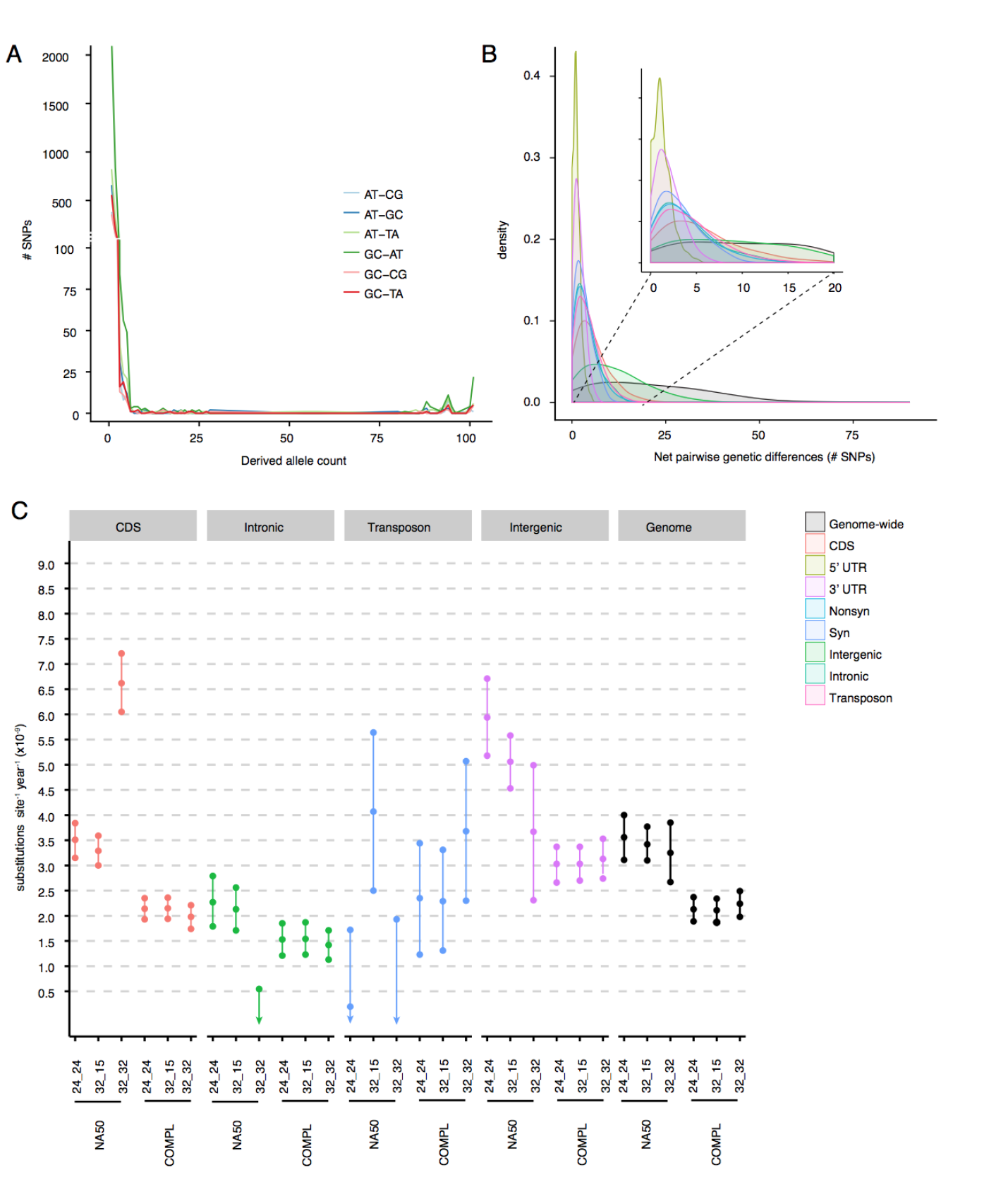
Substitution spectrum and rates. **(A)** Site frequency spectrum for all transitions and transversions. **(B)** Distributions of “net” pairwise genetic distances between historic and modern samples used to calculate mutation rates per genomic annotation (from quality 32_15 and complete information per site). UTRs were excluded because of the small number of SNPs. **(C)** Mutation rates calculated for different genomic annotations and quality thresholds (32_32, 32_15, 24_24) and missing values (NA50: maximum 50% missing data per SNP; COMPL: missing data 0%). Mean and 95% confidence intervals are shown.

**Fig S4.**
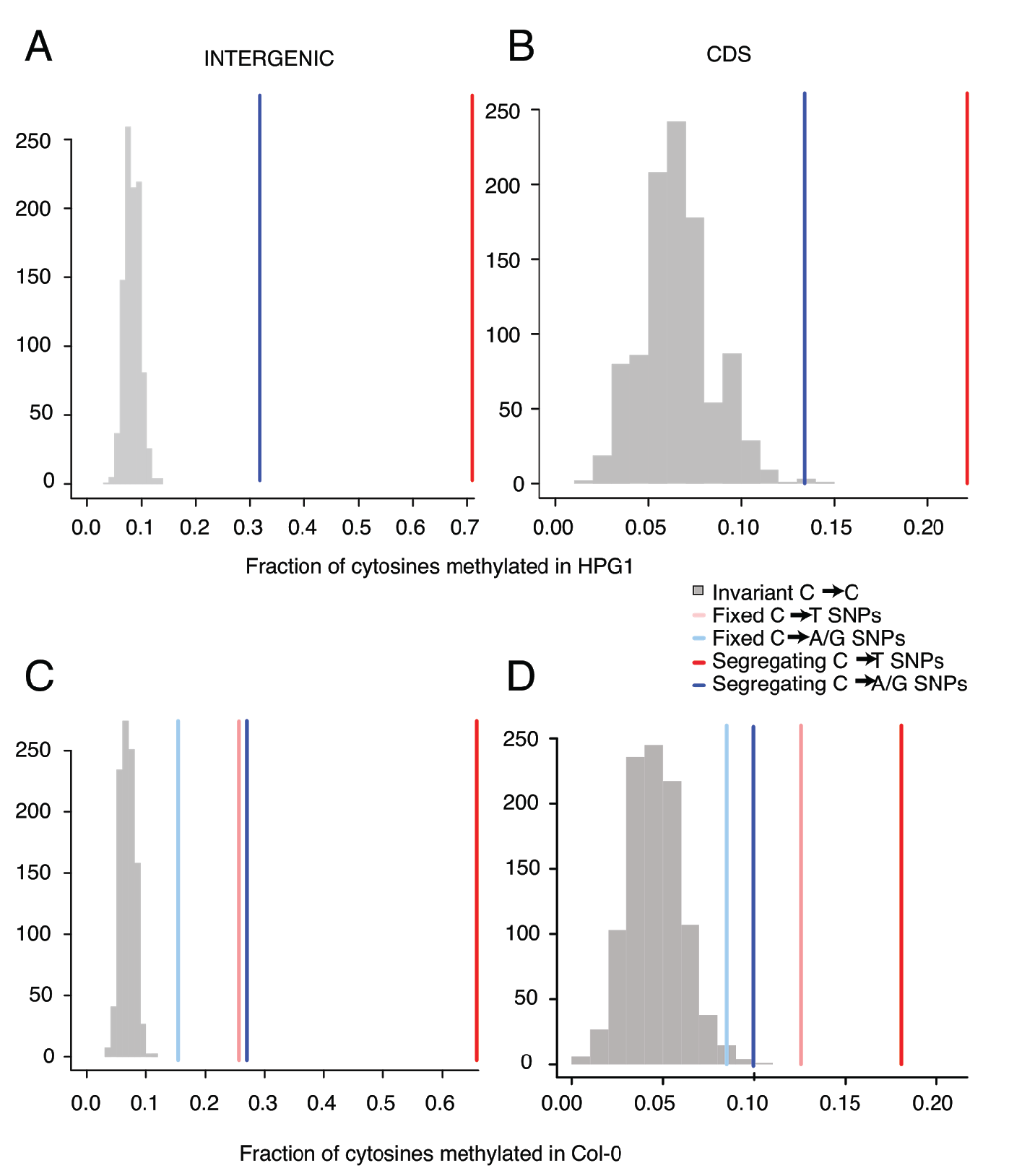
Relationship between methylation and substitutions. **(A, B)** Fraction of methylation of cytosines in HPG1 pseudo-reference(7) at intergenic (A) or coding regions (B). **(C, D)** Fraction of methylation of cytosines in Col-0 reference genome(5) at intergenic (C) or coding regions (D). In each of the four comparisons, a grey histogram represents distribution of methylation of 1,000 random sets of invariant cytosines. Lines represent average methylation degree at those sites in HPG1 that changed from cytosine to thymine (red). We differentiate those substitutions that are shared-fixed-across all individuals (light red) or whose allele are present at an intermediate-segregating-frequency (dark red). Likewise, average methylation is shown for sites that changed from cytosine to adenine (blue) that that are fixed (light blue) or segregating (dark blue). The fact that the average methylation is higher in new substitutions than in invariant positions supports a connection between methylation and mutability of sites.

**Fig S5.**
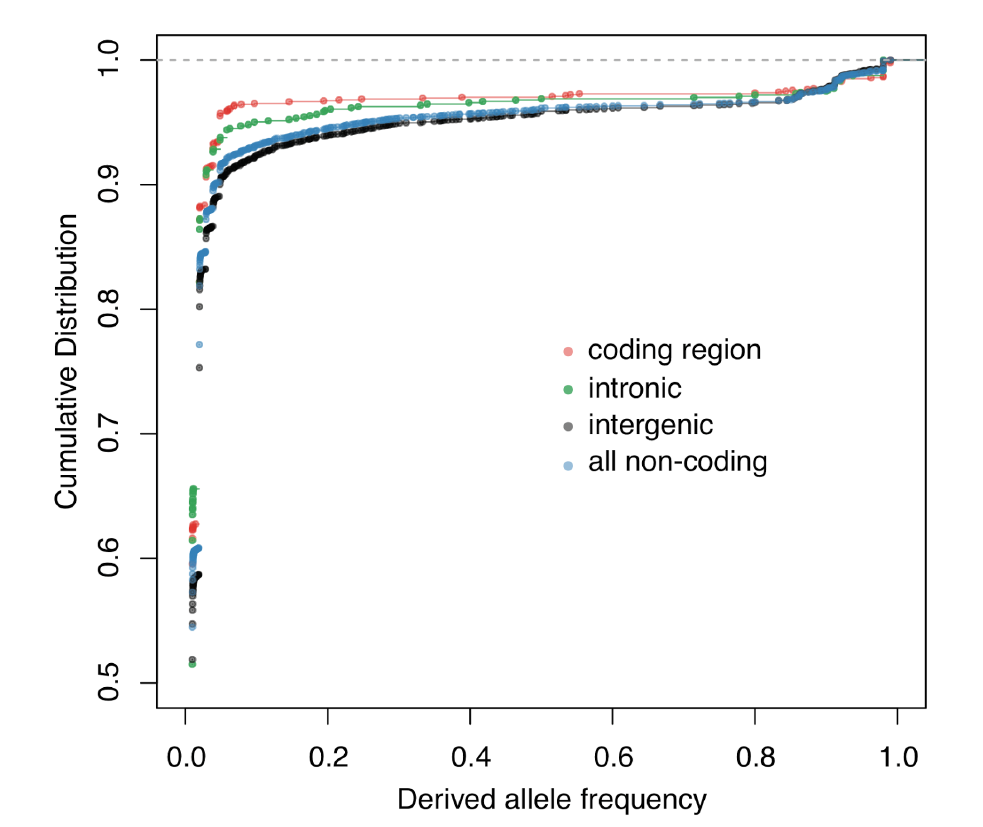
Comparison of Site Frequency Spectra across genomic annotations. Cumulative empirical distribution, at different genomic annotations, of the unfolded Site Frequency Spectrum of SNPs oriented based on the order of appearance of alleles in the herbarium genomes.

**Fig S5.**
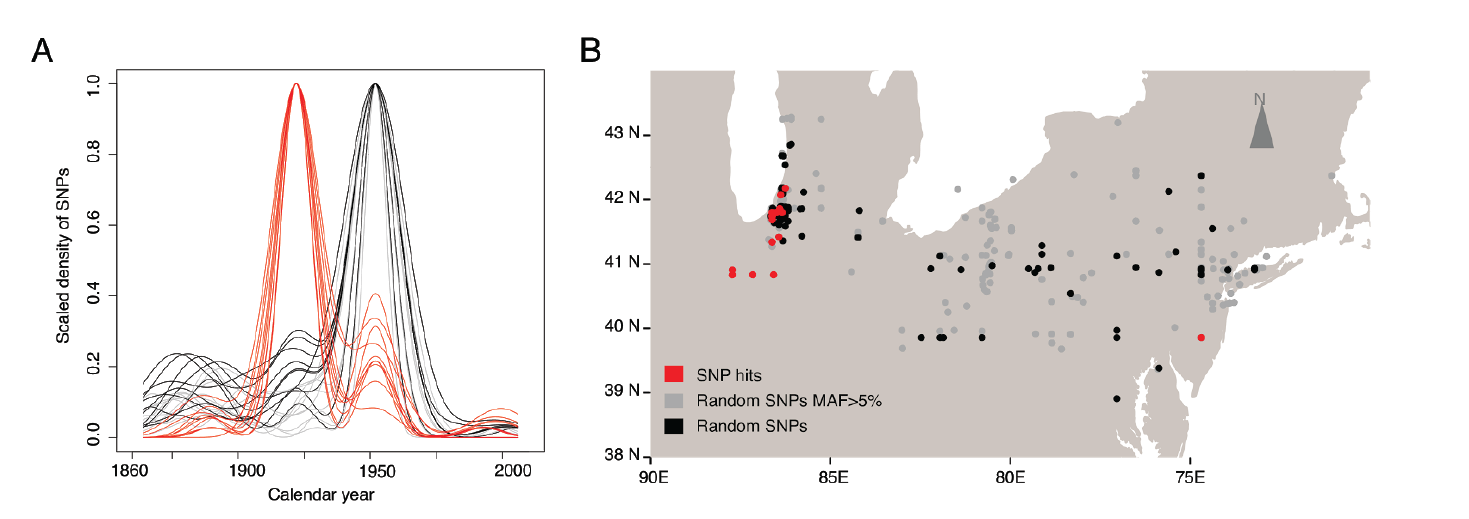
Spatial and temporal emergence of root-associated mutations. **(A)** Age distribution of derived SNPs with a significant trait association (the herbarium sample in which they were first recorded) (red), compared with genome-wide SNPs with at least 5% minor allele frequency (grey), or without frequency cutoff (black). **(B)** Spatial centroid of all samples carrying a derived allele. Since it is an average location, centroids can be in a body of water. Ten random draws of 50 SNPs for each category were used to produce the density lines in (A) and points in (B).

**Fig S6.**
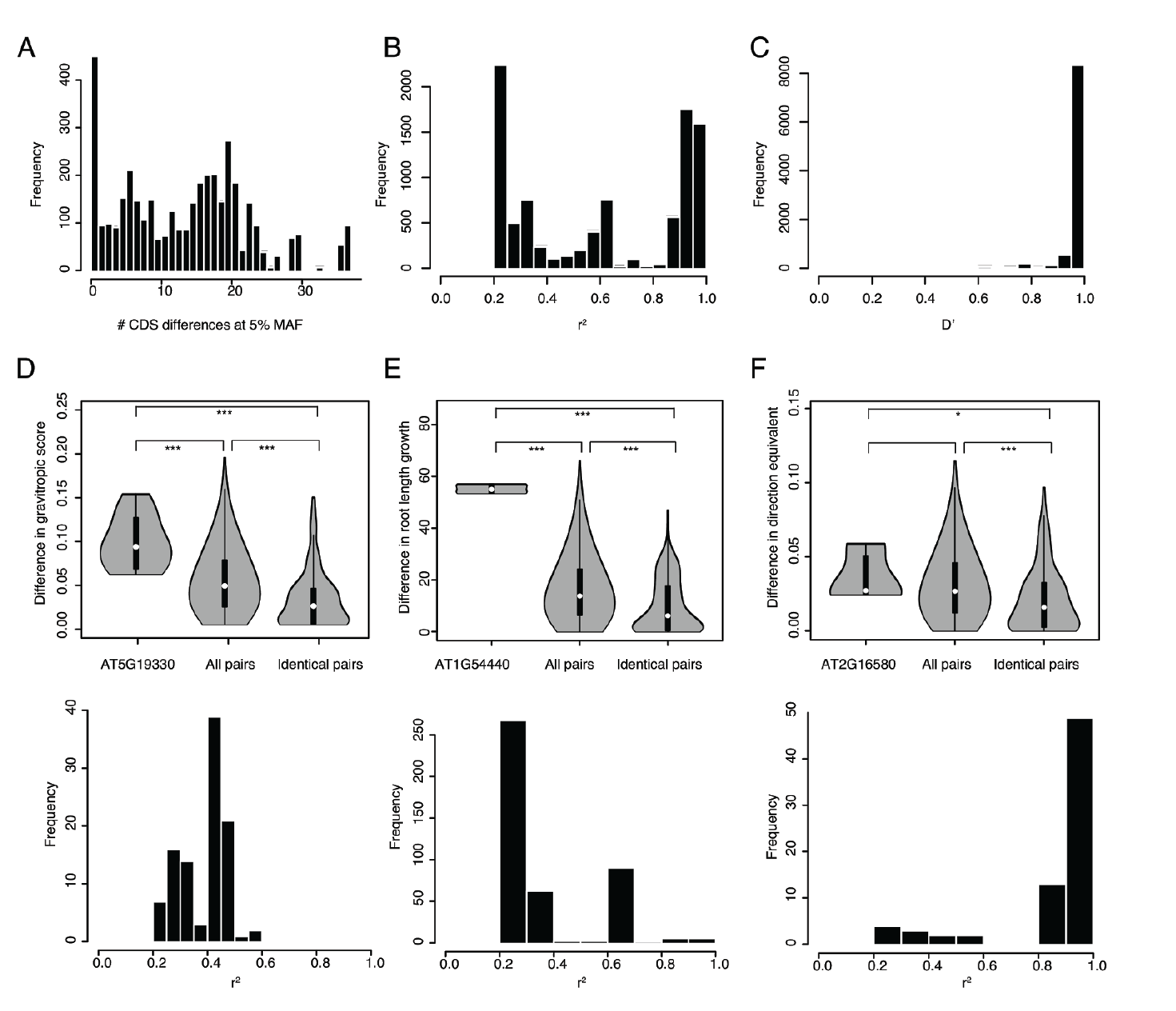
Linkage disequilibrium of significant SNPs. **(A-F)** Linkage disequilibrium between SNPs with significant trait associations. Histogram of genetic distances **(A)** between samples when evaluating only coding regions at 5% minimum allele frequency. Linkage disequilibrium between SNP hits measured as *r*^2^ **(B)** and *D*’ **(C)**. Three significant SNPs were further studied to exemplify the power of association analyses with HPG1. For each, phenotypic differences between accessions that differ in the focal SNP and that are otherwise virtually genetically identical are compared both with all pairs of accessions and with pairs of accessions completely identical for coding regions. Below each violin plot is the histogram of linkage disequilibrium of the focal SNP with all other SNP hits. The three focal SNPs evaluated are located in AT5G19330 **(D)**, AT1G54440 **(E)** and AT2G16580 **(F)**.

**Table S1.** Sample information. (Abbreviation H* indicates herbarium samples that cluster with the modern HPG1 clade rather than the historic HPG1 clade in Fig. 3., highlighted as a star in the map from Fig. 1. Abbreviations of herbarium collections or seed sources: UCONN = University of Connecticut Herbarium; CFM = Chicago Field Museum; NY = New York Botanical Garden; ABRC = Arabidopsis Biological Resources Center; OSU = Ohio State University.)

| Accession | Latitude (°N) | Longitude (°E) | State | Date collected | Alternative name | Collector/ Herbarium | Average coverage (x) | Number of covered positions (≥3x)<br>(mapped against HPG1 reference) | Number of covered positions (≥3x)<br>(mapped against Col_0 reference) | SNPs vs HPG1 reference | Belongs to HPG1 | Modern/ Herbarium | Column number in the available genome matrix |
| --- | --- | --- | --- | --- | --- | --- | --- | --- | --- | --- | --- | --- | --- |
| JK399 | 38.7155 | -75.635591 | DE | 1863 | 888124 | NY | 9 | 105,053,631 | 99,889,683 | 142 | yes | H | 101 |
| JK366 | 43.1921 | -77.0102 | NY | 1866 | 888144 | NY | 6.8 | 100,379,839 | 95,118,236 | 123 | yes | H | 94 |
| JK395 | 38.9068 | -77.036667 | DC | 1877 | 888134 | NY | 10.3 | 103,620,791 | 98,888,406 | 167 | yes | H | 100 |
| JK888141 | 40.732007 | -74.068455 | NJ | 1879 | 888141 | NY | 42 | 107,211,409 | 102,634,255 | 161 | yes | H | 103 |
| JK389 | 38.9068 | -77.036667 | DC | 1888 | 1365363 | NY | 9.9 | 106,042,465 | 100,826,958 | 151 | yes | H | 98 |
| JK362 | 38.9068 | -77.036667 | DC | 1889 | 1365364 | NY | 8.8 | 103,997,716 | 98,876,320 | 153 | yes | H | 93 |
| JK367 | 40.9249 | -74.0755 | NJ | 1890 | 1365344 | NY | 16.7 | 107,236,732 | 102,176,782 | 181 | yes | H | 95 |
| JK372 | 41.1222 | -74.3569 | NJ | 1890 | 1365332 | NY | 14.8 | 106,285,178 | 101,480,369 | 163 | yes | H | 96 |
| JK1365354 | 38.8782 | -77.09048 | VA | 1891 | 1365354 | NY | 36.4 | 106,718,326 | 102,458,166 | 169 | yes | H | 88 |
| JK376 | 39.97 | -83.01 | NY | 1891 | 1365337 | NY | 12.3 | 105,962,154 | 100,840,125 | 145 | yes | H | 97 |
| JK351 | 41.15 | -73.766667 | NY | 1894 | 1365333 | NY | 16.1 | 106,531,302 | 101,841,156 | 153 | yes | H | 90 |
| JK355 | 35.99 | -83.94 | TE | 1896 | 1365374 | NY | 14.3 | 106,391,637 | 101,455,311 | 192 | yes | H | 91 |
| JK356 | n/a | n/a | GA | 1897 | 1365375 | NY | 5.3 | 90,426,010 | 89,296,191 | n/a | no | H | 92 |
| JK393 | n/a | n/a | NC | 1897 | 1365370 | NY | 30.4 | 102,894,430 | 101,298,068 | n/a | no | H | 99 |
| JK346 | 40.643136 | -111.95177 | UT | 1903 | 102365 | NY | 29.1 | 107,223,283 | 102,450,446 | 222 | yes | H | 89 |
| JK2525 | 41.224343 | -73.06021 | CT | 1904 | 79391 | UNCONN | 12.5 | 105,025,845 | n/a | 138 | yes | H | 118 |
| JK2529 | n/a | n/a | OH | 1904 | 176849 | CFM | 11.4 | 100,620,441 | n/a | n/a | no | H | 121 |
| JK401 | 40.643136 | -111.95177 | UT | 1904 | 102364 | NY | 10.4 | 99,572,736 | 94,661,828 | 216 | yes | H | 102 |
| JK2513 | 41.102121 | -81.560547 | OH | 1911 | 25 | OSU | 18.2 | 106,309,854 | n/a | 176 | yes | H | 108 |
| JK2509 | n/a | n/a | CT | 1917 | 11 | OSU | 15.1 | 102,169,546 | n/a | n/a | no | H | 104 |
| JK2530 | 41.482862 | -86.822602 | IN | 1922 | 531679 | CFM | 22.2 | 107,043,540 | n/a | 161 | yes | H | 122 |
| JK2526 | 41.666667 | -73.508455 | CT | 1929 | 79409 | UNCONN | 16.3 | 107,026,827 | n/a | 161 | yes | H | 119 |
| JK2515 | 41.137296 | -81.863779 | OH | 1930 | 30 | OSU | 21.3 | 106,893,416 | n/a | 193 | yes | H | 110 |
| JK2511 | 41.721618 | -81.243317 | OH | 1934 | 14 | OSU | 5.6 | 95,822,372 | n/a | 109 | yes | H | 106 |
| JK2523 | n/a | n/a | OH | 1940 | 25707 | UNC | 13.1 | 101,421,749 | n/a | n/a | no | H | 116 |
| JK2520 | n/a | n/a | OH | 1945 | 54051 | UNC | 20.3 | 102,831,697 | n/a | n/a | no | H | 114 |
| JK2524 | 39.856783 | -74.686954 | NJ | 1952 | 63978 | UNC | 13.8 | 100,778,282 | n/a | n/a | no | H | 117 |
| JK2512 | 39.95607 | -81.953309 | OH | 1956 | 21 | OSU | 16.7 | 106,801,844 | n/a | 189 | yes | H | 107 |
| JK2514 | 39.95607 | -81.953309 | OH | 1969 | 27 | OSU | 28.4 | 107,044,415 | n/a | 219 | yes | H | 109 |
| JK2517 | n/a | n/a | OH | 1981 | 34 | OSU | 21.7 | 102,643,436 | n/a | n/a | no | H | 112 |
| JK2521 | n/a | n/a | OH | 1992 | 565960 | UNC | 2.9 | 62,673,938 | n/a | n/a | no | H | 115 |
| JK2518 | 41.867643 | -80.789021 | OH | 1993 | 40 | OSU | 14.8 | 106,578,197 | n/a | 177 | yes | H | 113 |
| JK2531 | 39.856783 | -74.686954 | NJ | 1952 | 1507461 | CFM | 15.1 | 106,158,181 | n/a | 177 | yes | H* | 123 |
| JK2510 | 39.688861 | -82.993218 | OH | 1930 | 13 | OSU | 21 | 106,305,970 | n/a | 178 | yes | H* | 105 |
| JK2527 | 41.509059 | -72.543694 | CT | 1975 | 79389 | UNCONN | 8.3 | 104,089,205 | n/a | 200 | yes | H* | 120 |
| JK2516 | 39.500862 | -82.472413 | OH | 1980 | 32 | OSU | 18.1 | 106,464,569 | n/a | 198 | yes | H* | 111 |
| CSHL_15 | 40.8585 | -73.4675 | NY | 1993 | CSHL-15 | ABRC | 39.3 | 108,189,771 | 105,955,885 | 243 | yes | M | 16 |
| CSHL_17 | 40.8585 | -73.4675 | NY | 1993 | CSHL-17 | ABRC | 41.5 | 108,194,960 | 105,982,511 | 240 | yes | M | 17 |
| FM_10 | 42.4489 | -76.5072 | NY | 1993 | FM-10 | ABRC | 44.6 | 108,203,215 | 106,052,866 | 269 | yes | M | 20 |
| FM_11 | 42.4489 | -76.5072 | NY | 1993 | FM-11 | ABRC | 44.4 | 108,214,008 | 106,040,276 | 288 | yes | M | 21 |
| HS_12 | 42.373 | -71.0627 | MA | 1993 | HS-12 | ABRC | 48.8 | 108,230,030 | 106,124,249 | 251 | yes | M | 25 |
| HS_17 | 42.373 | -71.0627 | MA | 1993 | HS-17 | ABRC | 55.3 | 108,242,062 | 106,155,362 | 254 | yes | M | 26 |
| Kno_10 | 41.2816 | -86.621 | IN | 1993 | Kno-10 | ABRC | 39.4 | 108,198,601 | 105,985,288 | 226 | yes | M | 32 |
| KNO_15 | 41.2816 | -86.621 | IN | 1993 | KNO-15 | ABRC | 43.6 | 108,219,683 | 106,069,077 | 231 | yes | M | 33 |
| Gre_0 | 43.178 | -85.2532 | MI | 1995 | Gre-0 | ABRC | 44.6 | 108,209,345 | 106,032,827 | 207 | yes | M | 22 |
| Tul_0 | 43.2708 | -85.2563 | MI | 1995 | CS6877 | ABRC | 31.2 | 108,140,393 | 105,806,418 | 221 | yes | M | 85 |
| CS8067 | 41.3599 | -122.755 | CA | 1996 | Buckhorn Pas | ABRC | 66.4 | 108,260,489 | 106,243,277 | 294 | yes | M | 15 |
| Tol_2 | 41.6639 | -83.5553 | OH | 1996 | CS8022 | ABRC | 61 | 108,241,333 | 106,194,209 | 238 | yes | M | 83 |
| Tol_3 | 41.6639 | -83.5553 | OH | 1996 | CS8023 | ABRC | 40.2 | 108,184,749 | 105,953,559 | 232 | yes | M | 84 |
| MIA_1 | 41.7976 | -86.6691 | MI | 1999 | MIA-1 | ABRC | 73.1 | 108,279,881 | 106,291,612 | 234 | yes | M | 56 |
| MIA_5 | 41.7976 | -86.6691 | MI | 1999 | MIA-5 | ABRC | 62.9 | 108,263,557 | 106,250,560 | 235 | yes | M | 57 |
| MIC_20 | 41.8266 | -86.4366 | MI | 1999 | MIC-20 | ABRC | 39.9 | 108,200,416 | 106,010,135 | 237 | yes | M | 58 |
| MIC_24 | 41.8266 | -86.4366 | MI | 1999 | MIC-24 | ABRC | 33.8 | 108,176,527 | 105,728,326 | 237 | yes | M | 59 |
| Brn_10 | 41.9 | -86.583 | MI | 2002 | Brn-10 | ABRC | 33.3 | 108,177,381 | 105,905,097 | 243 | yes | M | 7 |
| Brn_24 | 41.9 | -86.583 | MI | 2002 | Brn-24 | ABRC | 38.4 | 108,208,482 | 105,951,803 | 228 | yes | M | 8 |
| Haz_10 | 41.879 | -86.607 | MI | 2002 | Haz-10 | ABRC | 33.8 | 108,154,100 | 105,903,700 | 230 | yes | M | 23 |
| Haz_2 | 41.879 | -86.607 | MI | 2002 | Haz-2 | ABRC | 39.7 | 108,201,103 | 106,004,251 | 288 | yes | M | 24 |
| Ker_4 | 42.184 | -86.358 | MI | 2002 | Ker-4 | ABRC | 32.1 | 108,132,127 | 105,806,486 | 261 | yes | M | 30 |
| Ker_5 | 42.184 | -86.358 | MI | 2002 | Ker-5 | ABRC | 62.9 | 108,259,905 | 106,246,278 | 259 | yes | M | 31 |
| L_R_10 | 41.847 | -86.67 | MI | 2002 | L-R-10 | ABRC | 22.4 | 108,062,944 | 105,496,224 | 186 | yes | M | 49 |
| L_R_5 | 41.847 | -86.67 | MI | 2002 | L-R-5 | ABRC | 60.6 | 108,255,795 | 106,209,826 | 299 | yes | M | 50 |
| Lak_12 | 41.8 | -86.67 | MI | 2002 | Lak-12 | ABRC | 37.8 | 108,176,901 | 105,775,999 | 237 | yes | M | 36 |
| Lak_13 | 41.8 | -86.67 | MI | 2002 | Lak-13 | ABRC | 28.5 | 107,955,559 | 105,553,559 | 226 | yes | M | 37 |
| Map_35 | 42.166 | -86.412 | MI | 2002 | Map-35 | ABRC | 64.7 | 108,265,863 | 106,224,216 | 290 | yes | M | 51 |
| Map_42 | 42.166 | -86.412 | MI | 2002 | Map-42 | ABRC | 46 | 107,303,032 | 106,093,945 | n/a | no | M | 52 |
| Map_8 | 42.166 | -86.412 | MI | 2002 | Map-8 | ABRC | 33.4 | 108,155,999 | 105,921,907 | 287 | yes | M | 53 |
| Mdn_10 | 42.051 | -86.509 | MI | 2002 | Mdn-10 | ABRC | 34.9 | 108,106,772 | 105,906,924 | n/a | no | M | 54 |
| Mdn_8 | 42.051 | -86.509 | MI | 2002 | Mdn-8 | ABRC | 37.4 | 108,199,679 | 105,940,666 | 266 | yes | M | 55 |
| Paw_13 | 42.148 | -86.431 | MI | 2002 | Paw-13 | ABRC | 43 | 108,159,739 | 105,980,721 | 267 | yes | M | 70 |
| Paw_20 | 42.148 | -86.431 | MI | 2002 | Paw-20 | ABRC | 41.3 | 108,218,762 | 106,059,867 | 241 | yes | M | 71 |
| Riv_25 | 42.184 | -86.382 | MI | 2002 | Riv-25 | ABRC | 36.8 | 108,186,632 | 105,779,717 | 273 | yes | M | 76 |
| Riv_26 | 42.184 | -86.382 | MI | 2002 | Riv-26 | ABRC | 35.7 | 108,194,281 | 105,958,738 | 260 | yes | M | 77 |
| Yng_4 | 41.865 | -86.646 | MI | 2002 | Yng-4 | ABRC | 41.3 | 108,182,789 | 106,000,003 | 289 | yes | M | 86 |
| Yng_53 | 41.865 | -86.646 | MI | 2002 | Yng-53 | ABRC | 46 | 108,230,553 | 106,125,861 | 191 | yes | M | 87 |
| RRS_10 | 41.5609 | -86.4251 | IN | 2003 | RRS-10 | ABRC | 41.8 | 108,208,144 | 106,033,465 | 274 | yes | M | 80 |
| DuckLkSP38 | 43.3431 | -86.4045 | MI | 2004 | DuckLkSP38 | ABRC | 37.1 | 108,171,751 | 105,932,415 | 253 | yes | M | 18 |
| DuckLkSP40 | 43.3431 | -86.4045 | MI | 2004 | DuckLkSP40 | ABRC | 39.6 | 108,204,654 | 105,969,244 | 257 | yes | M | 19 |
| KBS_Mac_68 | 42.405 | -85.398 | MI | 2004 | KBS-Mac-68 | ABRC | 41.3 | 108,181,390 | 105,870,424 | 259 | yes | M | 27 |
| KBS_Mac_74 | 42.405 | -85.398 | MI | 2004 | KBS-Mac-74 | ABRC | 37.7 | 108,160,645 | 105,801,702 | 265 | yes | M | 28 |
| MNF_Che_47 | 43.5251 | -86.1843 | MI | 2004 | MNF-Che-47 | ABRC | 27.6 | 108,093,393 | 105,596,885 | 281 | yes | M | 60 |
| MNF_Che_49 | 43.5251 | -86.1843 | MI | 2004 | MNF-Che-49 | ABRC | 28.5 | 108,082,202 | 105,661,610 | 274 | yes | M | 61 |
| MNF_Pin_40 | 43.5356 | -86.1788 | MI | 2004 | MNF-Pin-40 | ABRC | 47.9 | 108,238,775 | 106,099,919 | 287 | yes | M | 62 |
| MNF_Pot_10 | 43.595 | -86.2657 | MI | 2004 | MNF-Pot-10 | ABRC | 61.4 | 108,189,553 | 106,228,588 | n/a | no | M | 63 |
| MNF_Pot_15 | 43.595 | -86.2657 | MI | 2004 | MNF-Pot-15 | ABRC | 25.2 | 108,543,185 | 107,022,924 | n/a | no | M | 64 |
| MSGA_10 | 43.2749 | -86.0891 | MI | 2004 | MSGA-10 | ABRC | 41.9 | 108,191,659 | 106,019,404 | 233 | yes | M | 65 |
| MSGA_12 | 43.2749 | -86.0891 | MI | 2004 | MSGA-12 | ABRC | 42.8 | 108,227,214 | 106,032,928 | 240 | yes | M | 66 |
| MSGA_61 | 43.2749 | -86.0891 | MI | 2004 | MSGA-61 | ABRC | 45.5 | 108,210,152 | 106,077,183 | 247 | yes | M | 67 |
| MuskSP_68 | 43.2483 | -86.3368 | MI | 2004 | MuskSP-68 | ABRC | 25.8 | 108,063,297 | 105,588,467 | 215 | yes | M | 68 |
| MuskSP_83 | 43.2483 | -86.3368 | MI | 2004 | MuskSP-83 | ABRC | 29.9 | 108,099,368 | 105,721,042 | 222 | yes | M | 69 |
| Pent_46 | 43.7623 | -86.3929 | MI | 2004 | Pent-46 | ABRC | 48.3 | 108,227,763 | 106,099,890 | 238 | yes | M | 72 |
| Pent_7 | 43.7623 | -86.3929 | MI | 2004 | Pent-7 | ABRC | 55.7 | 108,220,625 | 106,144,167 | 240 | yes | M | 73 |
| SLSP_67 | 43.665 | -86.496 | MI | 2004 | SLSP-67 | ABRC | 53.5 | 108,238,880 | 106,143,530 | 245 | yes | M | 81 |
| SLSP_69 | 43.665 | -86.496 | MI | 2004 | SLSP-69 | ABRC | 35.5 | 108,160,835 | 105,899,252 | 249 | yes | M | 82 |
| KNO2_41 | 41.273 | -86.625 | IN | 2005 | KNO2.41 | ABRC | 44.7 | 108,209,694 | 106,063,235 | 219 | yes | M | 34 |
| KNO2_54 | 41.273 | -86.625 | IN | 2005 | KNO2.54 | ABRC | 44 | 108,212,430 | 105,903,373 | 218 | yes | M | 35 |
| LI_EF_011 | 40.9064 | -73.1493 | NY | 2005 | LI-EF-011 | ABRC | 68.6 | 108,267,109 | 106,250,331 | 259 | yes | M | 38 |
| LI_EF_018 | 40.9064 | -73.1493 | NY | 2005 | LI-EF-018 | ABRC | 39 | 108,244,306 | 105,898,497 | 230 | yes | M | 39 |
| LI_OF_061 | 40.7777 | -72.9069 | NY | 2005 | LI-OF-061 | ABRC | 58 | 104,897,841 | 105,729,196 | n/a | no | M | 40 |
| LI_RR_096 | 40.9447 | -72.8615 | NY | 2005 | LI-RR-096 | ABRC | 63.5 | 108,264,679 | 106,251,487 | 261 | yes | M | 41 |
| LI_RR_097 | 40.9447 | -72.8615 | NY | 2005 | LI-RR-097 | ABRC | 40.8 | 108,211,310 | 105,992,095 | 249 | yes | M | 42 |
| LI_SET_019 | 40.9352 | -73.114 | NY | 2005 | LI-SET-019 | ABRC | 29.9 | 108,085,297 | 105,737,781 | 259 | yes | M | 43 |
| LI_SET_036 | 40.9352 | -73.114 | NY | 2005 | LI-SET-036 | ABRC | 41.5 | 108,216,592 | 106,006,605 | 238 | yes | M | 44 |
| LI_WP_039 | 40.9076 | -73.2089 | NY | 2005 | LI-WP-039 | ABRC | 104.8 | 108,301,282 | 106,273,259 | 239 | yes | M | 45 |
| LI_WP_041 | 40.9076 | -73.2089 | NY | 2005 | LI-WP-041 | ABRC | 76.5 | 108,287,248 | 106,322,146 | 235 | yes | M | 46 |
| PT1_52 | 41.3423 | -86.7368 | IN | 2005 | PT1.52 | ABRC | 50.6 | 108,240,431 | 106,154,252 | 219 | yes | M | 74 |
| PT1_85 | 41.3423 | -86.7368 | IN | 2005 | PT1.85 | ABRC | 46.1 | 108,220,150 | 106,097,633 | 233 | yes | M | 75 |
| RMX4_118 | 42.036 | -86.511 | MI | 2005 | RMX4.118 | ABRC | 41.8 | 106,178,554 | 105,685,651 | n/a | no | M | 78 |
| 11PNA1_14 | 42.0945 | -86.3253 | MI | 2006 | 11PNA1.14 | ABRC | 47.5 | 108,227,783 | 106,133,372 | 276 | yes | M | 1 |
| 328PNA062 | 42.0945 | -86.3253 | MI | 2006 | 328PNA062 | ABRC | 47.3 | 108,221,709 | 106,127,272 | 223 | yes | M | 2 |
| 627ME_13Y1 | 42.093 | -86.359 | MI | 2006 | n/a | ABRC | 53.4 | 107,908,679 | 106,148,671 | n/a | no | M | 3 |
| 627ME_1MI1 | 42.093 | -86.359 | MI | 2006 | 627ME-1MI1 | ABRC | 57.8 | 108,252,617 | 106,173,403 | 281 | yes | M | 4 |
| 627RMX_1MN4 | 42.0333 | -86.5128 | MI | 2006 | n/a | ABRC | 43.6 | 106,799,549 | 105,789,469 | n/a | no | M | 5 |
| 627RMX_1MN5 | 42.0333 | -86.5128 | MI | 2006 | n/a | ABRC | 50.6 | 106,885,430 | 105,897,441 | n/a | no | M | 6 |
| BRR107 | 40.8313 | -87.735 | IL | 2006 | BRR107 | ABRC | 28.5 | 108,896,513 | 107,320,745 | n/a | no | M | 9 |
| BRR12 | 40.8313 | -87.735 | IL | 2006 | BRR12 | ABRC | 43.9 | 108,190,572 | 106,031,493 | 232 | yes | M | 10 |
| BRR23 | 40.8313 | -87.735 | IL | 2006 | BRR23 | ABRC | 30.7 | 108,095,072 | 105,726,913 | 236 | yes | M | 11 |
| BRR4 | 40.8313 | -87.735 | IL | 2006 | BRR4 | ABRC | 44.7 | 108,180,840 | 106,033,507 | 219 | yes | M | 12 |
| BRR57 | 40.8313 | -87.735 | IL | 2006 | BRR57 | ABRC | 28.4 | 108,093,033 | 105,630,963 | 225 | yes | M | 13 |
| BRR60 | 40.8313 | -87.735 | IL | 2006 | BRR60 | ABRC | 42.9 | 108,281,285 | 106,199,572 | 229 | yes | M | 14 |
| KEN | 41.767 | -72.677 | CT | n/a | KEN | ABRC | 55.2 | 108,233,232 | 106,158,223 | 249 | yes | M | 29 |
| LP3413_31 | 41.6862 | -86.8513 | IN | n/a | LP3413.31 | ABRC | 55.9 | 108,244,332 | 106,190,596 | 227 | yes | M | 47 |
| LP3413_53 | 41.6862 | -86.8513 | IN | n/a | LP3413.53 | ABRC | 51.2 | 108,157,453 | 105,994,665 | 245 | yes | M | 48 |
| RMX413_85 | 42.036 | -86.511 | MI | n/a | RMX413.85 | ABRC | 38 | 106,816,221 | 105,483,632 | n/a | no | M | 79 |

**Table S2.**
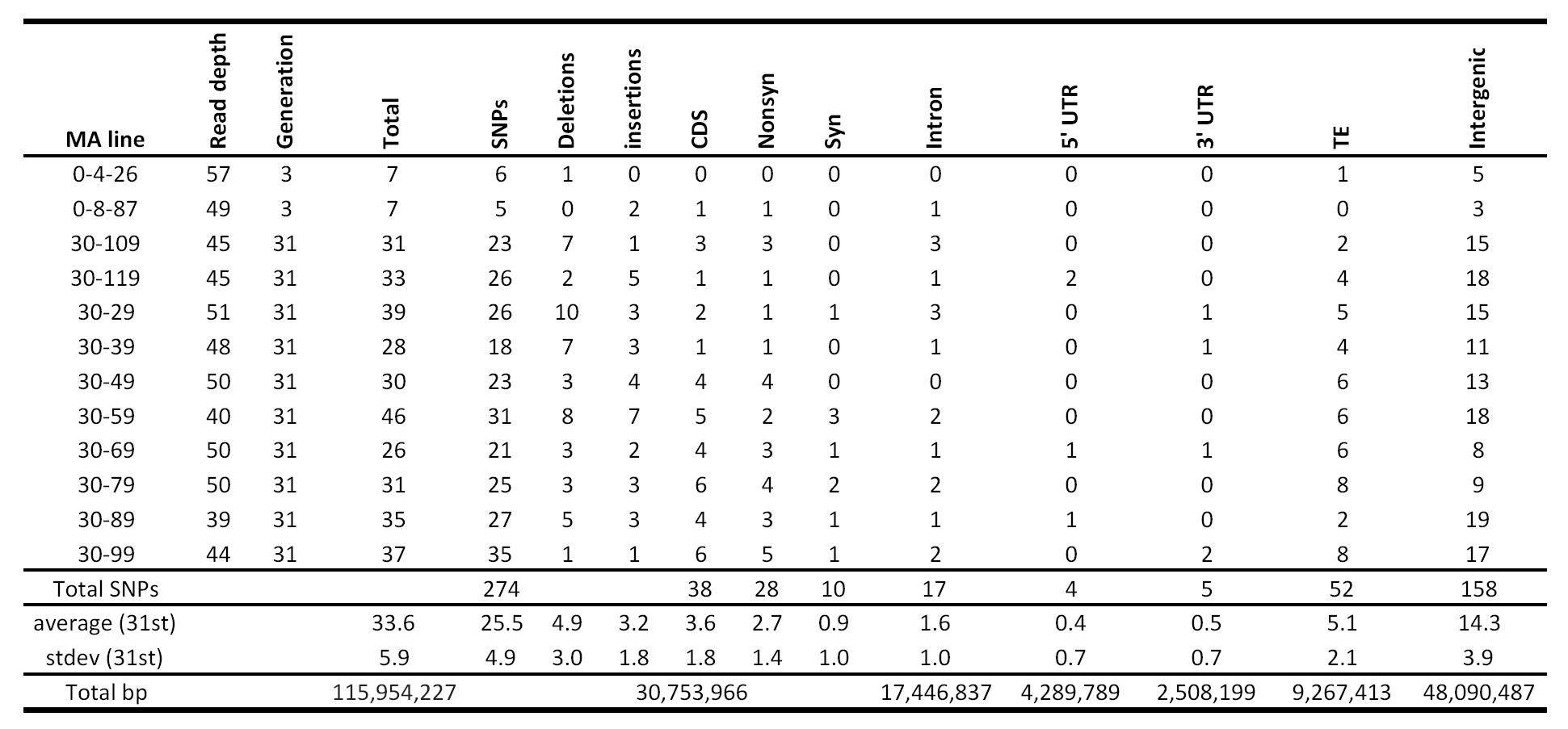
Sample information for Col-0 mutation accumulation lines. Information about each Mutation Accumulation (MA) line and their number of SNPs at different annotations. Also the total number of SNPs, average number of mutations and total bp covered in the genome per annotation are reported.

**Table S3.** Mutation rate estimates for different annotations in HPG1 and mutation accumulation lines. Mutation rates from MA lines are compared to HPG1 substitution rates from the dataset of 32_15 quality filter and complete information (see SOM) (Abbreviations: stat, descriptive statistic; bp, base pairs; lower and upper, lower and upper 95% CI; Nonsyn. and Syn., nonsynonymous and synonymous sites; UTR, untranslated region sites; HPG1 adj., substitution rate of HPG1 adjusted by a mean generation time of 1.3 years)

| Dataset | stat | CDS | Syn. | Nonsyn. | Intronic | 5' UTR | 3' UTR | Transposon | Intergenic | Genome |
| --- | --- | --- | --- | --- | --- | --- | --- | --- | --- | --- |
| MA | mean | 3.776 | n/a | n/a | 2.958 | 3.008 | 6.431 | 17.752 | 9.592 | 7.094 |
| MA | sem | 1.928 | n/a | n/a | 1.786 | 5.258 | 9.094 | 7.420 | 2.628 | 1.352 |
| MA | lower | 2.581 | n/a | n/a | 1.851 | -0.251 | 0.794 | 13.153 | 7.964 | 6.256 |
| MA | upper | 4.971 | n/a | n/a | 4.065 | 6.267 | 12.067 | 22.351 | 11.221 | 7.932 |
| HPG1 | mean | 2.149 | n/a | n/a | 1.540 | n/a | n/a | 2.290 | 3.029 | 2.114 |
| HPG1 | sem | 0.108 | n/a | n/a | 0.165 | n/a | n/a | 0.536 | 0.173 | 0.119 |
| HPG1 | lower | 1.943 | n/a | n/a | 1.231 | n/a | n/a | 1.314 | 2.698 | 1.871 |
| HPG1 | upper | 2.364 | n/a | n/a | 1.874 | n/a | n/a | 3.309 | 3.368 | 2.344 |
| HPG1 adj. | mean | 2.794 | n/a | n/a | 2.002 | n/a | n/a | 2.977 | 3.938 | 2.748 |
| HPG1 adj. | sem | 0.140 | n/a | n/a | 0.214 | n/a | n/a | 0.697 | 0.225 | 0.154 |
| HPG1 adj. | lower | 2.526 | n/a | n/a | 1.600 | n/a | n/a | 1.708 | 3.508 | 2.432 |
| HPG1 adj. | upper | 3.073 | n/a | n/a | 2.436 | n/a | n/a | 4.302 | 4.378 | 3.047 |
| Distribution of pairwise SNP differences | min | 0 | 0 | 0 | 0 | 0 | 0 | 0 | 0 | 0 |
|  | 1st qu. | 2 | 1 | 1 | 1 | 0 | 1 | 2 | 5 | 9 |
|  | median | 5 | 3 | 3 | 3 | 1 | 2 | 4 | 10 | 18 |
|  | mean | 5.6 | 3 | 3.1 | 3.8 | 1.2 | 1.9 | 4.3 | 11.3 | 21.1 |
|  | 3rd qu. | 8 | 5 | 4 | 5 | 2 | 3 | 6 | 16 | 31 |
|  | max. | 27 | 17 | 11 | 15 | 5 | 7 | 22 | 43 | 87 |
| Total number of SNPs |  | 971 | 531 | 448 | 629 | 74 | 158 | 656 | 2498 | 5013 |
| Total bp |  | 32119233 | n/a | n/a | 18132262 | 2632130 | 4480510 | 6209512 | 43601507 | 108434034 |

**Table S4.** Description of phenotypic and climatic variables for association mapping analyses. Mean and standard deviation (s.d.) across accessions for each phenotypic and climatic variables. Broad sense heritabilities (H2) were calculated from between line and within line (between replicate) variance in ANOVA. P-value corresponds to F test. Narrow sense heritabilities (h2) were calculated employing linear mixed models and kinship matrix from mean accession values. P-values correspond to Likelihood Ratio test.

| Variable | Description | mean | s.d. | H2 | p-value | h2 | p-value |
| --- | --- | --- | --- | --- | --- | --- | --- |
| FT_V0 | Time from germination until the first flower opens (days) under 0 days of vernalization | 101 | 4.53 | 0.009 | 7.28E-03 | 0.017 | 1.97E-25 |
| FT_V1 | Time from germination until the first flower opens (days) under 14 days of vernalization | 107 | 4.12 | 0.013 | 6.87E-04 | 0.395 | 1.83E-25 |
| FT_V2 | Time from germination until the first flower opens (days) under 28 days of vernalization | 102 | 3.22 | 0.012 | 1.04E-03 | 0.429 | 3.37E-27 |
| FT_V3 | Time from germination until the first flower opens (days) under 63 days of vernalization | 110 | 1.32 | 0.010 | 5.11E-03 | 0.226 | 9.52E-25 |
| B_V0 | Time from germination until the first developed bud (days) under 0 days of vernalization | 88.8 | 4 | 0.013 | 8.99E-04 | 0.018 | 2.26E-25 |
| B_V1 | Time from germination until the first developed bud (days) under 14 days of vernalization | 93.9 | 3.84 | 0.009 | 7.45E-03 | 0.340 | 3.98E-25 |
| B_V2 | Time from germination until the first developed bud (days) under 28 days of vernalization | 89.2 | 2.13 | 0.005 | 6.92E-02 | 0.252 | 2.22E-25 |
| B_V3 | Time from germination until the first developed bud (days) under 63 days of vernalization | 101 | 0.45 | 0.006 | 5.79E-02 | 0.177 | 1.99E-24 |
| Fecundity | Pixel area of inflorescence (correlation with number of fruits, rho=0.84) | 0.02 | 0.0042 | 0.001 | 3.56E-01 | 0.240 | 1.02E-22 |
| seed_size | Average seed size (mm2) | 0.134 | 0.0053 | 0.016 | 4.73E-03 | 0.149 | 3.58E-24 |
| GR_rootLength | Average root growth rate | 181 | 14.9 | 0.131 | 4.76E-77 | 0.640 | 3.13E-29 |
| GR_shootArea | Average of shoot area growth rate | 2279 | 253 | 0.053 | 2.33E-24 | 0.812 | 1.77E-31 |
| rootLength | Average root length | 467 | 35.8 | 0.048 | 2.01E-21 | 0.409 | 2.57E-28 |
| dirEquivalent | Average root direction index. Score for average pixel-by-pixel deviations from growth relative to vector of gravity | 0.393 | 0.0277 | 0.059 | 2.62E-28 | 0.544 | 1.14E-26 |
| stdDevXY | Average root linearity coefficient of linear determination; R2 of linear regression line fitted to pixels of primary root skeleton | 0.725 | 0.0429 | 0.018 | 4.54E-06 | 0.303 | 1.41E-25 |
| meanRootWidth | Average root width | 5.27 | 0.177 | 0.038 | 5.30E-16 | 0.359 | 1.52E-25 |
| rootWidth20 | Average width over first interval of the primary root length (0 to 20%) at hypocotyl/root junction | 5.75 | 0.124 | 0.018 | 5.11E-06 | 0.166 | 3.37E-25 |
| rootWidth40 | Average width over first interval of the primary root length (20 to 40%) at hypocotyl/root junction | 5.35 | 0.19 | 0.033 | 3.87E-13 | 0.291 | 1.76E-25 |
| rootWidth60 | Average width over first interval of the primary root length (40 to 60%) at hypocotyl/root junction | 5.2 | 0.212 | 0.039 | 1.49E-16 | 0.405 | 6.51E-26 |
| rootWidth80 | Average width over first interval of the primary root length (60 to 80%) at hypocotyl/root junction | 5.11 | 0.241 | 0.045 | 4.67E-20 | 0.381 | 5.47E-26 |
| rootWidth100 | Average width over first interval of the primary root length (80 to 100%) at hypocotyl/root junction | 4.9 | 0.222 | 0.038 | 4.06E-16 | 0.351 | 8.81E-26 |
| gravitropicDir | Average root angle between root vector and the vertical axis of the picture (assumed vector of gravity) (°) | -7.22 | 2.56 | 0.024 | 7.69E-09 | 0.210 | 4.68E-27 |
| gravitropicScore | Average score for root angle intervals | 0.1 | 0.0457 | 0.044 | 2.83E-19 | 0.642 | 7.56E-27 |
| TotLen.EucLen | Average root tortuosity: Total root length divided by Euclidian length | 1.1 | 0.0097 | 0.009 | 6.83E-03 | 0.422 | 2.53E-25 |
| GR.TL | Average relative root growth rate: Root growth rate divided by total length at the earlier time point | 0.673 | 0.0796 | 0.011 | 1.20E-03 | 0.393 | 2.69E-24 |
| BIO1 | Annual Mean Temperature (°C x 10) | 98.1 | 12.8 | n/a | n/a | 0.066 | 3.22E-40 |
| BIO2 | Mean Diurnal Range (Mean of monthly (max temp - min temp)) | 107 | 7.65 | n/a | n/a | 0.073 | 1.02E-40 |
| BIO3 | Isothermality (BIO2/BIO7) (x 100) | 28.9 | 1.8 | n/a | n/a | 0.361 | 4.91E-39 |
| BIO4 | Temperature Seasonality (standard deviation x 100) | 9169 | 483 | n/a | n/a | 0.383 | 4.68E-47 |
| BIO5 | Max Temperature of Warmest Month (°C x 10) | 283 | 10.1 | n/a | n/a | 0.152 | 3.78E-40 |
| BIO6 | Min Temperature of Coldest Month (°C x 10) | -80.9 | 18 | n/a | n/a | 0.275 | 4.79E-42 |
| BIO7 | Temperature Annual Range (BIO5-BIO6) (°C x 10) | 364 | 17.5 | n/a | n/a | 0.239 | 6.31E-42 |
| BIO8 | Mean Temperature of Wettest Quarter (°C x 10) | 176 | 55.1 | n/a | n/a | 0.016 | 3.58E-43 |
| BIO9 | Mean Temperature of Driest Quarter (°C x 10) | -7.11 | 48.7 | n/a | n/a | 0.000 | 3.58E-43 |
| BIO10 | Mean Temperature of Warmest Quarter (°C x 10) | 213 | 10.8 | n/a | n/a | 0.205 | 3.33E-40 |
| BIO11 | Mean Temperature of Coldest Quarter (°C x 10) | -24.1 | 18.2 | n/a | n/a | 0.270 | 1.71E-41 |
| BIO12 | Annual Precipitation (mm) | 990 | 109 | n/a | n/a | 0.219 | 3.94E-44 |
| BIO13 | Precipitation of Wettest Month (mm) | 103 | 6.72 | n/a | n/a | 0.206 | 1.53E-40 |
| BIO14 | Precipitation of Driest Month (mm) | 54.1 | 16.7 | n/a | n/a | 0.104 | 1.51E-40 |
| BIO15 | Precipitation Seasonality (Coefficient of Variation) | 17.8 | 5.51 | n/a | n/a | 0.157 | 8.93E-40 |
| BIO16 | Precipitation of Wettest Quarter (mm) | 291 | 19.7 | n/a | n/a | 0.269 | 1.55E-42 |
| BIO17 | Precipitation of Driest Quarter (mm) | 191 | 44.8 | n/a | n/a | 0.084 | 3.67E-42 |
| BIO18 | Precipitation of Warmest Quarter (mm) | 277 | 25.2 | n/a | n/a | 0.342 | 7.42E-44 |
| BIO19 | Precipitation of Coldest Quarter (mm) | 197 | 47 | n/a | n/a | 0.022 | 2.68E-42 |

**Table S5.**
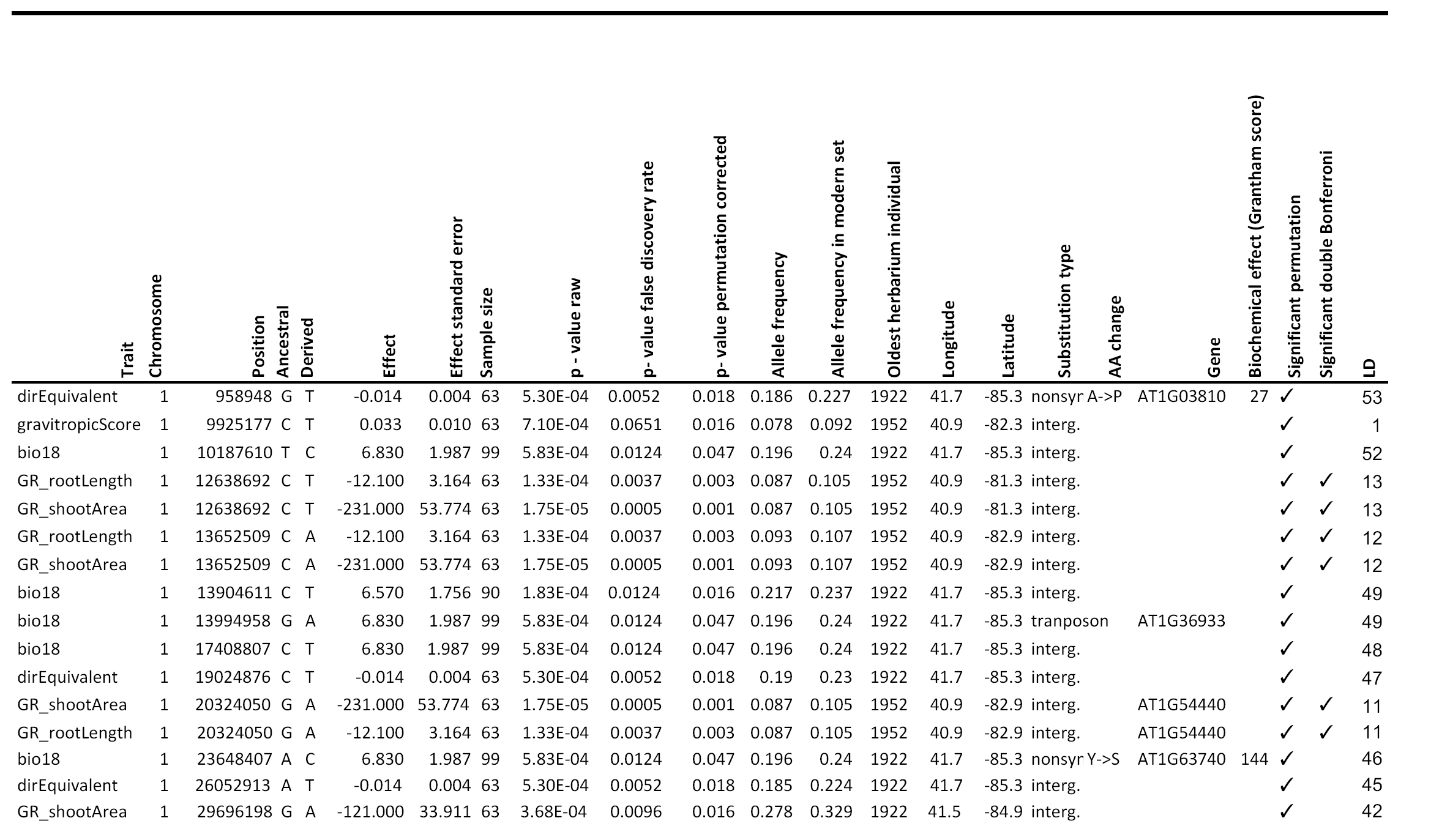
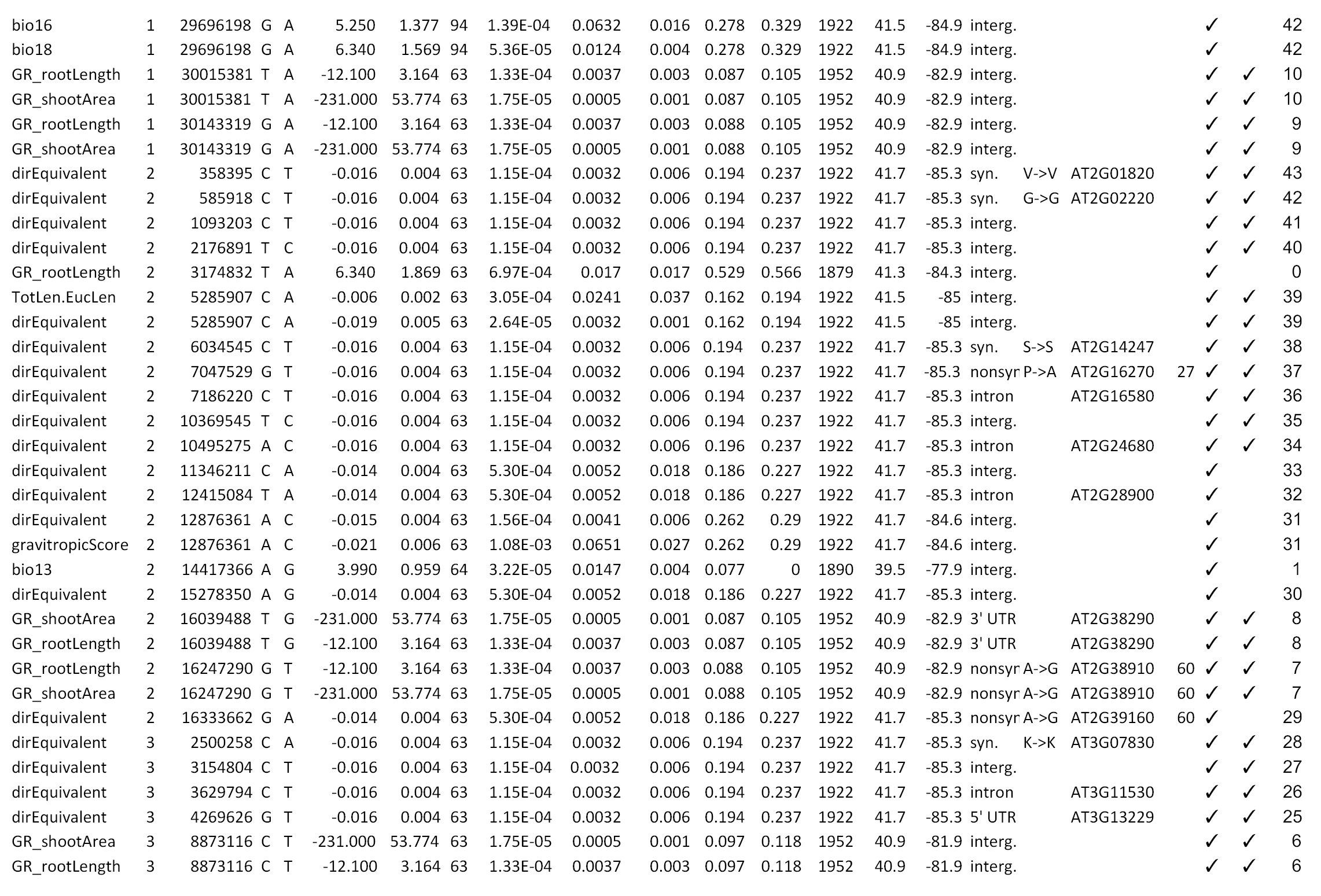
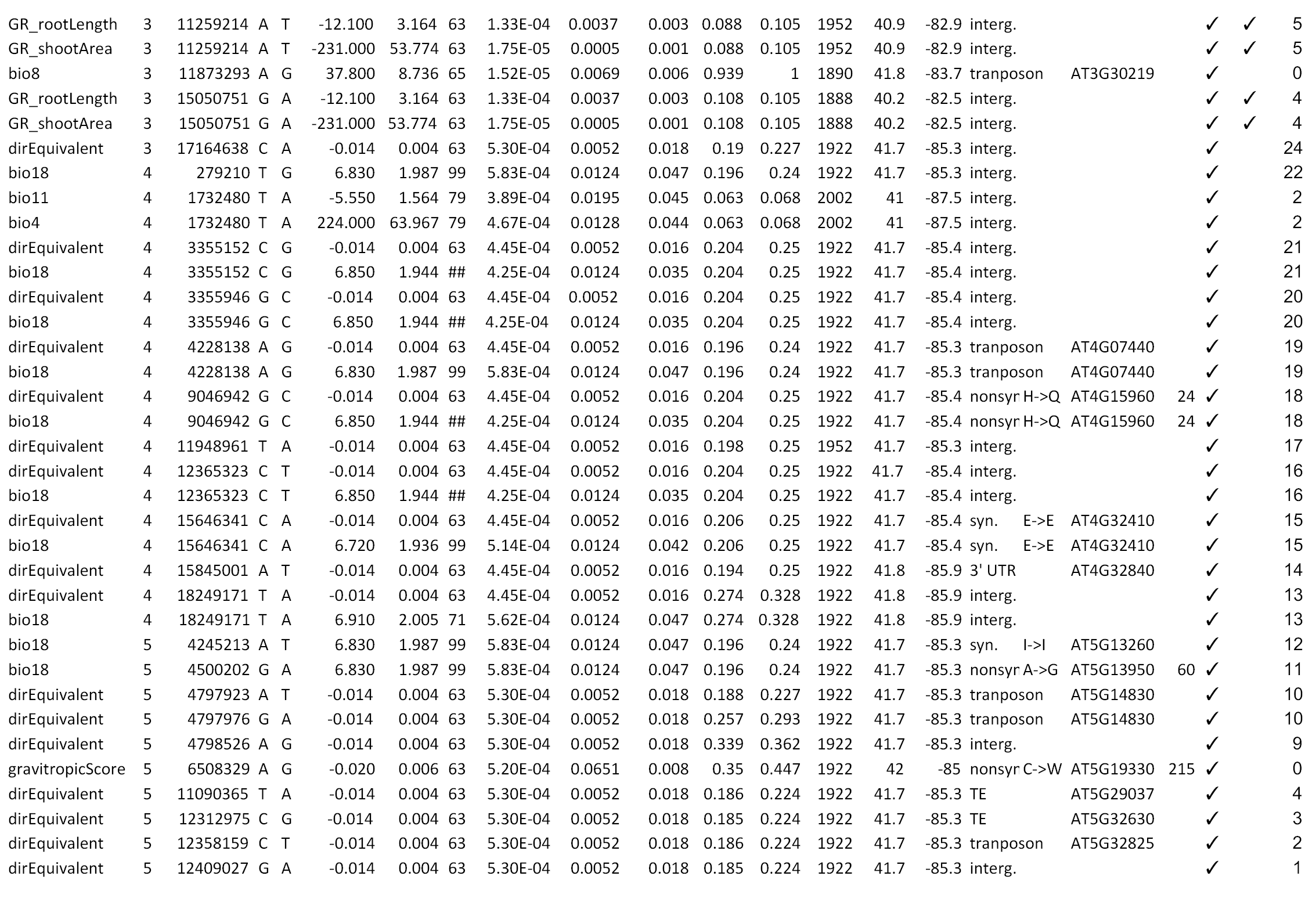
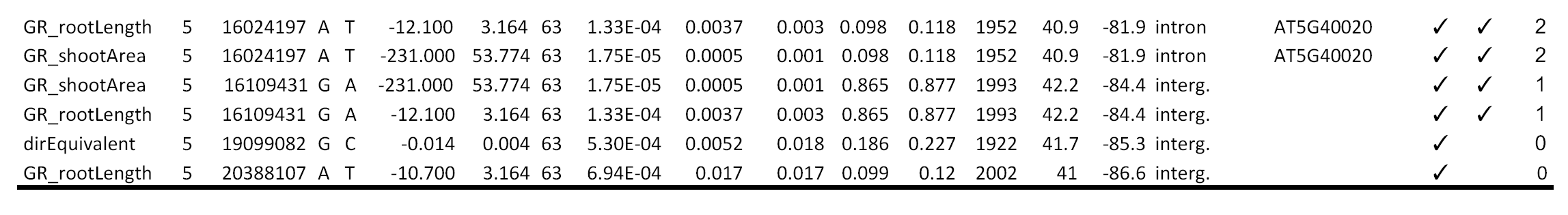
SNP hits from association analyses and several descriptors. SNP hits significant at the 5% level after permutation correction are shown. Additionally, if raw p-values pass a double Bonferroni threshold of 0.01% are marked with a “ tick”. (Abbreviations: nonsyn. and syn., nonsynonymous and synonymous changes; regular one-letter abbreviation was used for amino acid changes)

